# Scalable methods for analyzing and visualizing phylogenetic placement of metagenomic samples

**DOI:** 10.1101/346353

**Authors:** Lucas Czech, Alexandros Stamatakis

## Abstract

The exponential decrease in molecular sequencing cost generates unprecedented amounts of data. Hence, scalable methods to analyze these data are required. Phylogenetic (or Evolutionary) Placement methods identify the evolutionary provenance of anonymous sequences with respect to a given reference phylogeny. This increasingly popular method is deployed for scrutinizing metagenomic samples from environments such as water, soil, or the human gut.

Here, we present novel and, more importantly, highly scalable methods for analyzing phylogenetic placements of metagenomic samples. More specifically, we introduce methods for (a) visualizing differences between samples and their correlation with associated meta-data on the reference phylogeny, (b) clustering similar samples using a variant of the fc-means method, and (c) finding phylogenetic factors using an adaptation of the Phylofactorization method. These methods enable to interpret metagenomic data in a phylogenetic context, to find patterns in the data, and to identify branches of the phylogeny that are driving these patterns.

To demonstrate the scalability and utility of our methods, as well as to provide exemplary interpretations of our methods, we applied them to 3 publicly available datasets comprising 9782 samples with a total of approximately 168 million sequences. The results indicate that new biological insights can be attained via our methods.

## 2 Introduction

The availability of high-throughput DNA sequencing technologies has revolutionized biology by transforming it into an ever more data-driven and compute-intense discipline [1]. In particular, Next Generation Sequencing (NGS) [2], as well as later generations [3–6], have given rise to novel methods for studying microbial environments [7–10]. These technologies are often used in metagenomic studies to sequence organisms in water [11–13] or soil [14,15] samples, in the human microbiome [16–18], and a plethora of other environments. These studies yield a large set of short anonymous DNA sequences, so-called reads, for each sample. Reads that are obtained from specific parts of the genome are called meta-barcoding reads; most often, reads are amplified before sequencing and later de-replicated again, resulting in so-called amplicons. A typical task in metagenomic studies is to identify and classify these sequences with respect to known reference sequences, either in a taxonomic or a phylogenetic context.

Conventional methods like BLAST [19] are based on sequence similarity or identity. Such methods are fast, but only attain satisfying accuracy levels if the query sequences (e.g., the environmental reads or amplicons) are sufficiently similar to the reference sequences. Furthermore, BLAST might yield suboptimal results [20], and the best BLAST hit does often *not* represent the most closely related species [21].

Alternatively, so-called phylogenetic (or evolutionary) placement methods [22–24] identify query sequences based on a phylogenetic tree of reference sequences. Thereby, they incorporate information about the evolutionary history of the species under study and hence provide a more accurate means for read identification. We outline the method in more detail below, and describe the standard data analysis pipeline in S2 Text. The result of a phylogenetic placement is a mapping of the query sequences to the branches of the reference tree. Such a mapping also elucidates the evolutionary distance between the query and the reference sequences.

The information provided by phylogenetic placement represents useful biological knowledge *per se*, which has already been used for developing several downstream analysis methods. For example, the placement of query sequences on the branches of the reference tree can be understood as a classification of the query sequences with respect to the given phylogeny. This classification can then be summarized by means of sequence abundances [25,26], or be used to obtain taxonomic annotations [27,28]. Phylogenetic placement can also be utilized to compare sets of environmental samples with each other (for example, from different locations or points in time). Note that distinct samples from one study are typically mapped to the same underlying reference tree, thus facilitating such comparisons. For instance, existing analysis methods such as Edge PCA and Squash Clustering [29] exploit the information provided by the reference tree to visualize differences between sets of metagenomic samples, or to cluster samples based on placement similarity. We show examples of Edge PCA and Squash Clustering results later in Fig 8.

In this paper, we present novel, scalable methods to analyze and visualize phylogenetic placement data. The remainder of this article is structured as follows. First, we introduce phylogenetic placement and related terminology, and provide an overview over existing post-analysis methods for phylogenetic placements. Then, we describe several novel methods for visualizing differences between the placement data of distinct environmental samples, and for visualizing their correlation with per-sample meta-data. Next, we propose a clustering algorithm for samples that is useful for analyzing extremely large environmental studies. Furthermore, we present an adaptation of the Phylogenetic Isometric Log-Ratio (PhILR) transformation and balances [30] to phylogenetic placement data. Lastly, we introduce an adaptation of Phylofactorization [31] to placements. Phylofactorization is a method that identifies those edges in the phylogeny that characterize patterns in the data related to per-sample meta-data. In order to evaluate our methods, we apply them to three real word datasets, namely, Bacterial Vaginosis (BV) [18], Tara Oceans (TO) [11,32,33], and Human Microbiome Project (HMP) [16,17], which are introduced in more detail later. We provide exemplary interpretations of the results obtained from these datasets, compare the results to existing methods, and analyze the run-time performance of our methods.

Our methods are implemented in our gappa tool, which is freely available under GPLv3 at http://github.com/lczech/gappa. We provide an overview of the tool in S2 Text. Furthermore, scripts, data, and other tools used for the tests and generating the figures presented here are available at http://github.com/lczech/placement-methods-paper.

### 2.1 Phylogenetic placement

In brief, phylogenetic placement calculates the most probable insertion branches for each given query sequence (QS) on a reference tree (RT). The QSs are reads or amplicons from environmental samples. The RT and the reference sequences it represents are typically assembled by the user so that they capture the expected species diversity in the samples. Nonetheless, we recently presented an automated approach for selecting and constructing appropriate reference sequences from large sequence databases [34]. Current implementations of phylogenetic placement furthermore expect the RT to be strictly bifurcating. Prior to the placement, the QSs need to be aligned against the reference alignment of the RT by programs such as P_A_P_A_R_A_ [35,36] or HMMALIGN, which is part of the HMMER suite [37,38]. The placement is then conducted by initially inserting one QS as a new tip into a branch of the tree, then re-optimizing the branch lengths that are most affected by this insertion, and thereafter evaluating the resulting likelihood score of the tree under a given model of nucleotide substitution, such as the Generalized Time-Reversible (GTR) model [39]. The QS is then removed from the current branch and subsequently placed into all other branches of the RT.

Thus, for each branch of the tree, the process yields a so-called *placement* of the QS, that is, an optimized position on the branch, along with a likelihood score for the whole tree. The likelihood scores for a QS are then transformed into probabilities, which quantify the uncertainty of placing the sequence on the respective branch [40, 41]. Those probabilities are called likelihood weight ratios (LWRs). The accumulated LWR sum over all branches for a single QS is 1.0. On the one hand, sequences that have one or a few closely related reference sequences in the reference tree thus exhibit high LWRs at the respective branches, while having LWRs close to 0 on most other branches. On the other hand, if suitable references are missing from the tree, or if the chosen genetic region of the sequences has a poor phylogenetic resolution with respect to the query sequences, the LWRs can be distributed across several parts of the tree, indicating a high degree of placement uncertainty. Fig 1 shows an example depicting the placements of one QS, including the respective LWRs.

**Fig 1.**
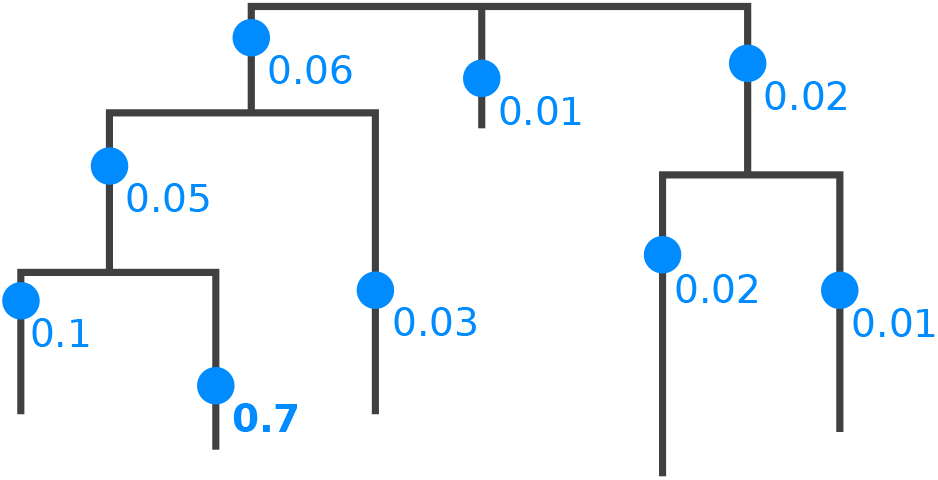
Phylogenetic Placement of a single Query Sequence. Each branch of the reference tree is tested as a potential insertion position, called a “placement” (blue dots). Note that placements have a specific position on their branch, due to the branch length optimization process. A probability of how likely it is that the sequence belongs to a specific branch is computed (numbers next to dots), which is called the *likelihood weight ratio* (LWR). The bold number (0.7) denotes the most probable placement of the sequence.

This process is repeated for every QS. Note that the placement process is conducted *independently* for each QS, and that the phylogenetic relationships among the QSs are not resolved. That is, for each QS, the algorithm starts calculating placements from scratch on the original RT.

In summary, the result of a phylogenetic placement analysis is a mapping of the QSs in a sample to positions on the branches of the RT. Each such position, along with the corresponding LWR, is called a placement of the QS.

### 2.2 Normalization

When placing multiple samples, for instance, from different geographical locations, typically, the same RT is used, to allow for comparing the phylogenetic composition of these samples. Then, each sample results in a set of placements on the RT for all QSs contained in the sample. If one intends to compare different samples based on the placement of their sequences, it is important to consider how to properly normalize the placements per sample. Normalization is required as the sample size (often also called library size), that is, the number of sequences per sample, can vary by several orders of magnitude, due to efficiency variations in the sequencing process or biases introduced by the amplification process. As a consequence, metagenomic sequence data are inherently compositional [42], which can lead to spurious statistical analyses [43–46]. This impedes statistical analyses of the data and hence needs to be considered in all analysis steps [30].

We here briefly outline common types of problems due to normalization that also affect phylogenetic placement data. Selecting an appropriate normalization strategy constitutes a common general problem in metagenomic studies. The appropriateness depends on data characteristics [47], but also on the biological question. For example, estimating indices such as the species richness are often implemented via rarefaction and rarefaction curves [48], which potentially omit a large amount of the available valid data [49]. Furthermore, the specific type of input sequence data has to be taken into account for normalization: Biases induced by the amplification process can potentially be avoided if, instead of amplicons, data based on shotgun sequencing are used, such as _mi_tags [50]. Moreover, the sequences can be clustered prior to a phylogenetic placement analysis, for instance, by constructing operational taxonomic units (OTUs) [51–54]. Analyses using OTUs focus on species diversity instead of simple abundances. OTU clustering substantially reduces the number of sequences, and hence greatly decreases the computational cost for placement analyses. Lastly, one may completely ignore the abundances (which are called the “multiplicities” of placements) of the placed sequences, reads, or OTUs, and only be interested in their presence/absence when comparing samples.

All of the above analysis strategies are also applicable to phylogenetic placement, for instance, by placing OTUs on the RT instead of sequences. Which of these strategies is deployed, depends on the specific design of the study and the research question at hand. The common challenge is that the number of sequences per sample differs, which affects most post-analysis methods, and can lead to conflicting interpretations and irreproducible results [45,55]. For example, if inappropriately normalized, the visualization methods presented here might highlight irrelevant branches or clades of the tree.

Before introducing our methods, we therefore first explain how the necessary normalizations of sample sizes can be performed. We also describe general techniques for interpreting and working with phylogenetic placement data. These are not methods of their own, but tools necessary for later. Some of these techniques have been used before as building blocks for methods such as Edge PCA and Squash Clustering [29,56].

### 2.3 Edge masses

Methods that compare samples directly based on their sequences, such as the UniFrac distance [57,58], can benefit from rarefaction [47]. However, in the context of phylogenetic placement, rarefaction is not necessary. Thus, a larger amount of valid data can be kept. For a thorough comparison of UniFrac to methods based on phylogenetic placement, see [29], which furthermore applies Edge PCA and Squash Clustering on the BV dataset that we later user for evaluation.

In order to compare the placements obtained from a set of samples, it is convenient to think of the reference tree as a graph (when exploiting graph properties of the tree, we refer to the branches of the tree as edges). Then, the per-branch LWRs for a single QS can be interpreted as a distribution of mass points over the edges of the RT, including their respective placement positions on the branches, cf. Fig 1. This implies that each QS has a total accumulated mass of 1.0 on the RT. We call this the *mass interpretation* of the placed QSs, and henceforth use mass and LWR interchangeably. The *mass of an edge* or *edge mass* refers to the sum of the LWRs on that edge for all QSs of a sample, as shown in Fig 2(a). The edge masses can serve as a simplification of the data that summarizes a whole sample in a vector with *m* entries, for an RT with *m* edges. That is, instead of describing a sample by the placements of all its sequences on the RT, we describe it by the total per-branch mass of all its sequences. A larger example for the full RT is shown in Fig 3(a). The total mass of a sample is then the sum over all edge masses, which is identical to the number of QSs in the sample.

**Fig 2.**
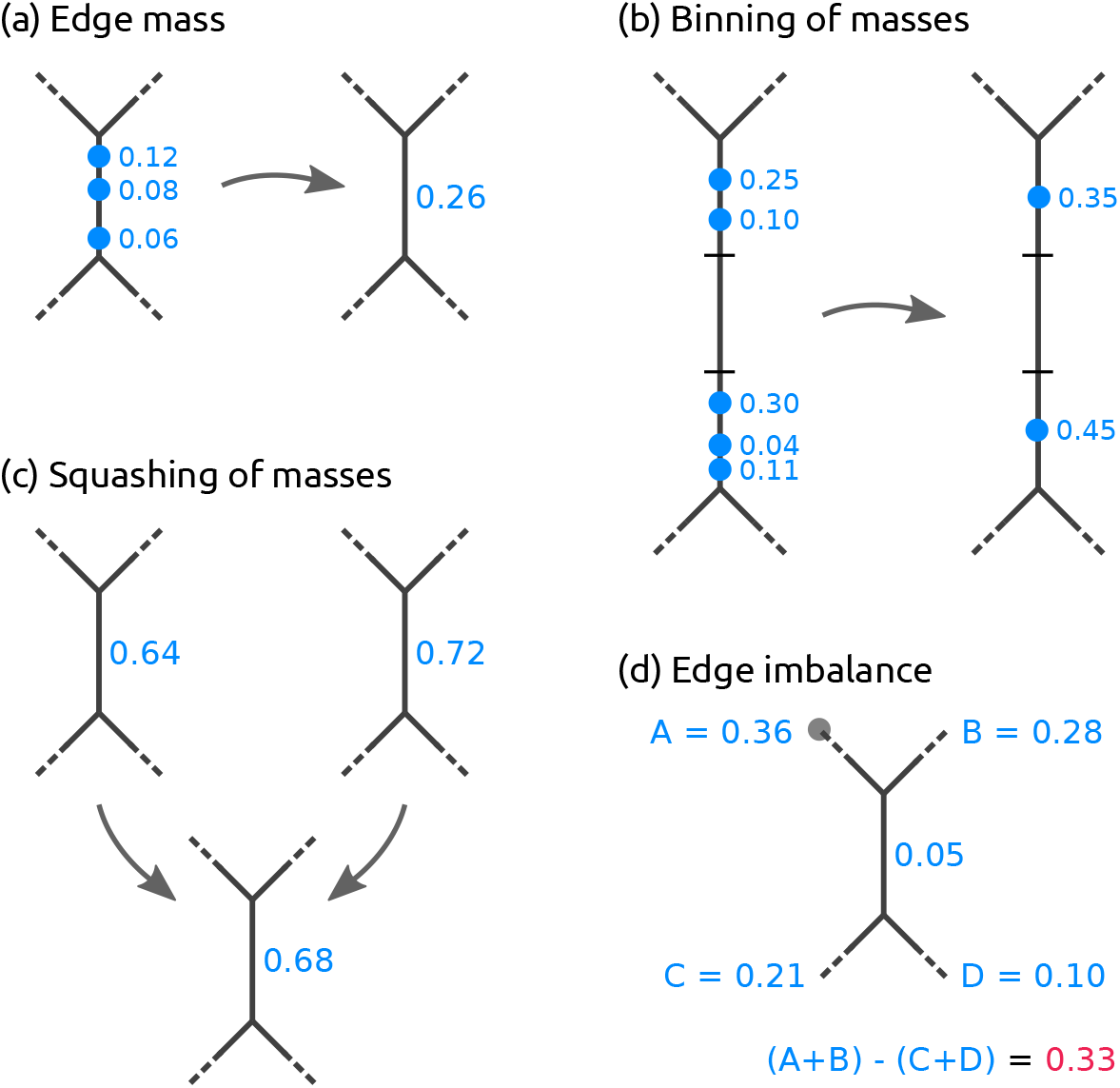
Operations on Placement Masses. (a) The *edge mass* or *mass of an edge* is the sum of likelihood weight ratios (LWRs) on that edge for all query sequences (QSs) in a sample. Here, three placements from three QSs on the branch are summarized. (b) In order to reduce time and memory of the computations, masses can be *binned* by summarizing them across QSs in intervals along the edges. (c) The masses on corresponding edges of the RT of two or more samples can be *squashed* to represent the average mass distribution of the samples. For simplicity, we here use equal weights, and show edge masses instead of individual LWRs. (d) The *edge imbalance* of an edge is the sum of masses on all edges on the root side of the edge (`A+B`, with the root in subtree A denoted as a gray dot) minus the sum of the masses on the edges on the non-root side (`C+D`), while ignoring the mass on the edge itself.

The key idea is to use the distribution of placement mass points over the edges of the RT to characterize a sample. This allows for normalizing samples of different size by scaling the total sample mass to unit mass 1.0. This is done by dividing the mass of each placement location by the total sum of all masses in the sample. In other words, absolute abundances are converted into relative abundances. This way, rare species, which might have been removed by rarefaction, can be kept, as they only contribute a negligible mass to the branches into which they have been placed. This approach is analogous to using proportional values for methods based on OTU count tables, that is, scaling each sample/column of the table by its sum of OTU counts [47]. Most of the methods presented here use normalized samples, that is, they use relative abundances.

As relative abundances are compositional data, certain caveats occur [43, 55, 59], which we discuss where appropriate.

When working with large numbers of QSs, the mass interpretation allows to further simplify and reduce the data: The masses on each edge of the tree can be quantized into *b* discrete bins, as shown in Fig 2(b). That is, each edge is divided into *b* intervals (or bins) of the corresponding branch length, and all mass points on that edge are accumulated into their respective nearest bin. The parameter *b* controls the resolution and accuracy of this approximation. In the extreme case *b* := 1, all masses on an edge are grouped into one single bin (which is equivalent to only considering the edge mass as described above instead of individual LWRs). This *branch binning* process drastically reduces the number of mass points that need to be stored and analyzed in several of the methods we present, while only inducing a negligible decrease in accuracy. As shown in S2 Table, branch binning can yield a speedup of up to 75% for post-analysis run-times without altering the results of the analysis.

The interpretation of placements as masses on the edges of the tree further allows to summarize a set of samples by annotating the RT with their (weighted) average per-edge mass distribution, as shown in Fig 2(c). This procedure is called *squashing* [29]. Because phylogenetic placements represent point masses on the edges of the tree, squashing can be seen as joining the (weighted) set of placements of the samples on corresponding edges of the RT. The resulting masses can then be normalized again to obtain unit mass for the resulting average tree. This tree summarizes the (sub-)set of samples it represents; a larger example of squashing is shown in Fig 5.

### 2.4 Edge imbalances

So far, we have only considered the per-edge masses. Often, however, it is also of interest to “summarize” the mass of an entire clade. For example, sequences of the RT that represent species or strains might not provide sufficient phylogenetic signal for properly resolving the phylogenetic placement of short sequences [60]. In these cases, the placement mass of a sequence can be spread across different edges representing the same genus or species, thus blurring analyses based on per-edge masses.

Instead, a clade-based summary can yield clearer analysis results. It can be computed by using the tree structure to appropriately transform the edge masses. Each edge splits the tree into two parts, of which only one contains the root (or top-level trifurcation) of the tree. For a given edge, its mass difference is then calculated by summing all masses in the root part of the tree and subtracting all masses in the other part, while ignoring the mass of the edge itself [29], as shown in Fig 2(d). This difference is called the *imbalance* of the edge [29]. It is usually normalized to represent unit total mass, as the absolute (not normalized) imbalance otherwise propagates the effects of differing sample sizes all across the tree. A larger example for the full RT is shown in Fig 3(a).

**Fig 3.**
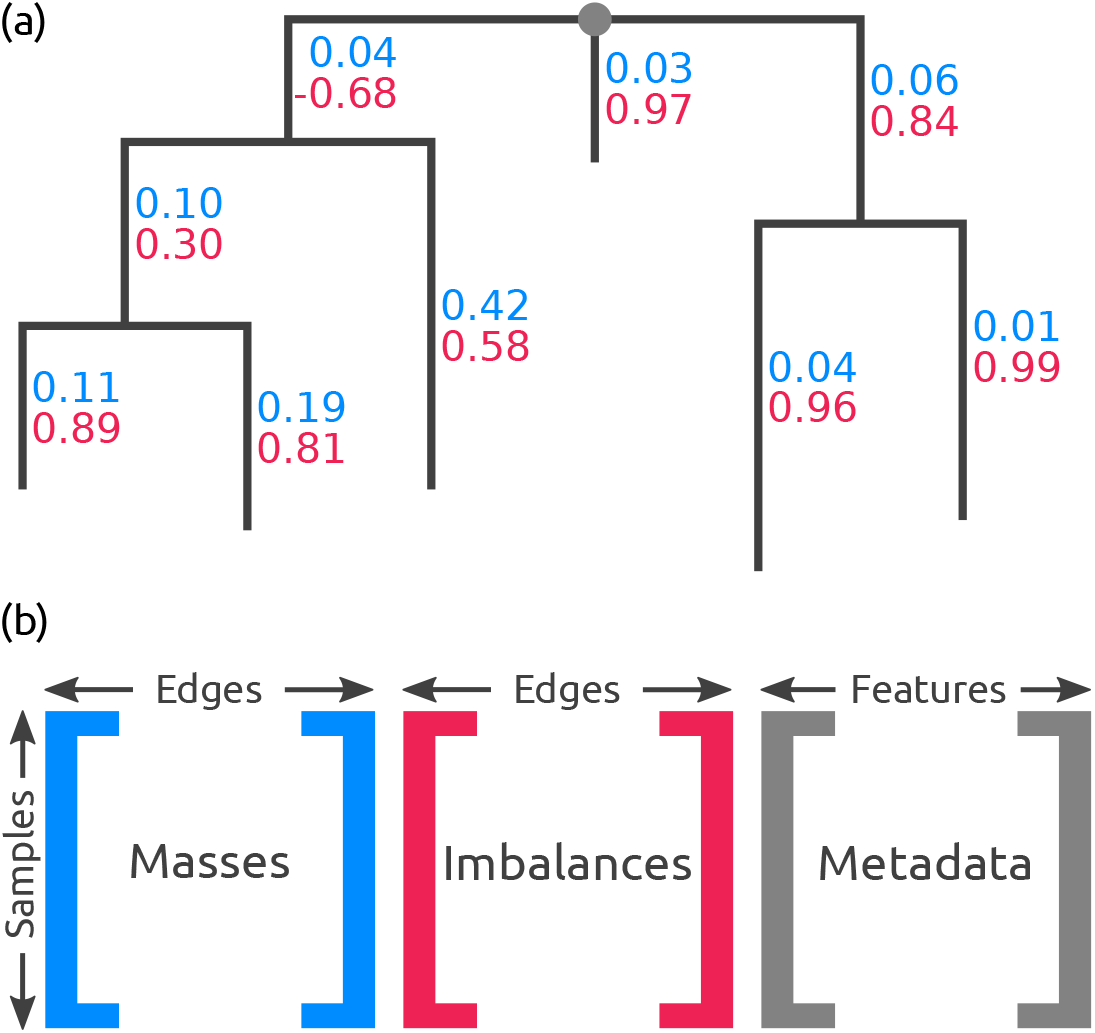
Edge Masses and Imbalances. (a) Reference tree where each edge is annotated with the normalized mass (first value, blue) and imbalance (second value, red) of the placements in a sample. The depicted tree is unrooted, hence, its top-level trifurcation (gray dot) is used as “root” node. (b) The masses and imbalances for the edges of a sample constitute the rows of the first two matrices. The third matrix contains the available meta-data features for each sample. These matrices are used to calculate, for instance, the Edge PCA or correlation coefficients.

The edge imbalance relates the masses on the two sides of an edge to each other. This implicitly captures the RT topology and reveals information about its clades. Furthermore, this transformation can also reveal differences in the placement mass distribution of nearby branches of the tree. This is in contrast to the KR distance [56], which we introduce later, as the KR distance yields low values for masses that are close to each other on the tree. Note that the imbalance of a leaf edge is simply the total mass of the tree minus the mass of the edge. It thus mostly contains irrelevant information and can often be omitted.

### 2.5 Placement data matrices

An example that shows all edge masses and edge imbalances of a sample on the reference tree is shown in Fig 3(a). Illustrations of the different use cases for edge mass and edge imbalance metrics are shown in the Results section. These values can be summarized by two matrices, which we use for several downstream edge- and clade-related analyses, respectively. These matrices have dimensions *n × m* for *n* samples and *m* edges of the RT. The edge masses matrix contains the sum of placement masses per edge for all QS in a sample, while the edge imbalance matrix transforms these masses as described before. Note that these matrices can either store absolute or relative abundances, depending on whether the placement mass was normalized.

Furthermore, many studies provide meta-data for their samples, for instance, the pH value or temperature of the samples’ environment. Such meta-data features can also be summarized in a per-sample matrix, where each column corresponds to one feature. The three matrices are shown in Fig 3(b). For example, the Edge principal components analysis (Edge PCA) [29] is a method that utilizes the imbalance matrix to detect and visualize edges with a high heterogeneity of mass difference between samples. Edge PCA further allows to annotate its plots with meta-data variables, for instance, by coloring, thus establishing a connection between differences in samples and differences in their meta-data [18]. In the following, we propose several new techniques to analyze placement data and their associated meta-data.

## 3 Methods

In this section, we introduce novel methods for analyzing and visualizing the phylogenetic placement data of a set of environmental samples. Each such sample represents a geographical location, a body site, a point in time, etc. In the following, we represent a sample by the placement locations of its metagenomic QSs, including the respective per-branch LWRs. Furthermore, for a specific analysis, we assume the standard use case, that is, all placements were computed on the same fixed reference tree (RT) and reference alignment. We initially describe our novel methods. We then assess their application to real world data and their computational efficiency in Section Results and discussion.

### 3.1 Visualization

A first step in analyzing phylogenetic placement data often is to visualize them. For small samples, it is possible to mark individual placement locations on the RT, as offered for example by iTOL [61], or even to create a tree where the most probable placement per QS is attached as a new branch, as implemented in the guppy tool from the pplacer suite [22], RAxML-EPA [23,62], and our tool gappa. For larger samples, one can alternatively display the per-edge placement mass, either by adjusting the line widths of the edges according to their mass, or by using a color scale, as offered in ggtree [63], guppy, and gappa. Using per-edge line widths or colors corresponds to binning all placements on an edge into one bin. For large datasets, the per-edge masses can vary by several orders of magnitude. In these cases, it is often preferable to use a logarithmic scaling, as shown in [15].

These simple visualizations directly depict the placement masses on the tree. When visualizing the accumulated masses of multiple samples at once, it is important to chose the appropriate normalization strategy for the task at hand. For example, if samples represent different locations, one might prefer to use normalized masses, as comparing relative abundances is common for this type of data. On the other hand, if samples from the same location are combined (e.g., from different points in time, or different size fractions), it might be preferable to use absolute abundances instead, so that the total number of sequences per sample can be visualized.

The visualizations provide an overview of the species abundances over the tree. They can be regarded as a more detailed version of classic abundance pie charts. Moreover, these visualizations can be used to assess the quality of the RT. For example, placements into inner branches of the RT may indicate that appropriate reference sequences (i) have not been included or (ii) are simply not yet available.

Here, we introduce visualization methods that highlight (i) regions of the tree with a high variance in their placement distribution (called *Edge Dispersion*), and (ii) regions with a high correlation to meta-data features (called *Edge Correlation*).

#### 3.1.1 Edge Dispersion

The Edge Dispersion is derived from the edge masses or edge imbalances matrix by calculating a measure of dispersion for each of the matrix columns, for example the standard deviation *σ*. Because each column corresponds to an edge, this information can be mapped back to the tree, and visualized, for instance, via color coding. This allows to examine which edges exhibit a high heterogeneity of placement masses across samples, and indicates which edges discriminate samples.

As the abundances of different species, and hence also the edge mass values, can span many orders of magnitude, it might be necessary to scale the variance logarithmically. Often, one is more interested in the branches with high placement mass. In these cases, using the standard deviation or variance is appropriate, as they also indicate the mean mass per edge. On the other hand, by calculating the per-edge Index of Dispersion [64], that is, the variance-mean-ratio *σ*^2^/*μ*, differences on edges with small mass also become visible. As Edge Dispersion relates placement masses from different samples to each other, the choice of the normalization strategy is important. When using normalized masses, the magnitude of dispersion values needs to be cautiously interpreted [59].

Edge Dispersion can also be calculated for edge imbalances. As edge imbalances are typically normalized to [−1.0,1.0], their dispersion can be visualized directly without any further normalization steps. An example for an Edge Dispersion visualization is shown in Fig 4(a), and discussed in Section Visualization.

**Fig 4.**
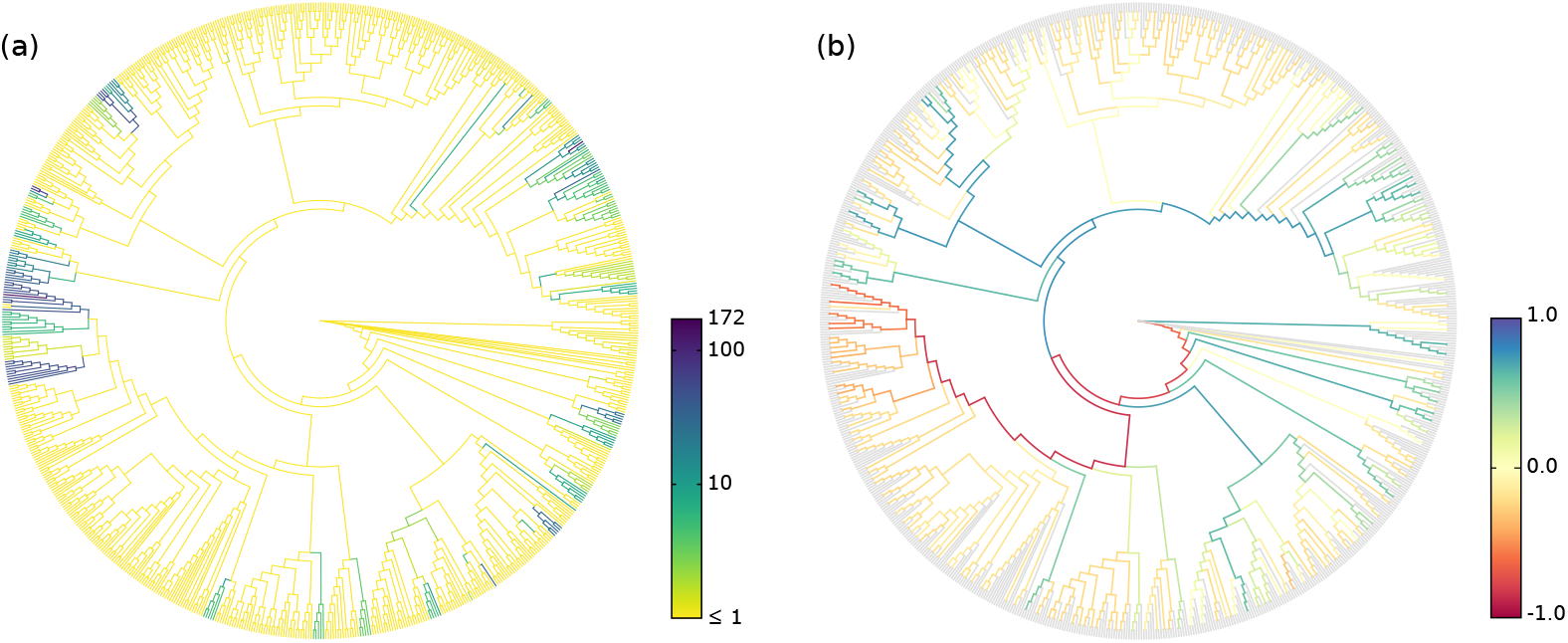
Examples of Edge Dispersion and Edge Correlation. We applied our novel visualization methods to the Bacterial Vaginosis (BV) dataset to compare them to the existing examinations of the data. (a) Edge Dispersion, measured as the standard deviation of the edge masses across samples, logarithmically scaled. (b) Edge Correlation, in form of Spearman’s Rank Correlation Coefficient between the edge imbalances and the Nugent score, which is a clinical standard for the diagnosis of Bacterial Vaginosis. Tip edges are gray, because they do not have a meaningful imbalance. This example also shows the characteristics of edge masses and edge imbalances: The former highlights individual edges, the latter paths to clades.

#### 3.1.2 Edge Correlation

In addition to the per-edge masses, the Edge Correlation further takes a specific meta-data feature into account, that is, a column of the meta-data matrix. The Edge Correlation is calculated as the correlation between each edge column and the feature column, for example by using the Pearson Correlation Coefficient or Spearman’s Rank Correlation Coefficient [64]. This yields a per-edge correlation of the placement masses or imbalances with the meta-data feature, and can again be visualized via a color coding of the edges. It is inexpensive to calculate and hence scales well to large datasets. As typical correlation coefficients are within [−1.0,1.0], there is again no need for further normalization. This yields a tree where edges or clades with either a high linear or monotonic correlation with the selected meta-data feature are highlighted. Fig 4(b) shows an example of this method.

In contrast to Edge PCA [29] that can use meta-data features to annotate samples in its scatter plots, our Edge Correlation method directly assesses the influence of a feature on the branches or clades of the tree. It can thus, for example, help to identify and visualize dependencies between species abundances and environmental factors such as temperature or nutrient levels. Again, the choice of the normalization strategy is important to draw meaningful conclusions. However, the correlation is *not* calculated between samples or sequence abundances. Hence, even when using normalized samples, the pitfalls regarding correlations of compositional data [59] do not apply here.

The method further bears some conceptual similarity to Phylofactorization [31], for which we later present an adaptation to phylogenetic placements, called *Placement-Factorization*. Phylofactorization also takes meta-data features into account and can thereby identify relationships between changes in environmental variables and changes in abundances in clades of the tree. It typically uses linear regression in form of a Generalized Linear Model (GLM) to assess these relationships. Note that the correlation coefficient used in our Edge Correlation can be interpreted as the slope of the regression line of the standardized values, which establishes a connection between Edge Correlation and Phylofactorization. The advantage of using correlations here instead of a GLM lies in its simplicity regarding result interpretation: Here, we are interested in whether changes in a meta-data feature are connected to an increase or decrease of abundances on branches or in clades of the tree, which is readily accessible via correlation coefficients.

However, using a GLM instead of correlation constitutes a potentially useful alternative, for example by visualizing the fit of the model with the data. This can be understood as fitting a regression line to the scatter plot of meta-data variables on one axis, and edge masses (i.e., per-branch abundances) on the other axis, and measuring how well this line predicts the data. We later discuss this idea in more detail in the context of the objective function of Placement-Factorization. Using GLMs in that way allows to assess the influence of multiple meta-data features simultaneously. For some types of meta-data, this might be advantageous, but might also hinder a clear interpretation of the relationships between different meta-data features and the abundances. We show some results of this idea as a by-product of Placement-Factorization in Fig 10 and S15 Fig, but leave a full exploration of this as a method of its own as future work.

### 3.2 Clustering

Given a set of metagenomic samples, one key question is how much they differ from each other. A common distance metric for microbial communities is the (weighted) UniFrac distance [57,58]. It uses the fraction of unique and shared branch lengths between phylogenetic trees to determine their difference. UniFrac has been generalized and adapted to phylogenetic placements in form of the phylogenetic Kantorovich-Rubinstein (KR) distance [29,56]; see there for a thorough comparison of UniFrac and KR distance. In other contexts, the KR distance is also called Wasserstein distance, Mallows distance, or Earth Mover’s distance [65–68]. The KR distance between two metagenomic samples is a metric that describes by at least how much the normalized mass distribution of one sample has to be moved across the RT to obtain the distribution of the other sample. The distance is symmetrical, and increases the more mass needs to be moved, and the larger the respective displacement (moving distance) is. As the two samples being compared need to have equal masses, the KR distance operates on normalized samples; that is, it compares relative abundances.

Given such a distance measure between samples, a fundamental task consists in clustering samples that are similar to each other. Standard linkage-based clustering methods like UPGMA [69–71] are solely based on the distances between samples, that is, they calculate the distances of clusters as a function of pairwise sample distances. Squash Clustering [18,29] is a method that also takes into account the intrinsic structure of phylogenetic placement data. It uses the KR distance to perform agglomerative hierarchical clustering of samples. Instead of using pairwise sample distances, however, it merges (*squashes*) clusters of samples by calculating their average per-edge placement mass, as described in the Introduction. A further example of the squashing of two samples is shown in Fig 5. Thus, in each step, Squash Clustering operates on the same type of data, that is, on mass distributions on the RT: In the beginning, each item considered in the clustering is one sample, while later steps of the clustering operate on the merged (squashed) samples, until only one large cluster remains that combines all samples. This results in a hierarchical clustering tree, where tips correspond to samples, and branch lengths correspond to KR distances between clusters. We show examples of this clustering tree later in Fig 8.

**Fig 5.**
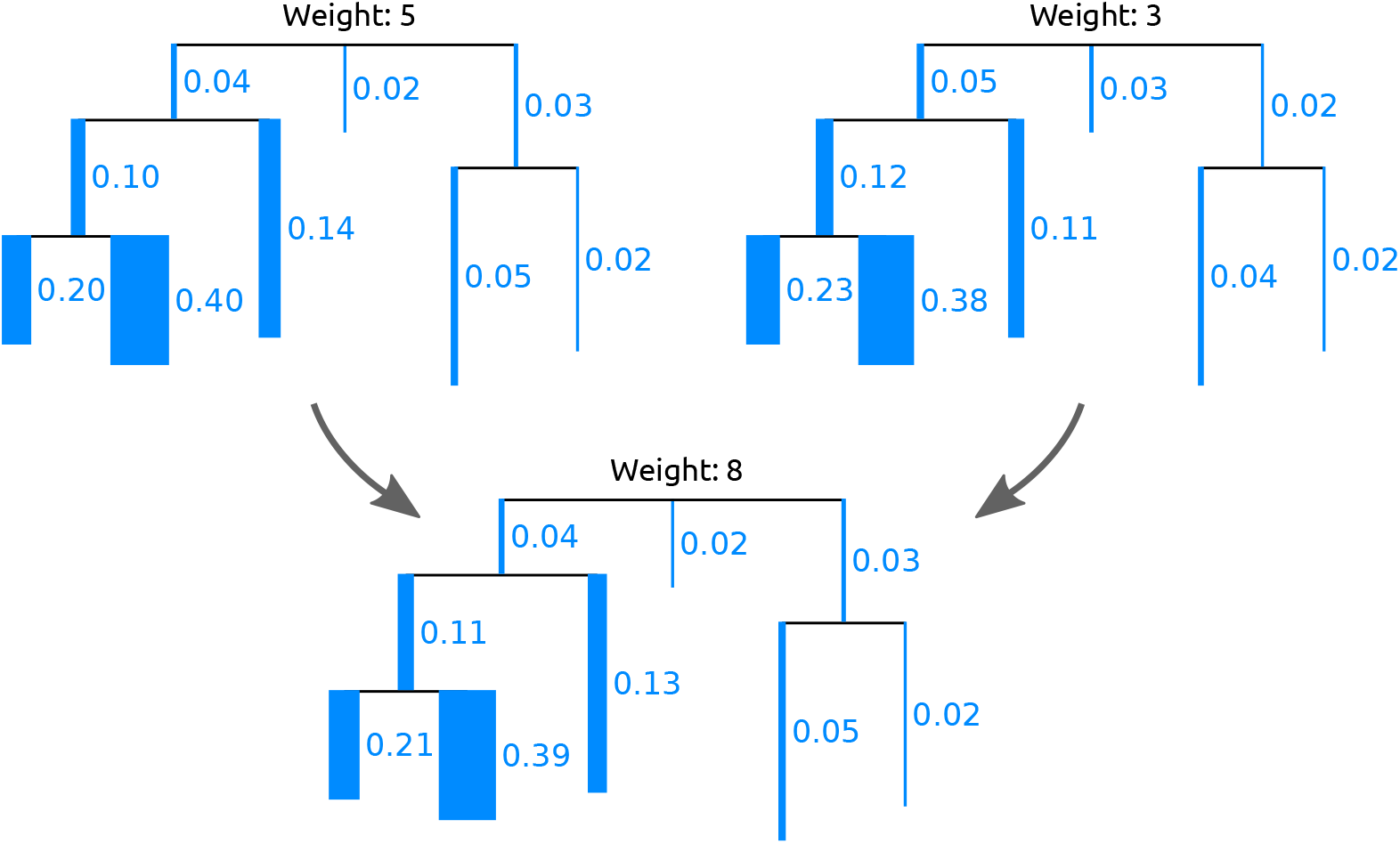
Squashing of Edge Masses. Two trees are merged (*squashed*) by calculating the weighted average of the respective mass distributions on their branches. By squashing, a cluster of (similar) samples can be summarized and visualized. For simplicity, we here show the masses per edge and visualize them as branch widths. In practice however, each placement location of each query sequence is considered individually. The Figure is based on the similar Figure 3/2 of Matsen *et al*. [29]; see there for more details on squashing.

#### 3.2.1 Phylogenetic *k*-means

The number of tips in the resulting clustering tree obtained through Squash Clustering is equal to the number *n* of samples that are being clustered. Thus, for datasets with more than a few hundred samples, the clustering result becomes hard to inspect and interpret visually. We propose a variant of *k*-means clustering [72] to address this problem, which we call *Phylogenetic k-means*. It uses a similar approach as Squash Clustering, but yields a predefined number of *k* clusters. It is hence able to work with arbitrarily large datasets. Note that we are clustering samples here, instead of sequences [73]. We discuss the choice of a reasonable value for *k* later.

The underlying idea is to assign each of the *n* samples to one of *k* cluster centroids, where each centroid represents the average mass distribution of all samples assigned to it. Note that all samples and centroids are of the same data type, namely, they are mass distributions on a fixed RT. It is thus possible to calculate distances between samples and centroids, and to calculate their average mass distributions, as described earlier. Our implementation follows Lloyd’s algorithm [74], as shown in Algorithm 1.

##### Algorithm 1 Phylogenetic *k*-means

~~~
1: initialize *k Centroids*
2:   **while** not converged **do**
3:   assign each *Sample* to nearest *Centroid*
4:   update *Centroids* as mass averages of their *Samples*
5:**return** Assignments and *Centroids*
~~~

By default, we use the *k*-means++ initialization algorithm [75] to obtain a set of *k* initial centroids. It works by subsequent random selection of samples to be used as initial centroids, until *k* centroids have been selected. In each step, the probability of selecting a sample is proportional to its squared distance to the nearest already selected sample. An alternative initialization is to select samples as initial clusters entirely at random. This is however more likely to yield sub-optimal clusterings [76].

Then, each sample is assigned to its nearest centroid, using the KR distance. Lastly, the centroids are updated to represent the average mass distribution of all samples that are currently assigned to them. This iterative process alternates between improving the assignments and improving the centroids. Thus, the main difference to normal *k*-means is the use of phylogenetic information: Instead of euclidean distances on vectors, we use the KR distance, and instead of averaging vectors to obtain centroids, we use the average mass distribution on the tree.

The process is repeated until it converges, that is, the cluster assignments do not change any more, or until a maximum number of iterations have been executed. The second stopping criterion is added to avoid the super-polynomial worst case running time of *k*-means, which however almost never occurs in practice [77,78].

The result of the algorithm is an assignment of each sample to one of the *k* clusters. As the algorithm relies on the KR distance, it clusters samples with similar relative abundances. The cluster centroids can be visualized as trees with a mass distribution, analogous to how Squash Clustering visualizes inner nodes of the clustering tree. That is, each centroid can be represented as the average mass distribution of the samples that were assigned to it, as shown in Fig 5. This allows to inspect the centroids and thus to interpret how the samples were clustered. Examples of this are shown in S8 Fig.

The key question is how to select an appropriate *k* that reflects the number of “natural” clusters in the data. There exist various suggestions in the literature [79–84]; we assessed the Elbow method [79] as explained in S10 Fig, which is a straight forward method and yields reasonable results for our test datasets. Additionally, for a quantitative evaluation of the clusterings, we used the *k* that arose from the number of distinct labels based on the available meta-data for the data. For example, the HMP samples are labeled with 18 distinct body sites, describing where each sample was taken from, c.f. Fig 9.

#### 3.2.2 Algorithmic improvements

In each assignment step of the algorithm, distances from all *n* samples to all *k* centroids are computed, which has a time complexity of 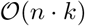. In order to accelerate this step, we can apply branch binning as introduced in Section Edge masses. For the BV dataset, we found that even using just 2 bins per edge does not alter the cluster assignments. Branch binning reduces the number of mass points that have to be accessed in memory during KR distance calculations; however, the costs for tree traversals remain unaltered. Thus, we observed a maximal speedup of 75% when using one bin per branch, see S2 Table for details.

Furthermore, during the execution of the algorithm, empty clusters can occur, for example, if *k* is greater than the number of natural clusters in the data. Although this problem did not occur in our tests, we implemented the following solution: First, find the cluster with the highest variance. Then, choose the sample of that cluster that is furthest from its centroid, and assign it to the empty cluster instead. This process is repeated if multiple empty clusters occur at once in an iteration.

#### 3.2.3 Imbalance *k*-means

We further propose *Imbalance k-means*, which is a variant of *k*-means that makes use of the edge imbalance transformation. In order to quantify the difference in imbalances between two samples, we use the Euclidean distance between their imbalance vectors (that is, rows of the imbalance matrix). This is a suitable distance measure, as the imbalances implicitly capture the tree topology as well as the placement mass distributions. As a consequence, the expensive tree traversals required for Phylogenetic *k*-means are not necessary here. The algorithm takes the edge imbalance matrix of normalized samples as input, as shown in Fig 3(b), and performs a standard Euclidean *k*-means clustering following Lloyd’s algorithm.

This variant of *k*-means tends to find clusters that are consistent with the results of Edge PCA, as both use the same input data (imbalances) as well as the same distance measure (Euclidean). Furthermore, as the method does not need to calculate KR distances, and thus does not involve tree traversals, it is several orders of magnitude faster than Phylogenetic *k*-means. For example, on the HMP dataset, it runs in a few seconds, instead of several hours needed for Phylogenetic *k*-means; see Section Performance for details.

### 3.3 Phylogenetic ILR transform and Phylofactorization

The concepts and methods presented above resemble two recent approaches for analyzing phylogenetic data: the Phylogenetic Isometric Log-Ratio (*PhILR*) transformation and balances [30], as well as Phylogenetic Factorization (*Phylofactorization*) [31]. These methods use a tree inferred from the OTU sequences of the samples (instead of a fixed reference tree), and annotate the abundances of OTUs per sample on the tips of this tree (instead of placement masses on the branches). The methods use these data to draw conclusions about compositional changes in clades of the tree in different samples, as well as relationships of per-clade OTU abundances with environmental meta-data variables.

In both of these approaches, a *balance* between OTU abundances in two subtrees of the underlying tree is computed. This is a measure of contrast that expresses which of the two subtrees comprises more OTUs. In the PhILR transform [30], these balances are computed for the two subtrees below each inner node of a rooted binary tree, while ignoring abundances in the respective remainder of the tree. In Phylofactorization however, these balances are computed for the two subtrees that are induced by the edges of the tree. This is highly similar to the concept of edge imbalances that we introduced above. Note that despite sharing a similar name and exhibiting conceptual similarities, balances and the previously introduced edge imbalances are distinct approaches that should not be confused. We later discuss respective similarities and differences in more detail.

Furthermore, we remark that the *Balance Trees* method [85] employs analogous concepts by calculating the balance of nodes using the isometric log-ratio of OTU abundances. However, instead of using a phylogenetic tree, it assumes any binary partitioning of the OTUs, e. g., obtained from a UPGMA clustering of the OTUs based on a meta-data feature. These nodes thus correspond to specific meta-data values, again allowing for statements about the changes in OTU abundances that occur with changing environmental variables. As we already have a binary partitioning in form of the reference tree, we do not further consider the Balance Trees approach here.

In the following, we present adaptations of the PhILR transform (balances) and of Phylofactorization to phylogenetic placement data. The main adaptation step consists in placing masses on the branches of our (fixed) reference tree, instead of only considering masses (abundances) at the tips of the OTU tree. Here, we focus on balances that contrast the subtrees induced by edges of the tree, as used by Phylofactorization [31], because this is more natural in the context of phylogenetic placement data. The same concepts could however also be employed for subtrees below nodes, as used by the PhILR transform [30].

#### 3.3.1 Phylogenetic ILR transform for phylogenetic placements

In the introduction, we briefly explained the inherently compositional nature of metagenomic sequence data [42]. For a thorough discussion of the implications of this, see [30]. One solution is to transform the data into an unconstrained space that is not compositional. This can, for example, be achieved via the Isometric Log-Ratio (ILR) transform [86], which, given a compositional space, creates a new coordinate system with an orthonormal basis [87]. The ILR transform requires a sequential binary partitioning of the underlying original space [88]. As suggested in [30], a bifurcating phylogenetic tree (e.g., our RT) represents such a partitioning, which also provides a meaningful way of interpreting the resulting coordinates. This so-called *Phylogenetic ILR* (PhILR) transform yields an ILR coordinate system that captures the evolutionary relationships of the phylogeny [30]. The resulting coordinates are called *balances* [86,87]. The balances obtained from an ILR transform represent the log-ratio of the geometric means of the data in the two subtrees. Hence, they can be interpreted as a contrast (log-ratio) between two aggregates (geometric means). Furthermore, due to the orthogonality of the ILR basis vectors, the balances can be used by conventional statistical tools without suffering from compositional artifacts.

In the following, we present an adaptation of the PhILR transform and balances to placement data, based on the work of [30]. See there for more details on the method and the underlying mathematical concepts, such as the connection between the ILR transform and the centered log-ratio (CLR) transform. We now describe the computation of the Phylogenetic ILR transform, along with the changes needed for phylogenetic placement data. We focus on the computation for a single sample. We assume that a fixed reference tree (RT) (a sequential binary partitioning) along with the per-branch placements of the sequences in the sample are given. The placements are represented by a vector ***c*** of size *m*, containing the absolute (not normalized) edge masses, where *m* is the number of edges in the tree. In other words, our input is a single row (one sample) of the edge masses matrix, as shown in Fig 3(b). The absolute masses are transformed into relative abundances as described in Section Edge masses: Each element of ***c*** is divided by the sum of all elements, yielding the relative masses vector ***x*** for the given sample. In compositional data analysis, this operation is known as the closure of the data [43].

The original PhILR furthermore allows to use per-taxon weights ***p*** in order to down-weigh the impact of low abundant taxa/OTUs [30,89]. In our adaptation, this weighting scheme is accordingly changed to weights ***p*** *per edge* of the RT. Unfortunately, the nomenclature of existing publications (namely, [29] and [30]) creates a conflict here: These weights are not to be confused with our terminology of likelihood weights and edge masses. The default case of edge weights ***p*** = (1,…, 1) represents no weighting, where each edge equally contributes to the balance, while any ***p*** ≠ (1,…, 1) is a generalized form of the ILR transform [30]. We later describe an appropriate choice of weights in Section Edge Weights. These weights are applied to the relative masses *x* to obtain the shifted composition ***y = x/p***, using element-wise division [89].

In the original PhILR, balances are calculated for the two subtrees below a given node of the tree [30]. In the context of Phylofactorization, this has been generalized to balances between any two disjoint sets *R* and *S* of taxa (tips of the tree) [31]. We here build on the latter, but again change *R* and *S* to refer to disjoint sets of edges of our reference tree. We use the notation ***y***_*R*_ and ***p***_*R*_ to refer to the subsets of masses and weights of the given sample at the edges in *R*. Then, the balance *y*\* between the sets *R* and *S* is computed as:

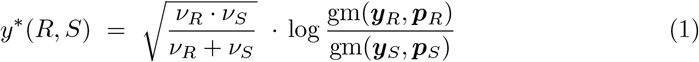

The first term is a scaling term that, for a given edge, is constant across all samples. It ensures unit length of the ILR basis elements, and uses the sums of weights in ***p***:

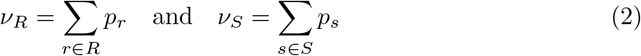

The second term is the log-ratio of geometric means, where gm(***y***_*R*_, ***p***_*R*_) is the weighted geometric mean of the values in ***y***_*R*_ with weights ***p***_*R*_:

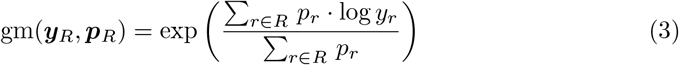

Note that if ***p*** = (1,…, 1), Eq (1) represents the original ILR transform without a weighting scheme [86], Eq (2) equals the number of edges in *R* and *S*, respectively, and Eq (3) is the standard (unweighted) geometric mean.

Balances as defined here can be computed as a measure of contrast between any disjoint sets *R* and *S* of edges. Interchanging *S* and *R* flips the sign of the balance; this is irrelevant for the subsequent steps presented here, as long as the interchange is applied consistently. When computing the balance between the edges in the two subtrees induced by some given edge e, the conceptual similarity with the previously described Edge imbalances becomes apparent: Imbalances use the difference of sums for contrasting and aggregating, while balances use the ratio of means for the same purpose. We show an example of this computation in Fig 6. Hence, balances represent a similar transformation of the placement data, that can also be used to conduct analyses, such as the Phylofactorization, as presented in the next section.

**Fig 6.**
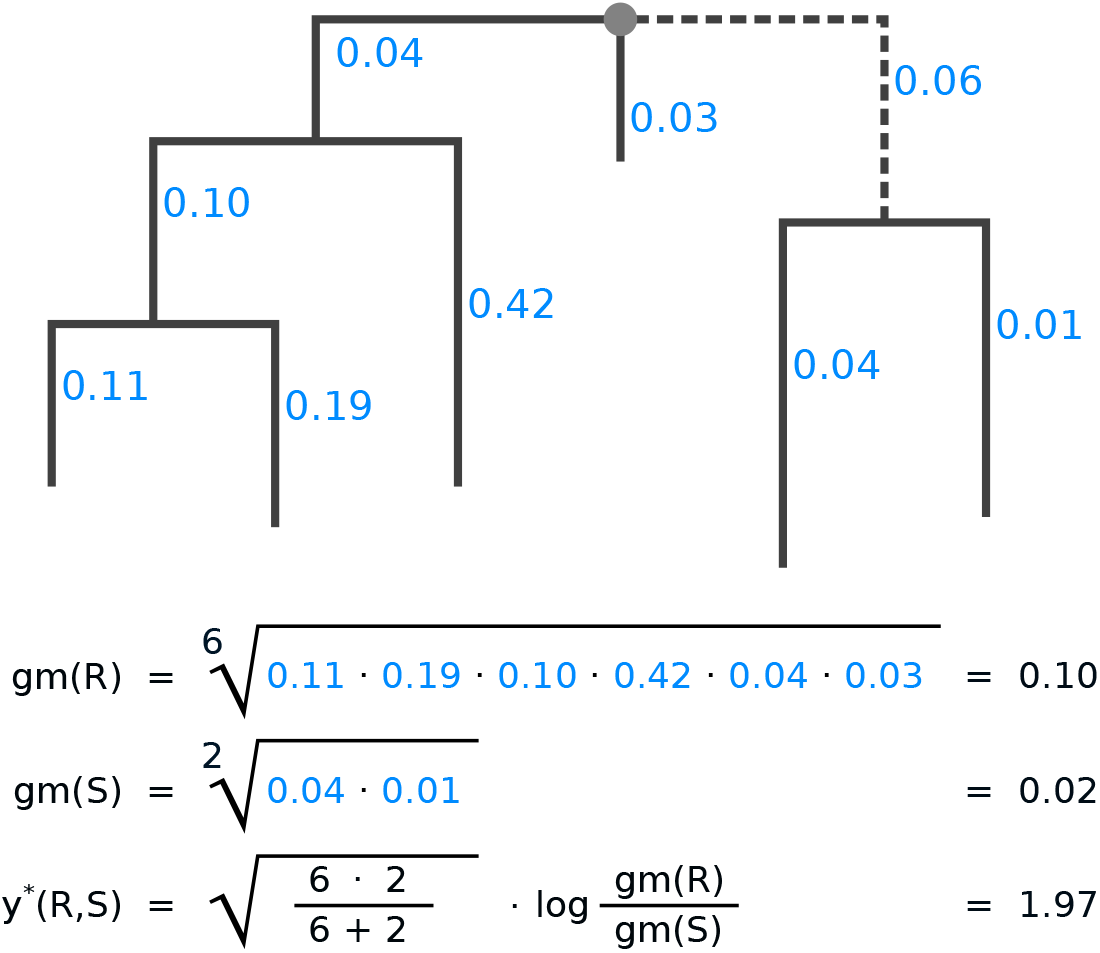
Example computation of the balances between two subtrees. The figure shows how the balance is computed for the two subtrees induced by the dashed edge of the tree, for one sample. Numbers next to edges are the accumulated placement mass of the sequences in the sample. We call the left hand side of the tree R, and the right hand side S, as seen from the dashed edge. For simplicity, we do not use weighting here; that is, we assume ***p*** = (1,…, 1). First, the geometric means for both subtrees are calculated, then, their balance. The balance is positive, indicating that subtree R contains more placement mass on (geometric) average.

We however remark that the using (unweighted) balances in our previously presented methods, such as Edge Correlation and *k*-means clustering, might lead to spurious results, due to the insensitivity of the geometric mean to singular large values. That is, individual branches that accumulate a large fraction of the placement mass (sequence abundances) might only insignificantly change the geometric mean of their clade.

However, such branches are typically the interesting ones, and hence should exert more influence on the transformation, which is exactly the purpose of the taxon weighting scheme. This further implies that balances are not indifferent to splittings of reference taxa into multiple representatives (pers. comm. with A. Washburne on 2018-11-23). We discuss the implications of this in more detail in the evaluation of the method.

#### 3.3.2 Edge Weights

The PhILR also allows for incorporating two distinct weighting schemes for the balances, one based on taxon abundances, and one based on the branch lengths of the underlying phylogeny [30]. As mentioned above, we implemented the former, while leaving the latter as future work.

We now describe how to adapt the taxon weights of [30] to our placement-based approach, that is, how an appropriate vector ***p*** of edge weights can be constructed. Originally, this weighting scheme down-weighs the influence of low abundant taxa [30], which are known to be less reliable and more variable [90]. Here, we accordingly down weigh edges with low placement mass, for the same reasons. We follow the approach of [30], and construct the edge weights by multiplicatively combining two terms:

- A measure for the central tendency of the absolute edge masses, for example, their mean across all samples. This is the main component of the weight that yields low values for edges with low mass and vice versa.
- A vector norm of the relative edge masses across the samples. This term additionally weighs edges by their specificity.

Our implementation allows to use the median, the arithmetic mean, and the geometric mean, as well as different *ℓ_p_*-norms (such as the Manhattan, Euclidean, and maximum norm), and the Aitchison norm [88]. We follow the advice of [30], and by default use the geometric mean (with pseudo-counts added to the masses to avoid skew from edges without any placement mass) and the Euclidean norm. In that case, the weights for edge *j* are computed as follows:

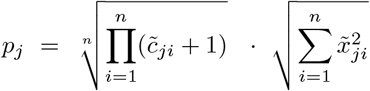

Here, *n* is the number of samples, 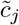 is the vector of absolute masses at edge *j* across all samples, and 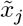 the vector of relative masses at edge *j* across all samples, both of length *n*. That is, these measures use the masses of all *n* samples; consequently, we use columns instead of rows of the edge masses matrix of Fig 3(b), where each column is used for the weights of the corresponding edge. The resulting edge weights ***p*** are then fixed and used across the balance computation of all samples.

### 3.4 Phylogenetic Factorization

Phylofactorization is a method to identify edges in a phylogenetic tree that drive patterns in the composition of microbial communities [31]. An edge constitutes a separation or split of groups of taxa into the two subtrees induced by the edge. In an evolutionary context, an edge denotes a difference in (putative) traits that may have arisen along the edge. The goal of Phylofactorization is to identify edges that are related to differences in per-sample meta-data features. To this end, it aggregates and contrasts the abundances in the subtrees (groups of taxa) induced by an edge, and evaluates how changes in environmental variables across samples are reflected in abundance changes.

The original method uses a tree inferred from the OTUs that are present in the set of samples, and iteratively identifies edges that split the tree into nested subtrees which exhibit the largest predictable differences between the taxa in these subtrees. Each such edge can be interpreted as a *phylogenetic factor* (or short, *phylofactor*) for splitting the tree: Once an edge has been selected in one iteration, its induced subtrees are then considered separately in subsequent iterations. The resulting factors are hence independent of each other, which ensures orthogonality of the factors. In other words, each factor describes a different dimension in which samples differ. Furthermore, by iteratively considering subtrees of decreasing size, nested factors can be found, which correspond to relationships within a subtree that only affect the taxa in the subtree itself. The algorithm stops after a predefined number of iterations/factors, or until a stopping criterion is met.

In a typical use case, each environmental sample is represented by its OTU abundances at the tips of the tree. Given a per-sample meta-data feature such as the pH-value, Phylofactorization can be employed to find edges where a change of the pH-value between samples predicts a change in OTU abundances in the two subtrees induced by the edge. For example, an increasing ph-value might indicate a relative increase in the OTU abundances in one subtree compared to another subtree. The resulting factorization can serve as a dimensionality-reduction mechanism, as an ordination and visualization tool, and as an inferential tool that can identify edges corresponding to changes in functional ecological traits [31].

We now present an adaptation of Phylofactorization to phylogenetic placements, which we call *Placement-Factorization*. We explain our adaptation following the description of the Generalized Phylogenetic Factorization (GPF) [91,92]. The GPF is a recent generalization of Phylofactorization that also allows for other types of input data than relative OTU abundances, for example, presence/absence data. It is hence suited for a wider range of community ecological data [92]. Conceptually and algorithmically, Phylofactorization, GPF, and our Placement-Factorization, work the same; we here use the mathematical notation of GPF as a scaffold to explain our adaptation. In the following, we briefly introduce the original method, outline the necessary adaptations, and explain how to use the balances obtained from (our adaptation of) the PhILR transform (as explained above) in the context of Placement-Factorization.

#### 3.4.1 Placement-Factorization

Phylofactorization can be understood as an iterative greedy graph-partitioning algorithm for a tree *T* [91]. In each iteration, a *winning edge e*\* is identified that splits the tree into two disjoint groups *R* and *S* of edges. To determine the winning edge, an *objective function* is maximized that expresses the intensity of the relationship between abundances and meta-data variables. We later discuss this objective function in more detail.

The input to the original Phylofactorization is an *n × m* data matrix *X*, for *j* = 1,…,*n* samples, and *i* = 1,…,*m* species (corresponding to the OTUs at the tips of the tree). The matrix can represent abundances, presence/absence data, or other data related to the species in the tree [92]. In our adaptation, we use the per-edge masses from the phylogenetic placement of the samples, as shown before in Fig 3(b). That is, instead of *m* species representing the tips of the tree, we use an *n* × *m* data matrix *X* where the *m* columns correspond to the edges of our reference tree (for consistency of notation, we re-use and re-purpose the index *m* here, and transpose *X* compared to the original notation). Lastly, Phylofactorization uses an *n × p* meta-data matrix *Z* for the *n* samples and *p* per-sample meta-data variables.

In analogy to the Generalized Phylofactorization [91,92], our adapted algorithm requires three functions:

1. An *aggregation function A_R_* = *A*(*X_j,R_, T*), which aggregates (summarizes) a subset *R* of edges for a sample *j*.
2. A *contrast function C_R,S_* = *C*(*A_R_, A_S_, T, e*), which contrasts (compares) the aggregates of two disjoint subsets *R* and *S* of edges on the two sides of an edge *e*.
3. An *objective function ω*(*C, Z*) that evaluates a contrast for all samples in the context of the per-sample meta-data, in order to determine the winning edge.

We later discuss appropriate choices for these functions. For now, we assume that we are given functions that allow identifying edges whose induced subtrees exhibit predictable differences in the edge masses *X* driven by changes in the meta-data *Z* of different samples. The algorithm starts by considering the entire tree *T* as one large “subtree”. Then, in each iteration, Phylofactorization and Placement-Factorization work as follows:

1. For each edge *e* that separates disjoint groups *R_e_* and *S_e_* of edges within the subtree that contains *e*:

a. Compute the aggregates *A_R_e__* = *A*(*X_j,R_e__, T*) and *A_S_e__* = *A*(*X_j,S_e__, T*).
b. Compute their contrast *C_e_* = *C*(*A_R_e__, A_S_e__, T, e*).
c. Compute the objective value *ω_e_* = *ω*(*C_e_, Z*). The aggregates *A_R_e__* and *A_S_e__*, as well as the contrast *C_e_* are computed separately for every sample. The value *ω_e_* of the objective function then expresses the relationship of the contrasts of all samples with their respective meta-data values in *Z*.
2. Select the winning edge *e*\* = argmax_*e*_(*ω_e_*) that maximizes the value of the objective function.
3. Partition the subtree that contains *e*\* into two disjoint subtrees, separated by *e*\*.
4. Repeat until a stopping criterion is met.

This closely follows the description of the algorithm in [91,92], see there for details. The difference between the algorithms is that the groups *R* and *S* in our case consist of reference tree edges, instead of species at the tips of the OTU tree. Because of this, the aggregates of edges that lead to tip nodes are empty, meaning that we do not consider those edges as candidates in the algorithm. This is analogous to “tip edges” not having a meaningful edge imbalance, as described above. An example of the first two iterations of the algorithm is shown in Fig 7.

**Fig 7.**
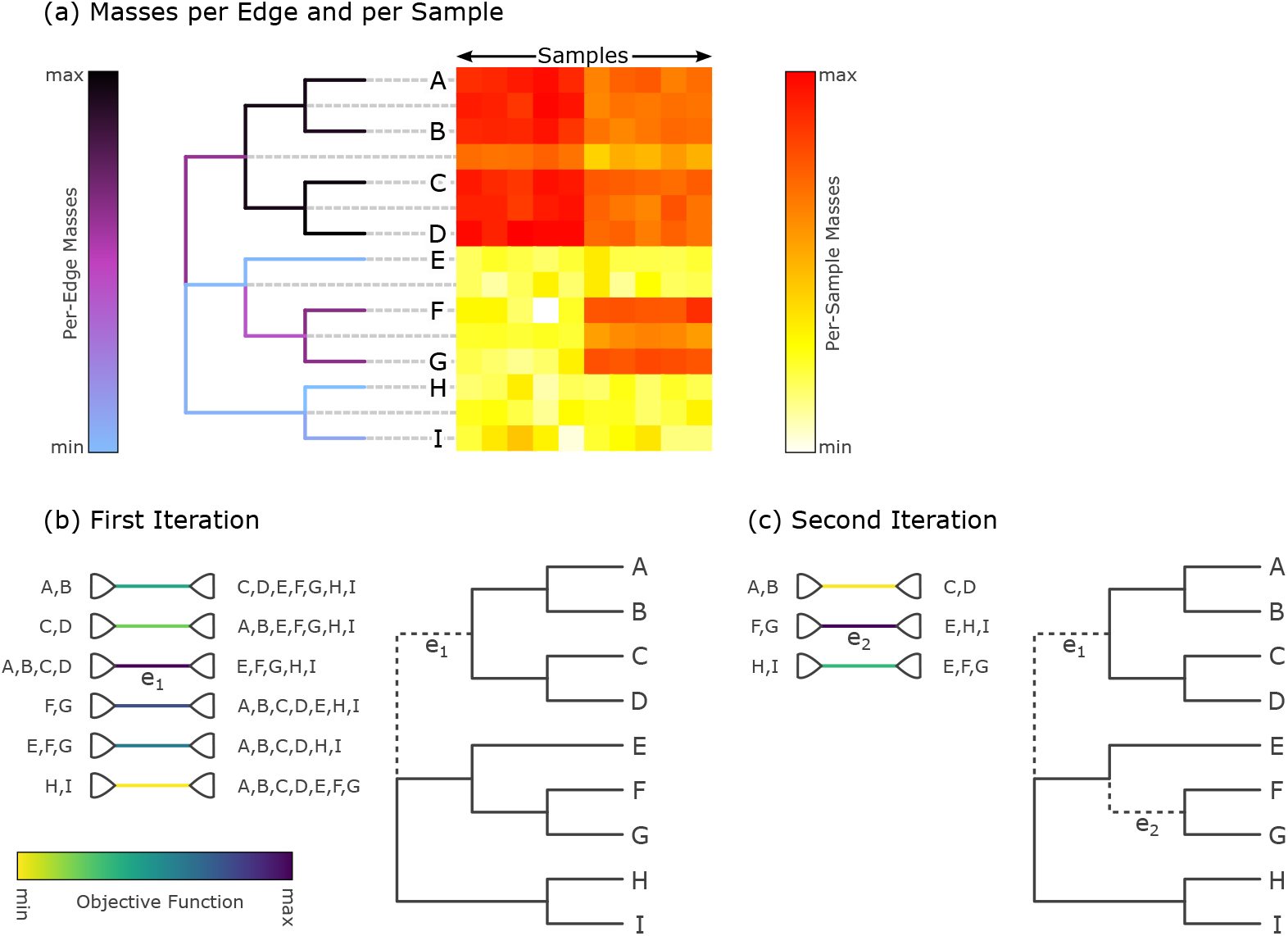
Input data and first two iterations of Placement-Factorization. The figure resembles Figure 2 of [31]. It shows the adaptation of concepts from Phylofactorization to phylogenetic placement data. (a) The input data is a set of samples with placement masses on each edge of the tree. The tree is colorized by the total mass across all samples, that is, by the row sums of the heat map. The heat map then shows the detailed mass per edge (rows) and per sample (columns). Note that the heat map also contains rows for each inner edge of the tree, as phylogenetic placement also considers these edges. We show an example of this visualization for empirical data in S1 Fig. (b) In the first iteration, the objective function for all inner edges is evaluated. Here, *e*_1_ is the winning edge that maximizes the objective function, which separates (`A, B, C, D`) from the rest of the tree. (c) In the second iteration, only the contrasts within the two subtrees are calculated, but not across the winning edges of previous iterations (here, e_1_). That is, the winning edge *e*_2_ maximizes the objective function that contrasts (F, G) with (`E, H, I`), but does not consider the edges in the subtree (`A, B, C, D`). Note that in our adaption, edges that lead to a tree tip are not considered as potential factors.

Each iteration further splits a subtree at the respective winning edge, so that after *i* iterations, *i* + 1 subtrees are produced. It is important to note that the winning edges of previous iterations split the tree into *disjoint* subtrees, and that in later iterations, the aggregates and contrasts induced by an edge are only computed *within* their respective subtrees. This ensures the previously mentioned orthogonality of the phylogenetic factors (winning edges), meaning that systematic dependencies between the contrasts of any two factors are eliminated, and that instead, nested relationships can be identified.

The original publication proposes a stopping criterion using a Kolmogorov-Smirnov (KS) test based on test statistics of the identified phylofactors [31,92]. Although these could be implemented for Placement-Factorization, we leave this as future work; our implementation currently runs for a given number *i* of iterations, and hence computes *i* phylofactors.

So far, we have assumed to be given the three functions required for Phylofactorization. The choice of these functions depends on the data *X*, the data *Z*, and the research question at hand. In order to be consistent and comparable with the original implementation [31], in our evaluation we used the same set of functions, namely the balances of the ILR transformation as explained above for aggregating and contrasting subtrees, and an objective function based on Generalized Linear Models (GLMs), which we explain in the following.

#### 3.4.2 Objective Function

Phylofactorization requires an objective function *ω*(*C_e_, Z*) that quantifies the relationship between *C_e_* and *Z* for a given edge *e*, where *C_e_* are the contrasts between the two subtrees induced by *e* for all samples (for example, the balances), and *Z* are the per-sample meta-data variables. That is, both *C_e_* and *Z* have size n, the number of samples, with *Z* potentially containing multiple columns (one for each meta-data feature). In order to identify the winning edge *e*\* of an iteration (the *phylofactor*), the function is evaluated for all edges, and the edge maximizing *ω* is selected. The choice of the objective function depends on the research question at hand; see [31] and [91] for a thorough discussion.

Our implementation is as general as the original Phylofactorization [31], in that it allows for an arbitrary objective function. For simplicity, and in line with the original publication, we here focus on functions that treat the meta-data variables *Z* as independent variables and the contrasts *C_e_* as dependent variables whose relationship with *Z* is assessed, for instance, via a predictive model. Then, the selected phylogenetic factors correspond to edges where a change in *Z* most strongly predicts a change in *C_e_* across the samples, that is, where the effect of the (independent) meta-data variables on the (dependent) underlying data (e.g., per-clade abundances) is most pronounced.

A powerful approach is to model the relationship between *C_e_* and *Z* via linear regression, that is, we assess how well *Z* can predict *C_e_*. In the simple one-dimensional case, this can be thought of as fitting a line through a scatter plot of the meta-data feature on the *x*-axis and the contrasts on the *y*-axis, where each point represents one sample. This concept is generalized via Generalized Linear Model (GLM) [93–95].

In short, GLMs allow to predict a single (response) variable using multiple input (explanatory) variables. Typically, the response variable is assumed to follow any distribution from the exponential family (normal, exponential, Poisson, Binomial, etc), which is given for balances as used here. In contrast to this, the explanatory variables (the meta-data features) are assumed to have a linear relationship with the response. Note that this mathematical restriction of the model does not mean that only meta-data features can be used that behave linearly; transformations and interactions of the features basically allow for arbitrary types of data. For example, categorical variables such as the body site where a sample was taken from can be transformed into so-called dummy variables that fulfill the requirements.

Once the model parameters of the GLM have been estimated, that is, once it has been fit to the data via some optimization algorithm, we need to evaluate the GLM for the purposes of Phylofactorization. We are interested in a value for *ω* that expresses how well the meta-data variables explain the balances. To this end, Phylofactorization and our adaptation thereof use the difference between the null deviance of the balances and the deviance obtained from the GLM. This difference expresses how much better the model explains the balances than just predicting them from their mean. For details on the usage of GLMs for Phylofactorization, see [31].

Predictive models such as GLMs expect the response variable (that is, the predicted values; here, the contrasts) to have certain statistical properties. In particular, linear models assume the deviation of response from the predicted value to be normally distributed. The ILR transform for compositional data has been proven to behave asymptotically normal [86,96], which allows their application within standard multivariate methods, and within GLMs as presented here.

Lastly, we note that depending on the research question, other objective functions can be used, see [31,91] for some examples. For instance, simple test statistics such as the variation in *C_e_* explained by regression on *Z* can be used. Furthermore, instead of predicting contrasts from meta-data, one could be interested in the opposite, that is, predicting a meta-data variable given the per-sample contrasts. In this case, the maximization of the objective function yields edges that best predict a certain feature of the data; this is suitable for identifying clades that can serve as a bio-indicator. Using GLMs for this allows to model any type of meta-data variable; for example, the binary information encoded in presence/absence data can be predicted using logistic regression. While our implementation supports all those use cases, they have been explored and discussed before [31,92]. For the sake of simplicity, we focus on linear (gaussian) modeling of *C_e_ ~ Z*, that is, predicting balances from meta-data.

#### 3.4.3 Phylofactorization for phylogenetic placements

In summary, Phylofactorization and our adaptation Placement-Factorization identify edges of the phylogeny that exhibit a predictable relationship between changes in meta-data variables and abundance changes in the subtrees induced by these edges. Our adaptation can be understood as a generalization of the original method [31,92], where masses/counts can be placed along the edges of the tree, instead of just at its tips.

While the original method uses abundances of taxa/OTUs per sample on a tree inferred from the OTU sequences, we use the placement masses on a fixed reference tree (RT). For many use cases, this has several advantages: The RT can be inferred from reference sequences that are longer than typical metagenomic reads used for OTU-based analysis, such as the 16S or 18S regions of the genome; hence, phylogenetic inference will be more reliable. Furthermore, the size of the RT can be chosen as needed, for example via our Phylogenetic Automatic (Reference) Tree (PhAT) method [34], instead of having to use the number of OTUs that result from the clustering and preprocessing steps. This also eliminates the need for the (mostly arbitrary) OTU cutoff step that is common to many metagenomic analyses, where OTUs with low abundance or low spread across samples are filtered out in order to keep the number of OTUs manageable. That is, with our approach, all sequences in a dataset can be placed and analyzed. Another advantage of a fixed RT is the availability of taxonomic annotation for the reference sequences. Often, in metabarcoding studies, the environmental sequences are anonymous and might not be closely related to any known species [11,15,32], which can hinder common taxonomic assignment methods [21].

Placing the sequences onto an RT with known taxonomic labels allows to easily interpret results within a given taxonomic framework. Using a taxonomically constrained RT can further improve interpretability. Lastly, using a fixed reference tree better allows to conduct cross- or meta-studies that compare samples from different sources, or to easily run analyses for samples that were added to the dataset later on. Using a fixed tree means that the context of interpretation remains unaltered. This is not easily possible with trees inferred from OTUs, as those change depending on the input sequences.

For further details on Phylofactorization, in particular the mathematical properties of the method, we refer to [92], which also covers different objective functions, elaborates on stopping criteria, and compares the method to other phylogenetic methods for analyzing ecological data. Compared to other tools and methods that use the phylogeny as a guide for analyzing microbial data, both, the original Phylofactorization as well as our adaptation allow for a direct interpretation of the results in terms of the edges of the tree, while avoiding nested dependencies between overlapping subtrees and circumventing issues associated with the compositional nature of the data.

### 3.5 Methods summary

We presented several novel methods for analyzing phylogenetic placement data. The methods are complementary, as each of them can identify different aspects and patterns in the underlying metagenomic samples. They are hence best used in combination, thereby enabling a thorough and comprehensive analysis and interpretation of the results. We also presented two transformations of phylogenetic placements (imbalances and balances) that shift the focus of the methods from a per-branch view of the data to a per-clade view.

In the following evaluation, we exemplify how the results obtained from the distinct methods can be interpreted in light of each other. Furthermore, we discuss strengths, weaknesses, appropriateness, and limitations of the methods and transformations.

## 4 Results and discussion

We used three real world datasets to evaluate our methods:

- Bacterial Vaginosis (BV) [18]. This small dataset was already analyzed with phylogenetic placement in the original publication. We used it as an example of an established study to compare our results to. It has 220 samples with a total of 15 060 unique sequences. See also [29] for a detailed analysis of this dataset which compares standard methods such as UniFrac [57,58] to methods like Edge PCA and Squash Clustering, which are based on phylogenetic placement.
- Tara Oceans (TO) [11,32,33]. This world-wide sequencing effort of the open oceans provides a rich set of meta-data, such as geographic location, temperature, and salinity. Unfortunately, the sample analysis for creating the official data repository is still ongoing. We thus were only able to use 370 samples with 27 697 007 unique sequences.
- Human Microbiome Project (HMP) [16,17]. This large data repository intends to characterize the human microbiota. It contains 9192 samples from different body sites with a total of 63 221 538 unique sequences. There is additional meta-data such as age and medical history, which is available upon special request. We only used the publicly available meta-data. See S1 Table for an overview of the dataset.

Details of the datasets (download links, data statistics, data preprocessing, etc.) are provided in S1 Text. At the time of writing, about one year after we initially downloaded the data, the TO repository has grown to 1170 samples, while the HMP even published a second phase and now comprises 23 666 samples of the 16S region. This further emphasizes the need for scalable data analysis methods.

Our test datasets represent a wide range of environments, number of samples, and sequence lengths. We use them to evaluate our methods and to exemplify which method is applicable to what kind of data. To this end, the sequences of the datasets were placed on appropriate phylogenetic RTs as explained in S1 Text, in order to obtain phylogenetic placements that our methods can be applied to. In the following, we present the respective results, and also compare our methods to other methods where applicable. As the amount and type of available meta-data differs for each dataset, we could not apply all methods to all datasets. Lastly, we also report the run-time performance of our methods on these data.

### 4.1 Visualization

#### 4.1.1 BV dataset

We re-analyzed the Bacterial Vaginosis (BV) dataset by inferring a tree from the original reference sequence set and conducting phylogenetic placement of the 220 samples. Bacterial Vaginosis is a disease of the vagina that manifests itself in form of an abnormal vaginal microbiome [18]. The characteristics of this dataset were already thoroughly explored in [18] and [29]. We use it here to give exemplary interpretations of our Edge Dispersion and Edge Correlation methods, and to evaluate them in comparison to existing methods. An overview of the placement result of this dataset is given in S1 Fig, where we show a tree and a heat map that indicate abundances/placement masses per edge and per sample.

Fig 4 shows our novel visualizations of the BV dataset. Edge Dispersion is shown in Fig 4(a), while Fig 4(b) shows Edge Correlation with the so-called Nugent score. The Nugent score [97] is a clinical standard for the diagnosis of Bacterial Vaginosis, ranging from 0 (healthy) to 10 (severe illness).

The connection between the Nugent score and the abundance of placements on particular edges was already explored in [29], but only visualized indirectly (i.e., not on the RT itself). For example, Figure 6 of the original study plots the first two Edge PCA components colorized by the Nugent score. We recalculated this figure for comparison in S7(i) Fig. In contrast, our Edge Correlation measure directly reveals the connection between Nugent score and placements on the reference tree: The clade on the left hand side of the tree in Fig 4(b), to which the red and orange branches lead to, are *Lactobacillus iners* and *Lactobacillus crispatus*, respectively, which were identified in [18] to be associated with a healthy vaginal microbiome. Thus, their presence in a sample is anti-correlated with the Nugent score, which is lower for healthy subjects. The branches leading to this clade are hence colored in red. On the other hand, there is a multitude of different other clades that exhibit a positive correlation with the Nugent score, that is, were green and blue paths lead to in the figure, again a finding already reported in [18].

Both trees in Fig 4 highlight the same parts of the tree: The dark branches with high deviation in Fig 4(a) represent clades attached to either highly correlated (blue) or anti-correlated (red) paths Fig 4(b). This indicates that edges that have a high dispersion also vary between samples of different Nugent score.

We further compared our methods to the visualization of Edge PCA components on the reference tree. To this end, we recalculated Figures 4 and 5 of [29], and visualized them with our color scheme in S5 Fig for ease of comparison. They show the first two components of Edge PCA, mapped back to the RT. The first component reveals that the *Lactobacillus* clade represents the axis with the highest heterogeneity across samples, while the second component further distinguishes between the two aforementioned clades within *Lactobacillus*. Edge Correlation also highlights the *Lactobacillus* clade as shown in Fig 4(b), but does not distinguish further between its sub-clades. This is because a high Nugent score is associated with a high abundance of placements in either of the two relevant *Lactobacillus* clades.

Further examples of variants of Edge Dispersion and Edge Correlation on this dataset are shown in S2 Fig and S3 Fig. We also conducted Edge Correlation using Amsel’s criteria [98] and the vaginal pH value as shown in S4 Fig, both of which were used in [18] as additional indicators of Bacterial Vaginosis. We again found similar correlations compared to the Nugent score.

#### 4.1.2 Tara Oceans dataset

We analyzed the Tara Oceans (TO) dataset to provide further exemplary use cases for our visualization methods. To this end, we used the unconstrained *Eukaryota* RT with 2059 taxa as provided by our Phylogenetic Automatic (Reference) Tree (PhAT) method [34]. The meta-data features of this dataset that best lend themselves to our methods are the sensor values for chlorophyll, nitrate, and oxygen concentration, as well as the salinity and temperature of the water samples. Other available meta-data features such as longitude and latitude are available, but would require more involved methods. This is because geographical coordinates yield pairwise distances between samples, whose integration into our correlation analysis methods is challenging. The Edge Correlation of the 370 samples with the nitrate concentration, the salinity, the chlorophyll concentration, and the water temperature are shown in S6 Fig.

We selected the diatoms and the animals as two exemplary clades for closer examination of the results. In particular, the diatoms show a high correlation with the nitrate concentration, as well as an anti-correlation with salinity, which represent well-known relationships [99,100]. See S6 Fig for details. These findings indicate that the method is able to identify known relationships. It will therefore also be useful to investigate or discover novel relationships between sequence abundances and environmental parameters.

#### 4.1.3 Performance

Both methods (Edge Dispersion and Edge Correlation) are computationally inexpensive, and thus applicable to large datasets. The calculation of the above visualizations took about 30 s each, which were mainly required for reading in the data. Furthermore, in order to scale to large datasets, we reimplemented Edge PCA, which was originally implemented as a command in the guppy program [22]. For the BV dataset with 220 samples, guppy required 9 min and used 2.2 GB of memory, while our implementation only required 33 s on a single core, using less than 600 MB of main memory. For the HMP dataset, as it is only single-threaded, guppy took 11 days and 75.1 GB memory, while our implementation needed 7.5 min on 16 cores and used 43.5 GB memory.

### 4.2 Clustering

We here evaluate our Phylogenetic *k*-means clustering (which uses edge masses and KR distances) and Imbalance *k*-means clustering (which uses edge imbalances and euclidean distances) methods in terms of their clustering accuracy. We used the BV as an example of a small dataset to which methods such as Squash Clustering [29] are still applicable, and the HMP dataset to showcase that our methods scale to datasets that are too large for existing methods.

#### 4.2.1 BV dataset

We again use the re-analyzed BV dataset to test whether our methods work as expected, by comparing them to the existing analysis of the data in [18] and [29]. To this end, we ran both Phylogenetic *k*-means and Imbalance *k*-means on the BV dataset. We chose *k* := 3, inspired by the findings of [18]. They distinguish between subjects affected by Bacterial Vaginosis and healthy subjects, and further separate the healthy ones into two categories depending on the dominating clade in the vaginal microbiome, which is either *Lactobacillus iners* or *Lactobacillus crispatus*. Any choice of *k* > 3 would simply result in smaller, more fine-grained clusters, but not change the general findings of these experiments. An evaluation of the number of clusters using the Elbow method is shown in S10 Fig. We furthermore conducted Squash Clustering and Edge PCA on the dataset, thereby reproducing previous results, in order to allow for a direct comparison between the methods, see Fig 8. Also, see [29] for a detailed interpretation of the results of Squash Clustering and Edge PCA on this dataset. The figure shows the results of Squash Clustering, Edge PCA, and two alternative dimensionality reduction methods, colorized by the cluster assignments *PKM* of Phylogenetic *k*-means (in red, green, and blue) and *IKM* of the Imbalance *k*-means (in purple, orange, and gray), respectively. We use two different color sets for the two methods, in order to make them distinguishable at first glance. Note that the mapping of colors to clusters is arbitrary and depends on the random initialization of the algorithm.

**Fig 8.**
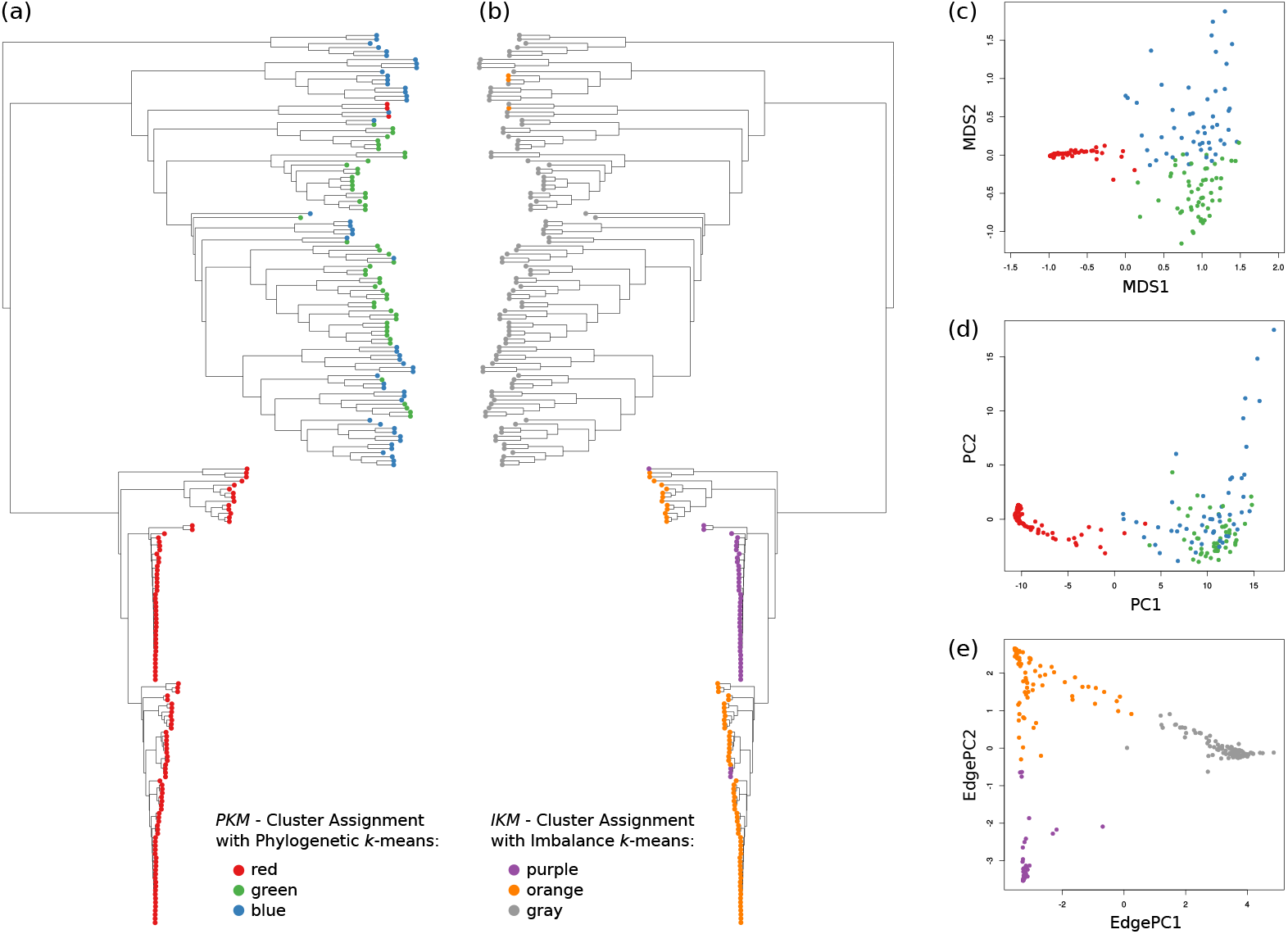
Comparison of *k*-means clustering to Squash Clustering and Edge PCA. We applied our variants of the *k*-means clustering method to the Bacterial Vaginosis (BV) dataset in order to compare them to existing methods. See [18] for details of the dataset and its interpretation. We chose *k* := 3, as this best fits the features of the dataset. For each sample, we obtained two cluster assignments: First, by using Phylogenetic *k*-means, we obtained a cluster assignment, which we here abbreviate as *PKM*. Second, by using Imbalance *k*-means, we obtained an assignment here abbreviated as *IKM*. In each subfigure, the 220 samples are represented by colored circles: red, green, and blue show the cluster assignments *PKM*, while purple, orange, and gray show the cluster assignments *IKM*. (a) Hierarchical cluster tree of the samples, using Squash Clustering. The tree is a recalculation of Figure 1(A) of Srinivasan *et al*. [18]. Each leaf represents a sample; branch lengths are Kantorovich-Rubinstein (KR) distances between the samples. We added color coding for the samples, using *PKM*. The lower half of red samples are mostly healthy subjects, while the green and blue upper half are patients affected by Bacterial Vaginosis. (b) The same tree, but annotated by *IKM*. The tree is flipped horizontally for ease of comparison. The healthy subjects are split into two sub-classes, discriminated by the dominating species in their vaginal microbiome: orange and purple represent samples were *Lactobacillus iners* and *Lactobacillus crispatus* dominate the microbiome, respectively. The patients mostly affected by BV are clustered in gray. (c) Multidimensional scaling using the pairwise KR distance matrix of the samples, and colored by *PKM*. (d) Principal component analysis (PCA) applied to the distance matrix by interpreting it as a data matrix. This is a recalculation of Figure 4 of [101], but colored by *PKM*. (e) Edge PCA applied to the samples, which is a recalculation of Figure 3 of Matsen *et al*. [101], but colored by *IKM*.

As can be seen in Fig 8(a), Squash Clustering as well as Phylogenetic *k*-means can distinguish healthy subjects from those affected by Bacterial Vaginosis. Healthy subjects constitute the lower part of the cluster tree. They have shorter branches between each other, indicating the smaller KR distance between them, which is a result of the dominance of *Lactobacillus* in healthy subjects. The same clusters are found by Phylogenetic *k*-means: As it uses the KR distance, it assigns all healthy subjects with short cluster tree branches to one cluster (shown in red). The green and blue clusters are mostly the subjects affected by the disease.

The distinguishing features between the green and the blue cluster are not apparent in the Squash cluster tree. This can however be seen in Fig 8(c), which shows a Multidimensional scaling (MDS) plot of the pairwise KR distances between the samples. MDS [64,102,103] is a dimensionality reduction method that can be used for visualizing levels of similarity between data points. Given a pairwise distance matrix, it finds an embedding into lower dimensions (in this case, 2 dimensions) that preserves higher dimensional distances as well as possible. Here, the red cluster forms a dense region, which is in agreement with its short branch lengths in the cluster tree. At the same time, the green and blue cluster are separated in the MDS plot, but form a coherent region of low density, indicating that *k* := 3 might be too large with Phylogenetic *k*-means on this dataset. That is, the actual clustering just distinguishes two clusters: healthy and sick patients (S10 Fig).

A similar visualization of the pairwise KR distances is shown in Fig 8(d). It is a recalculation of Figure 4 in the preprint [101], which did not appear in the final published version [29]. The figure shows a standard Principal Components Analysis (PCA) [64,103] applied to the distance matrix by interpreting it as a data matrix, and was previously used to motivate Edge PCA. However, although it is mathematically sound, the direct application of PCA to a distance matrix lacks a simple interpretation. Again, the red cluster is clearly separated from the rest, but this time, the distinction between the green and the blue cluster is not as apparent.

In Fig 8(b), we compare Squash Clustering to Imbalance *k*-means. Here, the distinction between the two *Lactobacillus* clades can be seen by the purple and orange cluster assignments. The cluster tree also separates those clusters into clades. The separate small group of orange samples above the purple clade is an artifact of the tree ladderization. The diseased subjects are all assigned to the gray cluster, represented by the upper half of the cluster tree. It is apparent that both methods separate the same samples from each other.

Lastly, Fig 8(e) compares Imbalance *k*-means to Edge PCA. The plot is a recalculation of Figure 3 of [101], which also appeared in Figure 6 in [29] and Figure 3 in [18], but colored using our cluster assignments. Because both methods work on edge imbalances, they group the data in the same way, that is, they clearly separate the two healthy groups and the diseased one from each other. Edge PCA forms a plot with three corners, which are colored by the three Imbalance *k*-means cluster assignments.

In S7 Fig, we report more details of the comparison of our *k*-means variants to the dimensionality reduction methods used here. Furthermore, exemplary visualizations of the cluster centroids are shown in S8 Fig, which further supports that our methods yield results that are in agreement with existing methods.

#### 4.2.2 HMP dataset

The HMP dataset is used here as an example to show that our method scales to large datasets. To this end, we used the unconstrained *Bacteria* RT with 1914 taxa as provided by our Phylogenetic Automatic (Reference) Tree (PhAT) method [34]. The tree represents a broad taxonomic range of *Bacteria*, that is, the sequences were not explicitly selected for the HMP dataset, in order to test the robustness of our clustering methods. We then placed the 9192 samples of the HMP dataset with a total of 118 701818 sequences on that tree, and calculated Phylogenetic and Imbalance *k*-means on the samples. The freely available meta-data for the HMP dataset labels each sample by the body site were it was taken from. As there are 18 different body site labels, we used *k* := 18. The result is shown in Fig 9. Furthermore, in S9 Fig, we show a clustering of this dataset into *k* := 8 broader body site regions to exemplify the effects of using different values of *k*. This is further explored by using the Elbow method as shown in S10 Fig.

**Fig 9.**
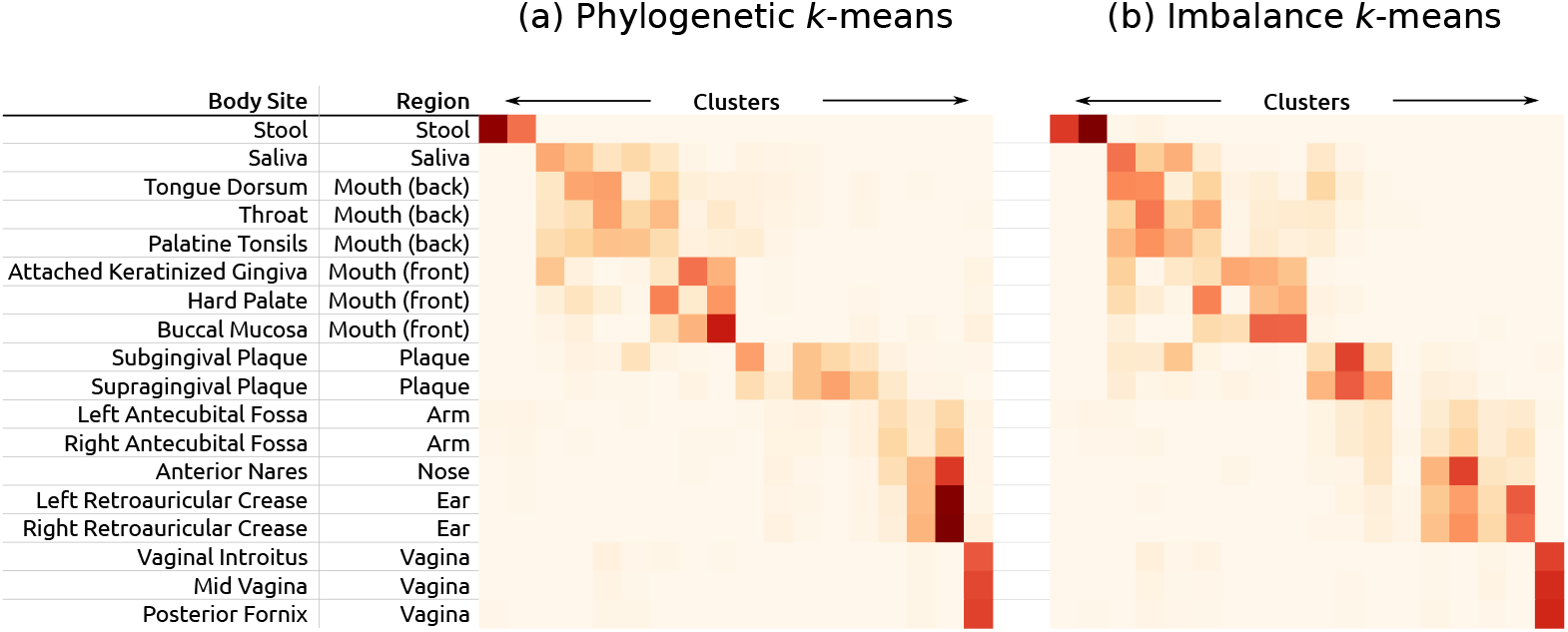
*k*-means cluster assignments of the HMP dataset with *k* := 18. Fig 9. Here, we show the cluster assignments as yielded by Phylogenetic *k*-means (a) and Imbalance *k*-means (b) of the Human Microbiome Project (HMP) dataset. We used *k* := 18, which is the number of body site labels in the dataset, in order to compare the clusterings to this “ground truth”. Each row represents a body site; each column one of the 18 clusters found by the algorithm. The color values indicate how many samples of a body site were assigned to each cluster. Similar body sites are clearly grouped together in coherent blocks, indicated by darker colors. For example, the stool samples were split into two clusters (topmost row), while the three vaginal sites were all put into one cluster (rightmost column). However, the algorithm cannot always distinguish between nearby sites, as can be seen from the fuzziness of the clusters of oral samples. This might be caused by our broad reference tree, and could potentially be resolved by using a tree more specialized for the data/region (not tested). Lastly, the figure also lists how the body site labels were aggregated into regions as used in S9 Fig. Although the plots of the two *k*-means variants generally exhibit similar characteristics, there are some differences. For example, the samples from the body surface (ear, nose, arm) form two relatively dense clusters (columns) in (a), whereas those sites are spread across four of five clusters in (b). On the other hand, the mouth samples are more densely clustered in (b).

Ideally, all samples from one body site would be assigned to the same cluster, hence forming a diagonal on the plot. However, as there are several nearby body sites, which share a large fraction of their microbiome [16], we do not expect a perfect clustering. Furthermore, we used a broad reference tree that might not be able to resolve details in some clades. Nonetheless, the clustering is reasonable, which indicates a robustness against the exact choice of reference taxa, and can thus by used for distinguishing among samples. For example, stool and vaginal samples are clearly clustered. Furthermore, the sites that are on the surface of the body (ear, nose, and arm) also mostly form two blocks of cluster columns.

#### 4.2.3 Performance

The complexity of Phylogenetic *k*-means is in 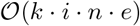, with *k* clusters, *i* iterations, and *n* samples, and *e* being the number of tree edges, which corresponds to the number of dimensions in standard euclidean *k*-means. As the centroids are randomly initialized, the number of iterations can vary; in our tests, it was mostly below 100. For the BV dataset with 220 samples and a reference tree with 1590 edges, using *k* := 3, our implementation ran 9 iterations, needing 35 s and 730 MB of main memory on a single core. For the HMP dataset with 9192 samples containing 119 million sequences, and a reference tree with 3824 edges, we used *k* := 18, which took 46 iterations and ran in 2.7 h on 16 cores, using 48 GB memory.

In contrast to this, Imbalance *k*-means does not need to conduct any expensive tree traversals, nor take single placement locations into account, but instead operates on compact vectors with one entry per edge, using euclidean distances. It is hence several orders of magnitude faster than Phylogenetic *k*-means, and only needs a fraction of the memory. For example, using again *k* := 18 for the HMP dataset, the algorithm executed 75 iterations in 2 s. It is thus also applicable to extremely large datasets.

Furthermore, as the KR distance is used in Phylogenetic *k*-means as well as other methods such as Squash Clustering, our implementation is highly optimized and outperforms the existing implementation in guppy [22] by orders of magnitude (see below for details). The KR distance between two samples has a linear computational complexity in both the number of QSs and the tree size. As a test case, we computed the pairwise distance matrix between sets of samples. Calculating this matrix is quadratic in the number of samples, and is thus expensive for large datasets. For example, in order to calculate the matrix for the BV dataset with 220 samples, guppy can only use a single core and required 86 min. Our KR distance implementation in genesis is faster and also supports multiple cores. It only needed 90 s on a single core; almost half of this time is used for reading input files. When using 32 cores, the matrix calculation itself only took 8 s. This allows to process larger datasets: The distance matrix of the HMP dataset with 9192 samples placed on a tree with 3824 branches for instance took less than 10 h to calculate using 16 cores in genesis. In contrast, guppy needed 43 days for this dataset. Lastly, branch binning can be used to achieve additional speedups, as shown in S2 Table.

### 4.3 Phylogenetic ILR Transform and Balances

As a first test of our adaptation of balances to placement data, we apply it to the BV dataset. Further assessment of balances for placement data, also with the HMP dataset, is implicitly conducted by the evaluation of Placement-Factorization below, which uses balances for aggregation and contrasting of subtrees.

As balances are conceptually similar to edge imbalances, we perform analogous evaluations. To this end, we computed the per-edge balance for all edges of the BV reference tree, across all 220 samples. That is, for each edge, we computed the balance between the two subtrees induced by the edge. This yields a matrix that we call *balance matrix*, and which corresponds to the imbalance matrix used for Edge PCA, c. f. Fig 3(b). Hence, a natural first visualization of the balances is to analyze their principal components, that is, to compute the PCA of the balance matrix. The first two components are shown in S11 Fig, for both variants of the balance computation (with and without taxon weighting). The components exhibit a separation of the samples by Nugent score, showing that they yield results comparable to Edge PCA.

In order to interpret what the axes of these principal components mean, we can again employ the visualization of PCA eigenvectors on the reference tree as used in Edge PCA [29], c. f., S5 Fig. We show the results for PCA of the balances in S12 Fig. As with Edge PCA, the principal components correspond to the *Lactobacillus* clade, with the first component mostly separating *Lactobacillus* from the rest of the tree, and the second component further distinguishing between *Lactobacillus crispatus* and *Lactobacillus iners*.

Furthermore, as mentioned in the methods description, balances could in principle be used as input to our previously presented methods, such as Edge Correlation and *k*-means clustering (which adequately might be called Balance *k*-means), in the same manner that we used imbalances before. We exemplify the correlation of the Nugent score with balances in S13 Fig. However, artifacts might arise from the underlying mathematical framework of balances, in particular the usage of the geometric mean without taxon weighting: The geometric mean is *not* sensitive to singular large values, such as the high amount of placement mass on one of the *Lactobacillus* branches. It only significantly increases if multiple high values are present, such as the multitude of bacterial taxa with high abundance in diseased patients of the BV dataset [18]. This can lead to spurious results, as shown in S13(b) Fig, where the correlation of the unweighted balances with the Nugent score yields unrealistically high negative correlations for almost all branches that have little placement mass on them. Moreover, this property of the geometric mean implies that it is sensitive to taxa splitting (pers. comm. with A. Washburne on 2018-11-23): For example, it *does* make a difference whether masses are focused on a single branch, or distributed across several representatives of the same species.

Hence, we do not recommend to use (unweighted) balances for computations such as correlations or *k*-means clustering. Note that when used with GLMs, such as in Phylofactorization, these issues do not arise: The winning edge is chosen to maximize the difference between the null deviance and the deviance of the linear model. That difference is small for clades with almost no mass (such as the ones affected by the issue above), so that the value of the objective function for such edges is lower than for edges with more mass. Hence, the factorization does not incorrectly identify these low-abundance clades as potential factors. The usage of balances in Placement-Factorization is further explored in the next section.

### 4.4 Placement-Factorization

In the following we present results from *Placement-Factorization*, and compare them to the above results from our other methods, as well as to the original Phylofactorization. For comparability with the original method, we solely use balances for aggregating and contrasting, and an objective function that maximizes the difference between the null deviance and the deviance obtained by a Generalized Linear Model (GLM). Other choices of functions for Phylofactorization have been explored in [31] and [92], see there for details. The exploration of their effect on Placement-Factorization is left as future work, although based on the consistency of our results with the original method, we conjecture that they behave according to the findings of the original publications.

Furthermore, we note that our implementation supports taxon weighting as introduced in the Phylogenetic ILR transform [30], which is however not (yet) supported by the original Phylofactorization [31]. We found this weighting scheme to be a natural and valuable addition in the balances computation that yields results closer to those obtained with imbalances. We suspect that this is because the weighting scheme can alleviate the issues of the geometric mean mentioned above.

#### 4.4.1 BV dataset

We analyzed the BV dataset [18] with our Placement-Factorization with and without taxon weighting, using balances for aggregating and contrasting, and GLMs for the objective function. As GLMs support multiple predictors at the same time, we used all three available meta-data features of the dataset simultaneously for the regression, that is, Nugent score [97], Amsel’s criteria [98], and the pH-value of the samples. We also tested with only the Nugent score to be consistent with our previous analyses, and to asses the robustness of the method with respect to the specific choice of meta-data features. We observe only minor differences in the ordering of the identified factors, that is, which clades were “winning” in which iteration. Hence, we focus on the results obtained with all three meta-data features taken into account.

For comparing with the original method, we clustered the dataset into OTUs using two different OTU clustering methods, vsearch [54] and swarm [52,53], and inferred two trees from these OTU clusterings. We used two distinct OTU clustering methods to asses how they affect factorization; see S1 Text for details on the preprocessing steps. We then conducted an analysis of both trees with the original Phylofactorization, again using balances and GLMs. We compare the results of Placement-Factorization to our previous analyses of the data as well as to the original Phylofactorization on the two alternative OTU trees.

In S3 Table, we compare the clades found by the two Phylofactorization variants (with vsearch and with swarm) to the clades found by Placement-Factorization *without* taxon weighting. Moreover, in S14 Fig we visualize the clades found by Placement-Factorization. These outcomes show that our results are consistent with the existing Phylofactorization, in that similar clades are split from the tree, albeit with some variation in the order by which clades are selected. The clades being split are also consistent with previous analyses of the dataset [18], as all taxa found by the first 10 factors of Placement-Factorization were also found to be relevant in the context of Bacterial Vaginosis in [18]. However, the vsearch-based Phylofactorization is the only evaluated variant that split the *Lactobacillus* clade in the first factor and further *Lactobacillus crispatus* from *Lactobacillus iners* in the second factor. The swarm-based variant and our Placement-Factorization without taxon weighting also identified these clades, but not in the first two iterations.

When using taxon weighting on the other hand, Placement-Factorization also finds these two clades in the first two factors, and is hence more consistent with existing analyses. However, due to small differences in the value of the objective function, the first iteration splits a larger clade than expected. We observed a similar behavior with the vsearch-based Phylofactorization; see the long list of taxa of the first factor in S3 Table. In order to identify the provenance of this effect and to correctly interpret the factors, we developed a novel visualization of the results: In Fig 10 and in S15 Fig, we show the reference tree, where each edge is colored by the value of the objective function at that edge. This type of visualization helps to assess the uncertainty involved in identifying the winning edge of a specific iteration.

**Fig 10.**
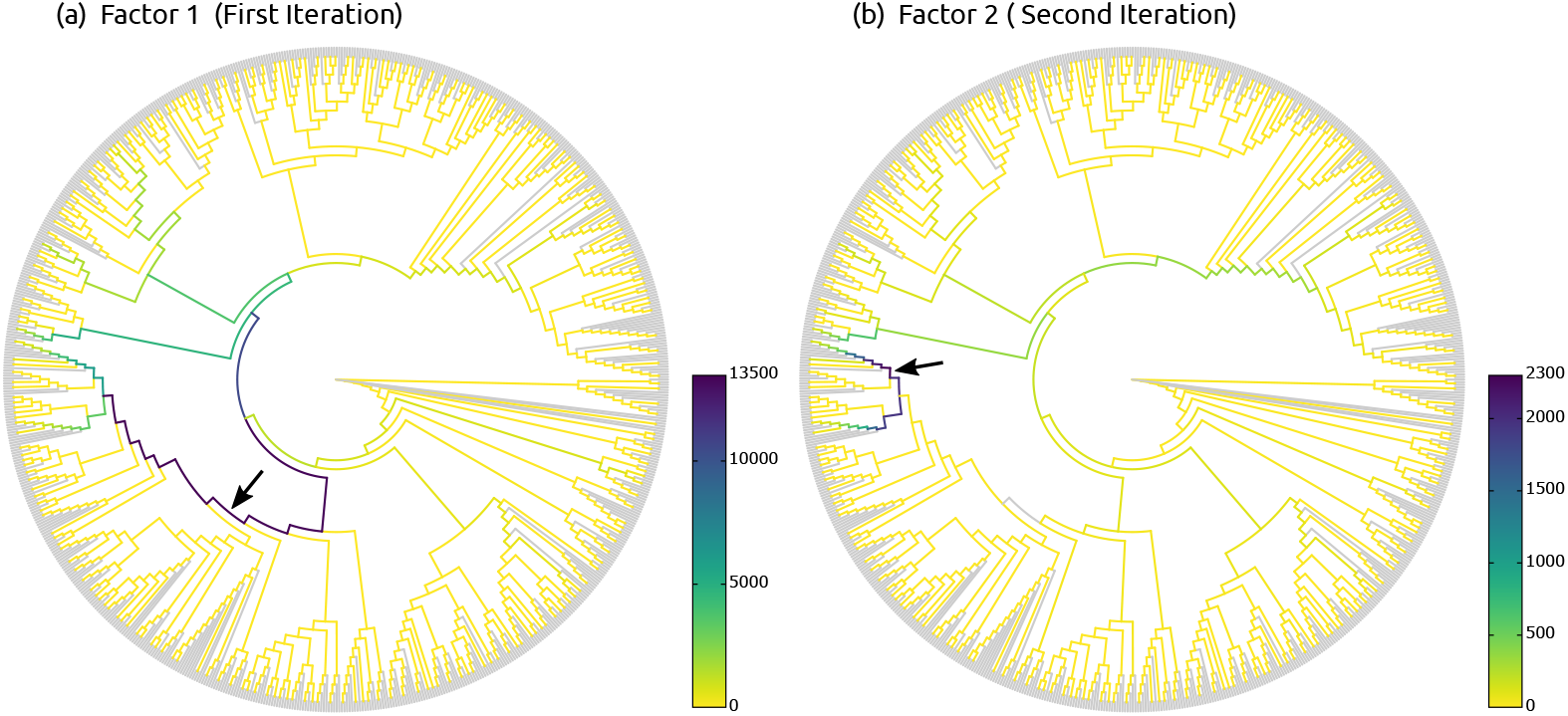
Objective values of Placement-Factorization of the BV dataset. Here, we show the values of the objective function for each inner edge, for the first two factors found by Placement-Factorization (with taxon weights) of the BV dataset. The winning edge of each iteration is marked by a black arrow. This novel visualization of phylogenetic factors helps to understand why a particular edge was chosen in an iteration: Here, the objective function of the first iteration in (a) yields high values for the path towards the *Lactobacillus* clade, consistent with previous findings. However, due to small differences, the winning edge of the first iteration is chosen to be relatively basal in the tree, meaning that a large clade is factored out. This obfuscates the fact that this factor is mostly concerning the *Lactobacillus* clade, and not so much the remaining taxa in that clade. This visualization thus aids interpretation of the found clades, and allows to identify the parts of a factored clade that are most relevant to the factor. In the second iteration in (b), the tree clearly shows the distinction between the two relevant clades of *Lactobacillus* again, consistent with previous findings. See also S15 Fig for the version of this visualization without taxon weights.

For example, the visualization in Fig 10(a) reveals that the large clade found by the first factor (marked with an arrow) does in fact include many branches and taxa with a low value of the objective function (yellow branches). These are branches with low placement mass that do not contribute much to this factor. Instead, there is a path of comparably high values of the objective function that leads down to the *Lactobacillus* clade. This indicates that there are several ‘good’ candidate edges for distinguishing patients by their health status, and that the smaller *Lactobacillus* clade is the actual clade of interest in this factor. The winning edge just happened to have a slightly higher value than other edges on this path. To address this issue, a proper statistical test of the significance of each winning edge compared to the other edges evaluated in the iteration could be employed. This is connected to the idea of *confidence regions* of each factor on the tree, as presented in [92], which we discuss later.

To further assess how the samples are split by individual factors, we used the balances at each iteration/factor as an ordination of the data, as suggested in [31], which we show in S16 Fig. These plots reveal that the splitting into healthy vs. diseased patients works both with and without taxon weighting, albeit the differences in the respective plot shapes are pronounced. As scatter plots of balances can only reasonably reveal the first two or three factors, S16 Fig serves as a caveat for this type of visualization: The BV dataset does indeed have two important features concerning the healthy patients, namely the *Lactobacillus* clade, and the further distinction into *Lactobacillus crispatus* and *Lactobacillus iners*. However, as discussed above and visible in S1 Fig, the diseased patients exhibit high abundances in a multitude of other clades, which cannot be expressed by just two or three factors. We later show a novel way for visualizing balances in the HMP dataset that can help to understand the balances of all factors.

In summary, we find that Placement-Factorization *without* taxon weighting behaves similar to the original Phylofactorization (which also does not employ a taxon weighting scheme), while Placement-Factorization *with* taxon weighting yields results that are more in line with our previous results based on imbalances. The latter is likely because taxon weighting has a similar effect of reducing the influence of low mass branches (low abundance taxa) as the summation-based aggregation step of imbalances.

#### 4.4.2 HMP dataset

The original publication of Phylofactorization used a dataset comprising oral and fecal samples from the human microbiome as one of their case studies [104], see Figure 4 and supplementary figures S3–S8 of [31]. For our comparison, we selected a suitable subset of the HMP dataset [16,17]: In particular, we selected all 600 stool samples of the dataset, as well as 600 randomly chosen samples from the mouth region, that is, from the samples labeled “Mouth (back)” and “Mouth (front)” in S1 Table. We again used the placement of these samples on the unconstrained *Bacteria* tree of our Phylogenetic Automatic (Reference) Tree (PhAT) method [34], containing 1914 taxa, to conduct Placement-Factorization. We henceforth assume that the oral/fecal dataset of [104] and our oral/fecal subset of the HMP dataset exhibit comparable sequence compositions. Furthermore, as the tree used for Phylofactorization in [31] is based on the OTUs of the sequences, it only contains taxa that are sufficiently abundant in the input. It thus differs from the more general reference tree used for our evaluation here. Therefore, we had to map the taxa found by Phylofactorization to the underlying Silva taxonomy [105,106] that was used for constructing our tree.

Despite these differences, Placement-Factorization yielded factors that are similar to the ones found by Phylofactorization. We compare the taxa identified by the first 10 factors of each variant. For simplicity, we only compare the clades on the non-root side of the (arbitrarily rooted) reference tree; the paraphyletic “remainder” clade is not taken into account. Furthermore, we do not consider the order of the factors here. Similar to the findings of the BV dataset above, Placement-Factorization *with* taxon weighting yielded larger clades than *without* taxon weighting, which again yielded larger clades than the OTU-based Phylofactorization. The latter is a consequence of the OTU tree containing fewer taxa than our broad *Bacteria* tree. We found that 84% of the taxa identified by Phylofactorization were also part of the factors of our variants, with the major difference being a set of *Proteobacteria* that were part of the split in the first factor of Phylofactorization [31], but not by our variants. This is most likely an artifact of the differing trees being used in the factorization. Furthermore, 95% of the taxa found by Placement-Factorization *without* taxon weighting were also part of the clades *with* taxon weighting. We visualized the clades found by all three variants in S17 Fig, which also provides further details on the comparison. Most of the factors found by the three variants agree with each other, with their disagreement mostly concerning the clade sizes. The actual differences in taxa (such as the *Proteobacteria* not found by our variants) serve as a caveat for the importance of the underlying reference tree: Differences in topology will inevitably be reflected in different factors, which might in turn suggest a different interpretation of results. In an ideal world with a known phylogeny of all of life, alternative OTU clusterings and alternative trees would simply collapse nodes at different depths (pers. comm. with A. Washburne on 2019-03-01). Unfortunately, real world data, and particularly different OTU clustering methods and tree inference methods, will yield discordant trees. The influence of uncertainty in the phylogeny is further discussed in [92].

Next, we investigated how well the factors found by Placement-Factorization separate oral from fecal samples. To this end, we again employed the balances of the winning edge of each factor for an ordination visualization [31], which we show in S18(a) Fig and S18(b) Fig. The ordination clearly separates the samples, both with and without taxon weighting. Again, ordination scatter plots can only reveal up to three dimensions/factors. In order to evaluate the separation of samples at later factors, we use a visualization of the factor balances, which we call *balance swarm plots*, and which are similar to the per-factor ordination plots used in [92]. These plots can show the ordination of arbitrarily many factors at the same time, as shown in Fig 11(a), as well as S18(c) Fig and S18(d) Fig.

**Fig 11.**
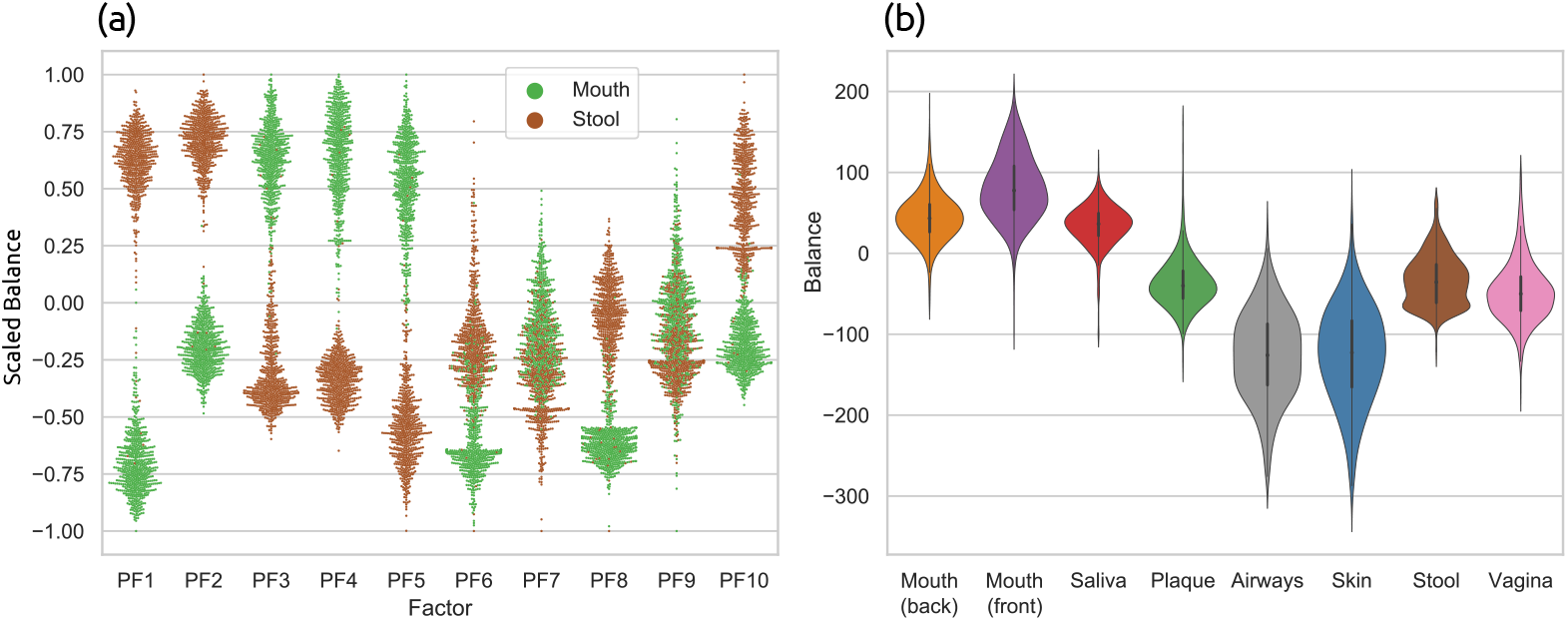
Ordination of Placement-Factorization of the HMP dataset. In this visualization of phylogenetic factors, we show the balances of the winning edge at different factors for all samples. Subfigure (a) shows the first 10 factors found by Placement-Factorization with taxon weighting on the oral/fecal subset of the HMP dataset. We call this a *balance swarm plot*. It can be understood as multi-dimensional scatter plot, where each dimension is shown separately: Each column corresponds to a factor (PF1-PF10), with the vertical axis being the balances, and horizontal space within each column used to spread samples at nearby positions, revealing their distribution density. The balances were scaled to the [−1.0,1.0] interval for better comparability across factors, while keeping the centering at 0. Subfigure (b) shows the first factor of Placement-Factorization with taxon weighting on the full HMP dataset. The violin plots in (b) extend on the idea of balance swarm plots by separating different groups of samples, based on their body site. This allows to clearly see the distribution of balances at the factor for all groups of samples. The exhaustive versions of these plots, with and without taxon weighting, and for more factors, are shown in the context of the typical two- and three-dimensional scatter plots in S18 Fig, S19 Fig, and S20 Fig. See there for more details.

As shown in Fig 11(a) and S18(c) Fig, eight out of the first ten factors found by Placement-Factorization *with* taxon weighting clearly separate the oral from the fecal samples. The remaining two factors (PF7 and PF9) separate most of the samples, but also have an interval of balances that contains samples from both body sites. Placement-Factorization *without* taxon weighting also separates samples based on their body site, as shown in S18(d) Fig, but with a less clear distinction. This is also obvious from the ordination scatter plots shown in S18(b) Fig.

Finally, we conducted Placement-Factorization on the whole HMP dataset with all 9192 samples, instead of just the oral/fecal subset, in order to evaluate how the method performs on large datasets with more than two categories (body sites) to distinguish. See S1 Table for an overview of the samples, as well as a list of the eight body site labels that we used for classifying the samples. We do not discuss the taxa that were split by each factor, as such an in-depth biological discussion is beyond the scope of this manuscript. Instead, we evaluate how well different body sites were separated by the factors. In S19 Fig, we show ordination plots of the first two and three factors. These plots already reveal that Placement-Factorization indeed separates samples from each other based on their body site. However, given the eight body site labels that we used, these plots are overloaded. Hence, we extended on the idea of balance swarm plots (as introduced above) by separating them into individual plots per factor, each showing the balance distribution of groups of samples. An example for the first factor is shown in Fig 11(b); we furthermore show the first four factors in S20 Fig. These visualizations indicate that Placement-Factorization separates samples mainly based on the distinction oral vs. remaining body sites, with a further separation of plaque samples in the oral region. This can, for example, be seen in Fig 11(b), where the first three groups “Mouth (back)”, “Mouth (front)”, and “Saliva” exhibit balances above 0, while all other groups have balances below 0. Further factors then separate vaginal samples and skin and airways samples from the rest of the samples, as shown in S20 Fig. Overall, Placement-Factorization can distinguish these samples by body site, at least to the extent that can expected from abundance differences in the samples. For example, it would be unrealistic to expect the algorithm to perfectly separate samples from the back and front of the mouth from each other.

In conclusion, Placement-Factorization yields factors of the oral/fecal data that are mostly consistent with the findings of Phylofactorization [31], and is also able to reasonably separate the samples of larger datasets with several categorical labels.

#### 4.4.3 Performance

The run time of Placement-Factorization depends on (a) the number of input samples, (b) the number of branches of the reference tree, and (c) the number of iterations to run. As the computations are conducted on the mass matrix instead of single placements, the performance and memory requirements of Placement-Factorization are independent from the total number of sequences/placements in the dataset. In each iteration, and for each edge of the tree (except the ones that won previous factors), the balances of all samples are computed, and the objective function is evaluated. In case of using a GLM to express the relationship of balances with meta-data, this involves fitting a model across all samples. In our implementation, all these computations are parallelized.

Our relatively small BV test dataset ran on a standard laptop with 4 cores, taking 30 s per iteration. The full HMP dataset with 9192 samples and our reference tree with 3825 branches required 13.0 GB of memory in our non-optimized prototype implementation, as took less than 90 s per iteration using 20 cores. Also, note that later iterations tend to become faster, as the splitting of the tree into subtrees reduces the number of edges that need to be taken into account in each balance computation. Hence, we conclude that Placement-Factorization is well suited even for very large datasets.

#### 4.4.4 Future directions

Phylofactorization is a very recent method whose full potential has just begun being explored [92]. We contributed novel ideas by (a) adapting the concept to phylogenetic placement, which can be thought of as placing abundances along branches of the tree instead of just at its tips; and (b) suggesting a novel visualization for objective function value at each edge of the reference tree, which helps in the interpretation of the factors being split in each iteration.

As discussed above, we found that both, the original Phylofactorization, as well as our Placement-Factorization, can split clades that are larger than one would expect from other types of analyses of the data. Considering the distribution of objective function values, as shown in Fig 10, it is likely that such large clades are the result of random variability along a path of branches that are equally relevant for the factor. Further research is needed to confirm this.

These findings suggest that it might be beneficial to introduce a significance value for each factor, which assesses how relevant the particular winning edge is compared to other edges that yielded a high objective value in an iteration. This idea is intrinsically connected (pers. comm. with A. Washburne on 2019-03-01) to the stopping function of the original Phylofactorization [31], which uses a Kolmogorov-Smirnov (KS) test to conservatively estimate when a sufficient number of factors have been identified. Another strongly connected idea is that of confidence regions of the phylogeny, defined by regions of the tree in which the “true” winning edge falls with a certain confidence [92]. Such a significance value for the winning edge might also enable a form of *soft* factorization, that does not greedily pick one winning edge per iteration.

Furthermore, the paths of high objective values as seen in Fig 10 indicate that there is a gradient of the objective function along the branches of the tree. This could be exploited in a gradient-ascending graph-walking algorithm to identify the phylogenetic factors of extremely large datasets without having to exhaustively evaluate the objective function at every edge (pers. comm. with A. Washburne on 2019-03-01).

## 5 Conclusion

We presented novel, scalable methods to analyze and visualize phylogenetic placements of metagenomic samples. Phylogenetic placement of the sequences on a fixed reference tree allows for an interpretation of the data in a phylogenetic context. The methods are computationally inexpensive, and are thus, as we have demonstrated, applicable to large datasets. They are built on top of a common set of concepts, and hence gear well into each other.

- *Edge Dispersion* highlights branches of the phylogenetic tree that exhibit variations in the number of placements, and thus allows to identify regions of the tree with a high placement heterogeneity. *Edge Correlation* additionally takes meta-data features into account, and identifies branches of the tree that correlate with quantitative features, such as the temperature or the pH value of the environmental samples. These methods complement existing methods such as Edge PCA [29], and represent data exploration tools that can help unravel new patterns in phylogenetic placement data, and hence, in metagenomic samples.
- We presented adapted variants of the *k*-means method, called *Phylogenetic *k*-means* and *Imbalance *k*-means*, which exploit the structure of phylogenetic placement data to identify clusters of environmental samples. The methods build upon ideas such as Squash Clustering [29] and can be applied to substantially larger datasets, as they use a pre-defined number of clusters. For future exploration, other forms of cluster analyses could be adapted to phylogenetic placement data, for example, soft *k*-means clustering [107,108] or density-based methods [109]. The main challenge when adopting such methods consists in making them phylogeny-aware, that is, to use mass distributions on trees instead of the typical ℝ^*n*^ vectors.
- We introduced an adaptation of the Phylogenetic ILR transform and balances [30] to phylogenetic placements. As balances are a transformation that yields orthogonal components (one for each node or branch of the tree), issues pertaining to the normalization of compositional data do not arise. With samples being represented as a vector of balances, numerous standard tools for data visualization, ordination, and clustering in the euclidean space can be readily applied to phylogenetic placement data. Applying these methods to placements instead of OTUs allows for more detailed analyses, as the entire original sequence data can be used. Furthermore, using a fixed reference tree instead of one inferred from the OTUs present in a set of samples enables comparative studies across datasets.
- Lastly, we presented an adaption of Phylofactorization [31,92], which we call *Placement-Factorization*. Placement-Factorization identifies branches of the reference tree, called *phylogenetic factors*, that exhibit a relationship with environmental meta-data features, that is, branches along which putative functional traits might have arisen in conjunction with changes in environmental variables. This factorization of the tree can be used as an ordination tool to visualize how samples are separated by changes along the factors, and as a dimensionality-reduction tool [31]. It thus complements Edge Correlation, but further allows to identify nested dependencies within sub-clades of the reference tree. We leave the adaptation of some of the original concepts of Phylofactorization to phylogenetic placements as future work, such as binned phylogenetic units (BPUs), stopping criteria for the iterations, as well as further experimentation with different objective functions and aggregation and contrast functions [31,91]. Based on our findings and experiments, we conjecture that these concepts should be readily applicable to our Placement-Factorization.

The presented methods take either the edge masses as input, that is, the abundances of metagenomic sequences on each branch of the reference tree, or a transformation of theses masses, such as edge imbalances or balances, and can hence analyze different aspects of the placements and the environmental samples. While edge masses reveal information about single branches, edge imbalances and balances take entire reference tree clades into account. Depending on the task at hand, either of them might be preferable and better suited for identifying patterns. We also elaborated on the limitations and caveats of these transformations. Furthermore, we again emphasize the importance of appropriately normalizing the samples as required, in order to cope with the compositional nature of metagenomic data. That is, depending on the type of sequence data, using either absolute or relative abundances is critical to allow for meaningful interpretation of the results.

We evaluated our novel methods on three real-world datasets (see S1 Text for details on the datasets and their preprocessing), and gave exemplary interpretations of the results. We further showed that these results are consistent with existing methods as well as empirical biological knowledge. Hence, our methods will also be useful to unravel new, unexplored relationships in metagenomic data. Each of them has their strengths and weaknesses, and focuses on different aspects of the data. The methods and their variants are hence best used in combination with each other, in order to obtain a thorough and comprehensive analysis.

The methods are implemented in our tool gappa, which is freely available under GPLv3 at http://github.com/lczech/gappa. Furthermore, gappa has recently been bundled into a bioconda package, which is available at https://anaconda.org/bioconda/gappa; note that this package is not maintained by ourselves. In S2 Text, we briefly describe the software and its commands, and also describe a typical phylogenetic placement analysis pipeline. The methods and transformations presented here constitute a toolbox of different techniques that can be combined with each other, and allow for further extension and experimentation. The implementation in gappa is mainly based on our library genesis, which is also freely available under GPLv3 at http://github.com/lczech/genesis. Furthermore, scripts, data and other tools used for the tests and figures presented here are available at http://github.com/lczech/placement-methods-paper.

## 6 Acknowledgments

We thank S. Srinivasan and E. Matsen for providing the Bacterial Vaginosis dataset [18] and for helping us understanding their methods and implementations. We further thank **A. Washburne** and **J. Silverman** for fruitful exchange about the underlying mathematics of their methods [30,31,91,92]. We also thank **M. Dunthorn, L. Rubinat-Ripoll, C. Berney, L. Guidi, G. Lentendu, A. Kozlov**, and **P. Barbera** for their feedback on our methods and help with this manuscript, and we thank **G. Douglas** for bundling gappa as a bioconda package. Lastly, we want to express our gratitude towards one of the anonymous reviewers of this manuscript for inspiring us to our adaptation of balances and phylofactorization.

## 7 Supporting information

### 8 S1 Text. Empirical datasets

The analyses and figures presented in the manuscript were conducted on distinct reference alignments and trees, suited for each dataset. Firstly, for the BV dataset, we used the set of reference sequences from the original study [18], and re-inferred a tree on them. Secondly, for the TO and HMP datasets, we used our Phylogenetic Automatic (Reference) Tree (PhAT) method [34] to construct sets of suitable reference sequences from the Silva database [105,106]. We used the 90% threshold consensus sequences; see [34] for details.

For all analyses, we used the following software setup: Unconstrained maximum likelihood trees were inferred using RAxML v8.2.8 [62]. For aligning reads against reference alignments and reference trees, we used a custom MPI wrapper for P_A_P_A_R_A_ 2.0 [35,36], which is available at [110]. We then applied the `chunkify` procedure as explained in [34] to split the sequences into chunks of unique sequences prior to conducting the phylogenetic placement, in order to minimize processing time. Phylogenetic placement was conducted using EPA-ng [24,111], which is a faster and more scalable phylogenetic placement implementation than RAxML-EPA [23] and pplacer [22]. Lastly, given the per-chunk placement files produced by EPA-ng, we executed the `unchunkify` procedure of [34] to obtain per-sample placement files. These subsequently served as the input data for the methods presented here.

We made the scripts, data and other tools used for the tests and figures presented here available at http://github.com/lczech/placement-methods-paper. See there for further details.

#### 8.1 Bacterial Vaginosis

We used the Bacterial Vaginosis dataset [18] in order to compare our novel methods to existing ones such as Edge PCA and Squash Clustering [29,56]. The dataset contains metabarcoding sequences of the vaginal microbiome of 220 women, and was kindly provided by Sujatha Srinivasan. This small dataset with a total of 426 612 query sequences, thereof 15 060 unique, was already analyzed with phylogenetic placement methods in the original publication [18] and in [29]. We re-inferred the reference tree of the original publication using the original alignment, which contains 797 reference sequences specifically selected to represent the vaginal microbiome. As the query sequences were already prepared, no further preprocessing was applied prior to phylogenetic placement. The available per-sample quantitative meta-data for this dataset comprises the Nugent score [97], the value of Amsel’s criteria [98], and the vaginal pH value. We used all three meta-data types in our analyses.

For our comparison of Placement-Factorization to the original Phylofactorization [31], we furthermore conducted OTU clustering of the sequences, using two different methods: We used vsearch v2.9.1 [54] as well as swarm v2.2.2 [52, 53] to obtain two sets of OTU clusters. We filtered the OTU table to remove low abundance OTUs, by only keeping those that appear in more than 10% of the samples. In order to assign each OTU to a fitting taxonomic path, we used the assign command of our toll gappa. To this end, we placed the OTUs on the BV reference tree mentioned above, in order to obtain taxonomic assignments for the OTUs that are in line with the taxonomic labels used in our other analyses of the dataset. Each set of OTUs was subsequently aligned with MAFFT v7.310 [112,113], using the L-INS-I strategy [114]. Finally, we inferred an OTU tree for each set, using the recent RAxML-NG v0.7.0 [115]. These two OTU trees were then used with the meta-data for conducting an analysis with Phylofactor, based on the excellent tutorials at https://github.com/reptalex/phylofactor. The results for the first ten factors for each of these two trees is for example shown in S3 Table of the supplement.

#### 8.2 Tara Oceans

The Tara Oceans (TO) dataset [11,32,33] that we used here contains amplicon sequences of protists, and is available at https://www.ebi.ac.uk/ena/data/view/PRJEB6610.

At the time of download, there were 370 samples available with a total of 49 023 231 sequences. As the available data are raw `fastq` files, we followed [116] to generate cleaned per-sample `fasta` files. For this, we used our tool PEAR [117] to merge the paired-end reads; cutadapt [118] for trimming tags as well as forward and reverse primers; and VSEARCH [54] for filtering erroneous sequences and generating per-sample `fasta` files. We filtered out sequences below 95 bps and above 150 bps, to remove potentially erroneous sequences. No further preprocessing (such as chimera detection) was applied. This resulted in a total of 48 036 019 sequences, thereof 27 697 007 unique. The sequences were then used for phylogenetic placement as explained above. We placed the sequences on the unconstrained *Eukaryota* reference tree obtained via our Phylogenetic Automatic (Reference) Trees (PhAT) method [34], which comprises 2059 taxa, thereof 1795 eukaryotic sequences. The remaining 264 taxa are *Archaea* and *Bacteria*, and were included as a broad outgroup. The TO dataset has a rich variety of per-sample meta-data features; in the context of this paper, we mainly focus on quantitative features such as temperature, salinity, as well as oxygen, nitrate and chlorophyll content of the water. Furthermore, each sample has meta-data features indicating the date, longitude and latitude, depth, etc. of the sampling location. This data might be interesting for further correlation analyses based on geographical information. We did not use them here, as for example longitude and latitude would require a more involved method that also accounts for, e.g., ocean currents. Furthermore, geographical coordinates yield pairwise distances between samples, which are not readily usable with our correlation analysis. Lastly, in order to use features such as the date, that is, in order to analyze samples over time, the same sampling locations would need to be visited at different times during the year, which is not the case for the Tara Oceans expedition.

#### 8.3 Human Microbiome Project

We used the Human Microbiome Project (HMP) dataset [16,17] for testing the scalability of our methods. In particular, we used the “HM16STR” data of the initial phase “HMP1”, which are available from http://www.hmpdacc.org/hmp/. The dataset consists of trimmed 16S rRNA sequences of the `V1V3, V3V5`, and `V6V9` regions. The data are further divided into a “by_sample” set and a “healthy” set, which we merged in order to obtain one large dataset, with a total of 9811 samples. We then removed 10 samples that were larger than 70 MB as well as 605 samples that had fewer than 1500 sequences, because we considered them as defective or unreliable outliers. Finally, we also removed 2 samples that did not have a valid body site label assigned to them. This resulted in a set of 9192 samples containing a total of 118 702 967 sequences with an average length of 413 bps. From these samples, sequences with a length of less than 150 bps as well as sequences longer than 540 bps were removed, as we considered them potentially erroneous. No further preprocessing (such as chimera detection) was applied. This resulted in a total of 116 520 289 sequences, of which 63 221 538 were unique. We then used the unconstrained *Bacteria* tree of our Phylogenetic Automatic (Reference) Trees (PhAT) method [34] for phylogenetic placement. The tree comprises 1914 taxa, thereof 1797 bacterial sequences. The remaining 117 taxa are *Archaea* and *Eukaryota*, and were included as a broad outgroup. Each sample is labeled with one of 18 human body site locations where it was sampled. This is the only publicly available meta-data feature.

For our re-analysis of the oral/fecal dataset of the original Phylofactorization [31], we used the test data provided at https://github.com/reptalex/phylofactor. We modified the scripts to produce 10 factors instead of the default of using their stopping criterion, in order to be comparable to our implementation and results. For the comparison with our Placement-Factorization, we selected a suitable oral/fecal subset of the HMP dataset as described in the main text.

### 9 S2 Text. Pipeline and implementation

#### 9.1 Phylogenetic Placement Pipeline

*Phylogenetic placement* (also called *evolutionary placement*) has been developed for conducting phylogenetic analyses of metagenomic sequence data [41]. It is implemented in tools such as pplacer [22], RAxML-EPA [23], and EPA-ng [24,111]. Instead of resolving the phylogeny of a set of metagenomic sequences, phylogenetic placement treats each sequence, called a *query sequence* (QS), separately. It evaluates how these QSs relate to an existing *reference tree* (RT) based on known reference sequences. For each QS, it computes the probabilities of *placing* the sequence on all branches of the RT, thereby classifying them into a phylogenetic context of related sequences, without the need to resolve relationships between the QSs.

In the most common use case, the QSs are reads or amplicons from environmental samples. Most often barcoding regions or marker genes such as 16S or 18S are used, but there also exist studies that use different, or even a set of, marker genes [9]. Furthermore, other types of sequences such as _mi_tags [50] can be used.

The RT and the reference sequences it represents are typically assembled by the user so that they capture the expected species diversity in the samples. To expedite this process, we recently proposed an automated approach for assembling suitable sets of reference sequences [34]. Distinct samples from one study are typically placed on the same underlying RT in order to facilitate comparisons between the samples.

We here assume to be given a set of suitable reference sequences, their alignment, and an RT inferred from them. In current implementations of phylogenetic placement, the RT has to be strictly bifurcating. Prior to the placement, the QSs need to be aligned against the reference alignment of the RT by programs such as P_A_P_A_R_A_ [35,36] or hmmalign [37,38]. The input to phylogenetic placement are (i) the reference tree (RT), (ii) its underlying alignment, and (iii) the aligned query sequences (QSs). The placement pipeline is shown in Fig 1.

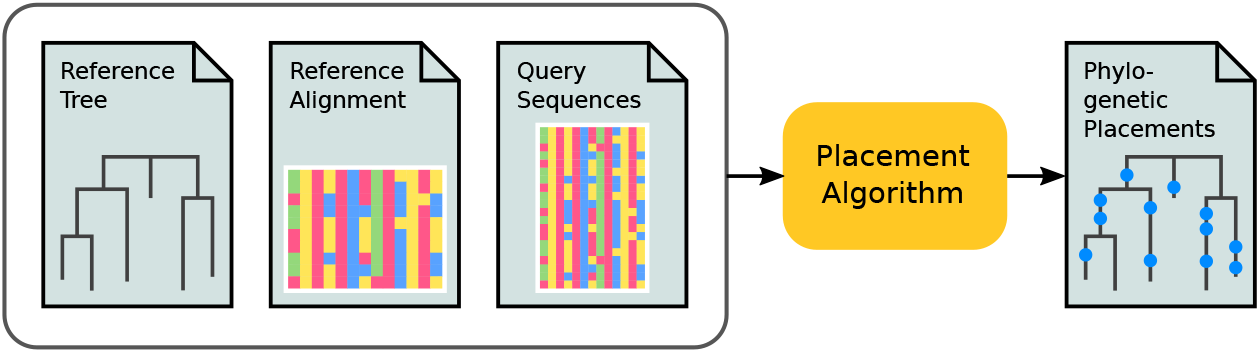

**Phylogenetic placement pipeline**. The input to phylogenetic placement are three files: the reference tree (RT), the corresponding reference alignment, and the aligned query sequences (QSs). The placement algorithm then computes the probabilities of placing the QSs on the branches of the RT, which are stored in an output file.

The output of phylogenetic placement are the probabilities of placing the QSs on the branches of the RT. The output data is usually stored in so-called `jplace` files [119]. It stores the RT in `Newick` format, including tip names and branch lengths. Its main part is the list of placements for each QS, which store the LWRs and placement positions along the branches of the RT.

#### 9.2 Method Implementation

The methods described in the manuscript are implemented in our tool gappa, which is freely available under GPLv3 at http://github.com/lczech/gappa. gappa internally uses our `C++11` library genesis, which offers functionality for working with phylogenies and phylogenetic placement data, and also contains methods to work with taxonomies, sequences and many other data types. genesis is also freely available under GPLv3 at http://github.com/lczech/genesis. This software design of using a library (genesis) for the core functions, and a separate program (gappa) for the user-facing command line interface, has the advantage of enabling experimentation and extension for future research.

gappa offers a command line interface for conducting typical tasks when working with phylogenetic placements. The methods that we described here are implemented via the following sub-commands:

- `dispersion`: The command takes a set of jplace files (the samples), and calculates and visualizes the Edge Dispersion per edge of the reference tree.
- `correlation`: The command takes a set of jplace samples, as well as a table containing metadata features for each sample. It then calculates and visualizes the Edge Correlation with the metadata features per edge of the reference tree.
- `phylogenetic-kmeans and imbalance-kmeans`: Performs *k*-means clustering of a set of jplace files according to our methods.
- `placement-factorization`: Performs our adaptation of Phylofactorization [31] to phylogenetic placement data, and outputs all relevant analysis results.
- `squash` and `edgepca`: Reimplementations of the two existing methods [29,56].

These are the gappa commands that are relevant for this paper. The tool also offers additional commands that are useful for phylogenetic placement data, such as visualization or filtering. At the time of writing this manuscript, gappa is under active development, with more functions planned in the near future. Furthermore, gappa has recently been bundled into a bioconda package, which is available at https://anaconda.org/bioconda/gappa; note that this package is not maintained by ourselves. Lastly, we provide prototype implementations, scripts, data, and other tools used for the tests and figures in this paper at http://github.com/lczech/placement-methods-paper.

**S1 Table.** HMP Dataset Overview. The table lists the 19 body site labels used by the Human Microbiome Project (HMP) [16,17]. We used this dataset to evaluate the applicability of our methods for phylogenetic placement. In order to simplify the visualization in several figures, we summarized some of the labels into eight location regions, as shown in the second column. The last column lists how many samples from each body site were used in our evaluation.

| Body Site | Region | Samples |
| --- | --- | --- |
| Tongue Dorsum | Mouth (back) | 610 |
| Palatine Tonsils | Mouth (back) | 599 |
| Throat | Mouth (back) | 638 |
| Attached Keratinized Gingiva | Mouth (front) | 600 |
| Hard Palate | Mouth (front) | 566 |
| Buccal Mucosa | Mouth (front) | 597 |
| Saliva | Saliva | 529 |
| Supragingival Plaque | Plaque | 608 |
| Subgingival Plaque | Plaque | 595 |
| Anterior Nares | Airways | 541 |
| Left Retroauricular Crease | Skin | 596 |
| Right Retroauricular Crease | Skin | 604 |
| Left Antecubital Fossa | Skin | 290 |
| Right Antecubital Fossa | Skin | 328 |
| Stool | Stool | 600 |
| Vaginal Introitus | Vagina | 292 |
| Mid Vagina | Vagina | 298 |
| Posterior Fornix | Vagina | 301 |
| Sum |  | 9192 |

**S2 Table.** Effect of Branch Binning on the KR Distance of the HMP Dataset. Here we show the effect of per-branch placement binning on the run-time and on the resulting relative error when calculating the pairwise KR distance matrix between samples, by example of the Human Microbiome Project (HMP) [16,17] dataset. Because of the size of the dataset (9192 samples) and reference tree (1914 taxa), we executed this evaluation in parallel on 16 cores. The first row shows the baseline performance, that is, without binning. When using fewer bins per branch, the run-time decreases, at the cost of slightly increasing the average relative error. Still, even when compressing the placement masses into only one bin per branch (that is, just using per-branch masses), the average relative error of the KR distances is around 1%, which is acceptable for most applications. However, considering that the run-time savings are not substantially better for a low number of bins, we recommend using a relatively large number of bins, e.g., 32 or more. This is because run-times of KR distance calculations also depend on other effects such as the necessary repeated tree traversals. We also conducted these tests on the BV dataset, were the relative error is even smaller.

| Bins | Time (h:mm) | Speedup | Relative $\Delta$ |
| --- | --- | --- | --- |
| - | 9:46 | 1.00 | 0.000000 |
| 256 | 6:58 | 1.40 | 0.000008 |
| 128 | 6:39 | 1.47 | 0.000015 |
| 64 | 6:30 | 1.50 | 0.000035 |
| 32 | 6:25 | 1.52 | 0.000124 |
| 16 | 6:13 | 1.57 | 0.000272 |
| 8 | 6:08 | 1.59 | 0.000669 |
| 4 | 6:07 | 1.60 | 0.002747 |
| 2 | 6:04 | 1.61 | 0.004284 |
| 1 | 5:35 | 1.75 | 0.011585 |

**S3 Table.** First ten factors of the BV dataset found by Phylofactorization. We analyzed the BV dataset with the original Phylofactorization, using two different methods for the OTU clustering of the data, namely vsearch [54] and swarm [52,53]; see S1 Text for details on the preprocessing. Here, we compare our Placement-Factorization of the dataset to these results. As the original implementation does not support taxon weighting, we also do not use it here. The table shows the clades split by the first 10 factors found by each variant. See also S14 Fig for a visualization of the clades found by our adaptation. As shown in S1 Fig and in the original study of the dataset [18], there are multiple different taxa that are associated with Bacterial Vaginosis. That is, there are several clades or branches of our reference tree where the placement mass differs between healthy and sick patients. It is thus expected that a phylo-factorization of these data exhibits some variation in the exact clade found, depending on the preprocessing and exact settings being used. Still, the table shows that—apart from ordering—the factored clades are mostly consistent across variants, and consistent with previous findings. All of the taxa found by the swarm-based Phylofactorization and by our Placement-Factorization, as well as all taxa except some of the *Streptococcus* found as part of the first factor of the vsearch-based Phylofactorization, were already shown to play important roles for this dataset [18]. The inclusion of *Streptococcus* in the vsearch variant is due to an inner edge that has a slightly higher value of the objective function than the actually more relevant edges leading to the *Lactobacillus* clade. We observed a similar behavior of large clades being split with our implementation when using taxon weights, as shown in Fig 10. Lastly, the normalized mutual information [120] between the three variants ranges between 71% and 81%, further showing that they mostly find the same clades.

|  | Original (vsearch) | Original (swarm) | Placement-Factorization |
| --- | --- | --- | --- |
| 1 | *Lactobacillus crispatus*,<br>*Lactobacillus jensenii*,<br>*Lactobacillus iners*,<br>*Lactobacillus coleohominis*,<br>*Lactobacillus gasseri*,<br>*Lactobacillus vaginalis*,<br>*Streptococcus agalactiae*,<br>*Streptococcus anginosus*,<br>*Streptococcus gallolyticus*,<br>*Streptococcus oralis*,<br>*Aerococcus christensenii* | *Sneathia sanguinegens*,<br>*Leptotrichia amnionii* | *Lactobacillus crispatus*,<br>*Lactobacillus jensenii*,<br>*Lactobacillus kalixensis* |
| 2 | *Lactobacillus crispatus* | *Lactobacillus crispatus* | *Sneathia sanguinegens*,<br>*Leptotrichia amnionii* |
| 3 | *Gardnerella vaginalis* | *Gardnerella vaginalis* | *Gardnerella vaginalis* |
| 4 | *Leptotrichia amnionii* | *Atopobium vaginae* | *Megasphaera* |
| 5 | *Megasphaera* | *Megasphaera* | *Lactobacillus crispatus* |
| 6 | *Atopobium vaginae* | *Eggerthella* | *Eggerthella* |
| 7 | *Eggerthella* | *Prevotella bivia*,<br>*Prevotella amnii* | *Prevotella timonensis*,<br>*Prevotella buccalis* |
| 8 | *Sneathia sanguinegens* | *Prevotella timonensis* | *Prevotella bivia*,<br>*Prevotella amnii* |
| 9 | *Prevotella timonensis* | BVAB2 | *Atopobium vaginae* |
| 10 | *Lactobacillus jensenii* | *Lactobacillus jensenii* | *Lactobacillus iners* |

**S1 Fig.**
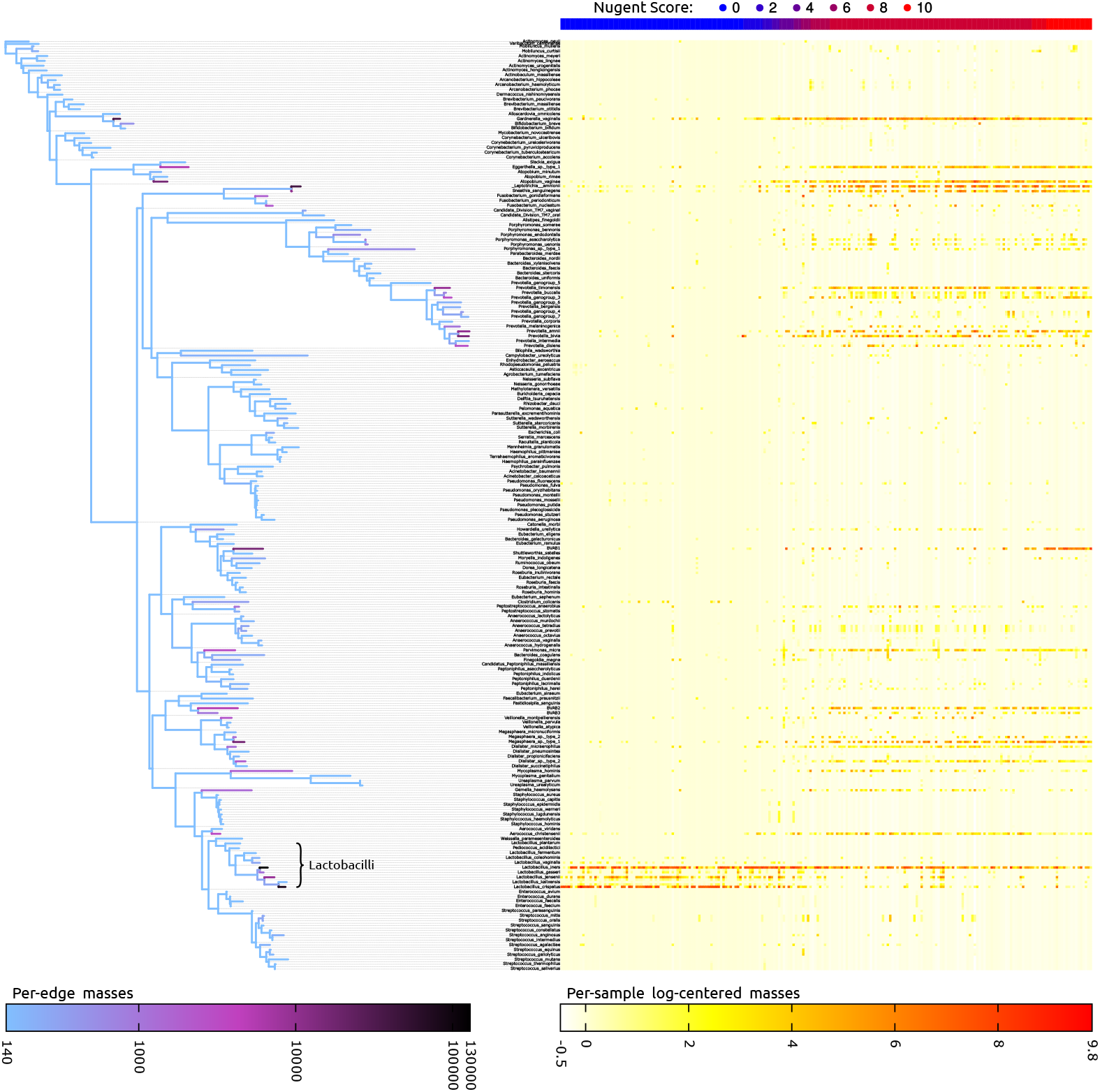
Visualization of per-edge and per-sample masses of the BV dataset. The figure provides an overview of the placement of the BV dataset [18]: The left hand side shows a condensed version of the original reference tree of [18], colored by log-scaled placement mass of all samples accumulated. For clarity and simplicity, in this figure we used a reference tree built from the consensus sequences of each original reference taxon, so that each species is represented by exactly one tip here. The *Lactobacillus* clade is highlighted in the tree, which is an important clade for this dataset. Note the two particularly dark branches, *Lactobacillus iners* and *Lactobacillus crispatus*, which are the major species associated with a healthy vaginal microbiome [18]. The right hand side shows a heat map that further resolves the placement masses per sample: Each row corresponds to a branch of the tree on the left (note that dashed lines also start from inner branches), and each column represents one sample. The values are log-centered in order to be consistent with typical OTU abundance heat map representations, for example as in [31]. The samples/columns are sorted by their Nugent score, from 0 at the left for the healthy patients to 10 at the right for the sick ones. The Nugent score of each sample is also shown at the top as a blue to red bar. Here again the high abundance of *Lactobacillus* in healthy patients is visible in the lower part of the matrix, while the diseased patients exhibit high placement masses at several other taxa [18].

**S2 Fig.**
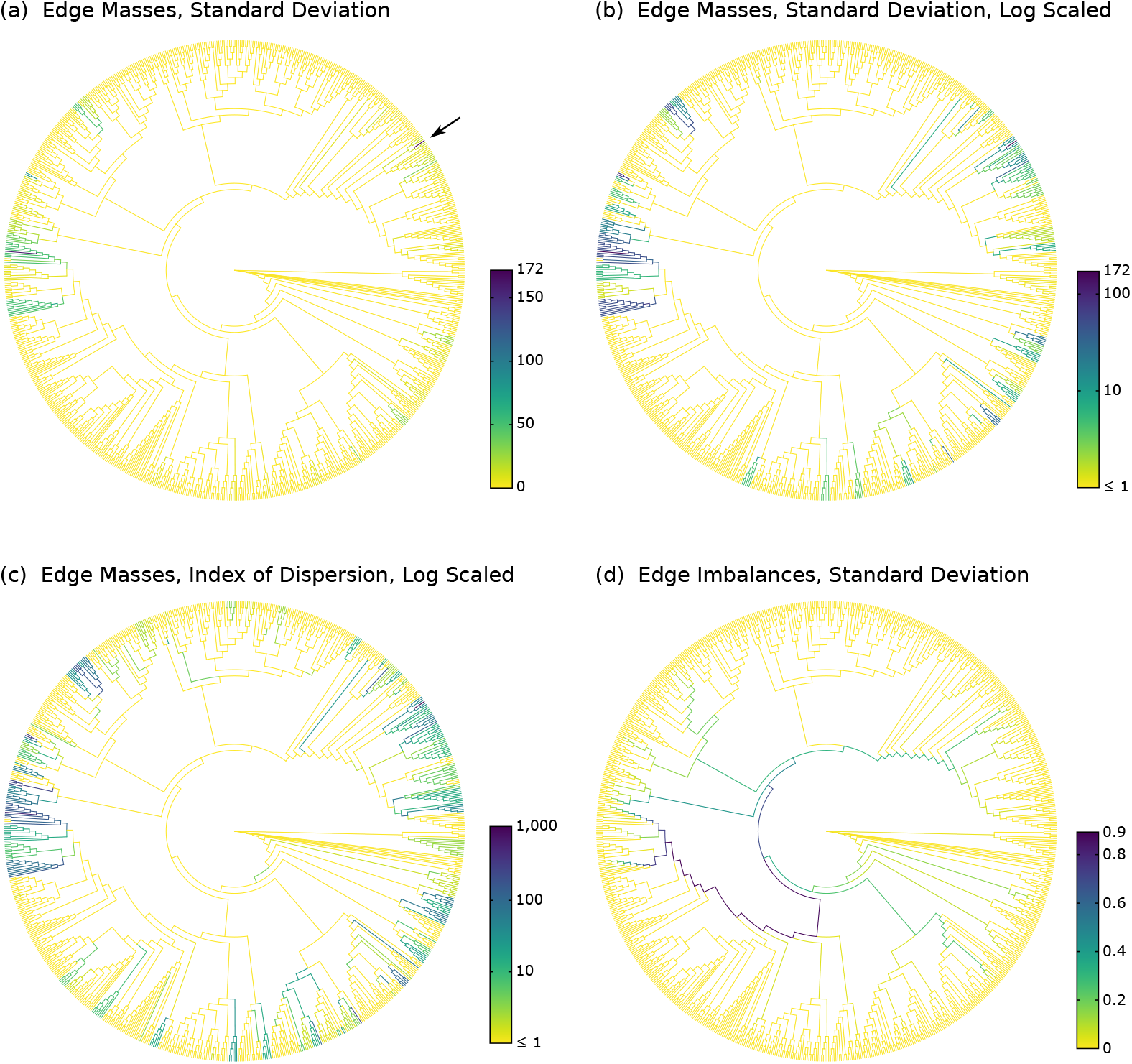
Examples of variants of Edge Dispersion. We re-analyzed the BV dataset to show variants of our Edge Dispersion method. All subfigures highlight the same branches and clades as found by other methods such as Edge PCA. The method is useful as a first exploratory tool to detect placement heterogeneity across samples. In contrast to Edge Correlation, it can however not explain the reasons of heterogeneity. Subfigure (a) shows the standard deviation of the absolute edge masses, without any further processing. It is striking that one outlier, marked with an arrow, is dominating, thus hiding the values on less variable edges. This outlier occurs at the species *Prevotella bivia* in one of the 220 samples, where 2781 out of 2782 sequences in the sample have placement mass on that branch. Upon close examination, this outlier can also be seen in Figure 1D of [18], but is less apparent there. Subfigure (b) is identical to Fig 4(a) of the main text and shows the standard deviation again, but this time using logarithmic scaling, thus revealing more details on the edges with lower placement mass variance. Furthermore, when comparing it to S3(c) Fig, we see that the same clades that exhibit a high correlation or anti-correlation with meta-data there are also highlighted here. Subfigure (c) shows the Index of Dispersion of the edge masses, that is, the variance normalized by the mean. Hence, edges with a higher number of placements are also allowed to have a higher variance. We again use a logarithmic scale because of the outlier. The figure reveals more details on the edges with lower variance, highlighted in medium green colors. Subfigure (d) shows the standard deviation of edge imbalances. Because we used imbalances of unit mass samples, the values are already normalized. The path to the *Lactobacillus* clade is again clearly visible, indicating that the placement mass in this clade has a high variance across samples. Note that imbalances can be negative; thus, the Index of Dispersion is not applicable to them.

**S3 Fig.**
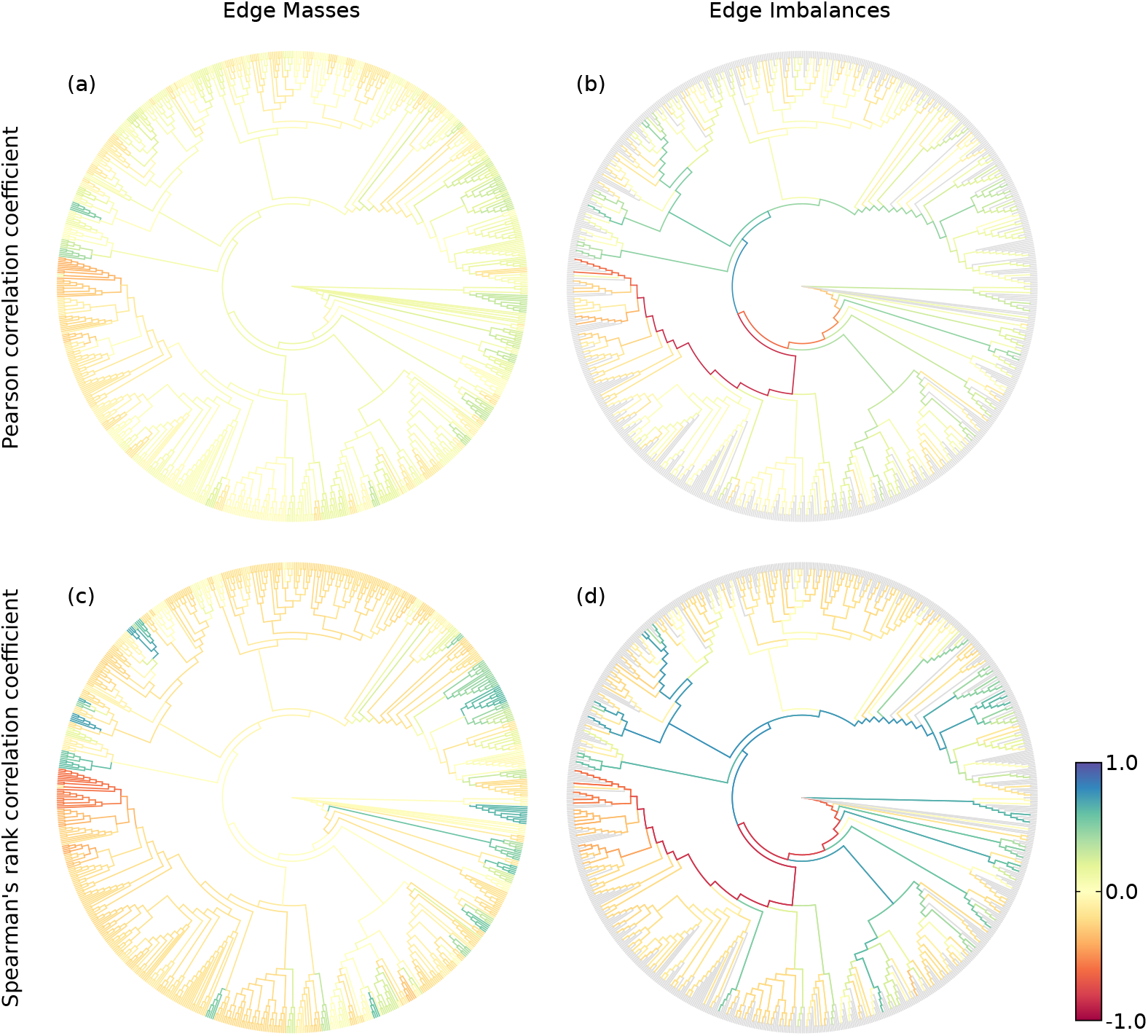
Examples of variants of Edge Correlation. We again use the BV dataset, and show the correlation of edge masses and imbalances with the Nugent score. The Nugent score measures the severeness of Bacterial Vaginosis, and ranges from 0 for healthy subjects to 10 for heavily affected patients. Subfigures (a) and (b) use the Pearson Correlation Coefficient, that is, they show the linear correlation with the meta-data feature, while subfigures (c) and (d) use Spearman’s Rank Correlation Coefficient and thus show monotonic correlations. Subfigure (d) is identical to Fig 4(b) of the main text. All subfigures show red edges or red paths at the *Lactobacillus* clade. This indicates that presence of placements in this clade is anti-correlated with the Nugent score, which is consistent with the findings of [18] and [29]. In other words, presence of *Lactobacillus* correlates with a healthy vaginal microbiome. On the other hand, blue and green edges, which indicate positive correlation, are indicative of edges that correlate to Bacterial Vaginosis. The extent of correlation is larger for Spearman’s Coefficient, indicating that the correlation is monotonic, but not strictly linear.

**S4 Fig.**
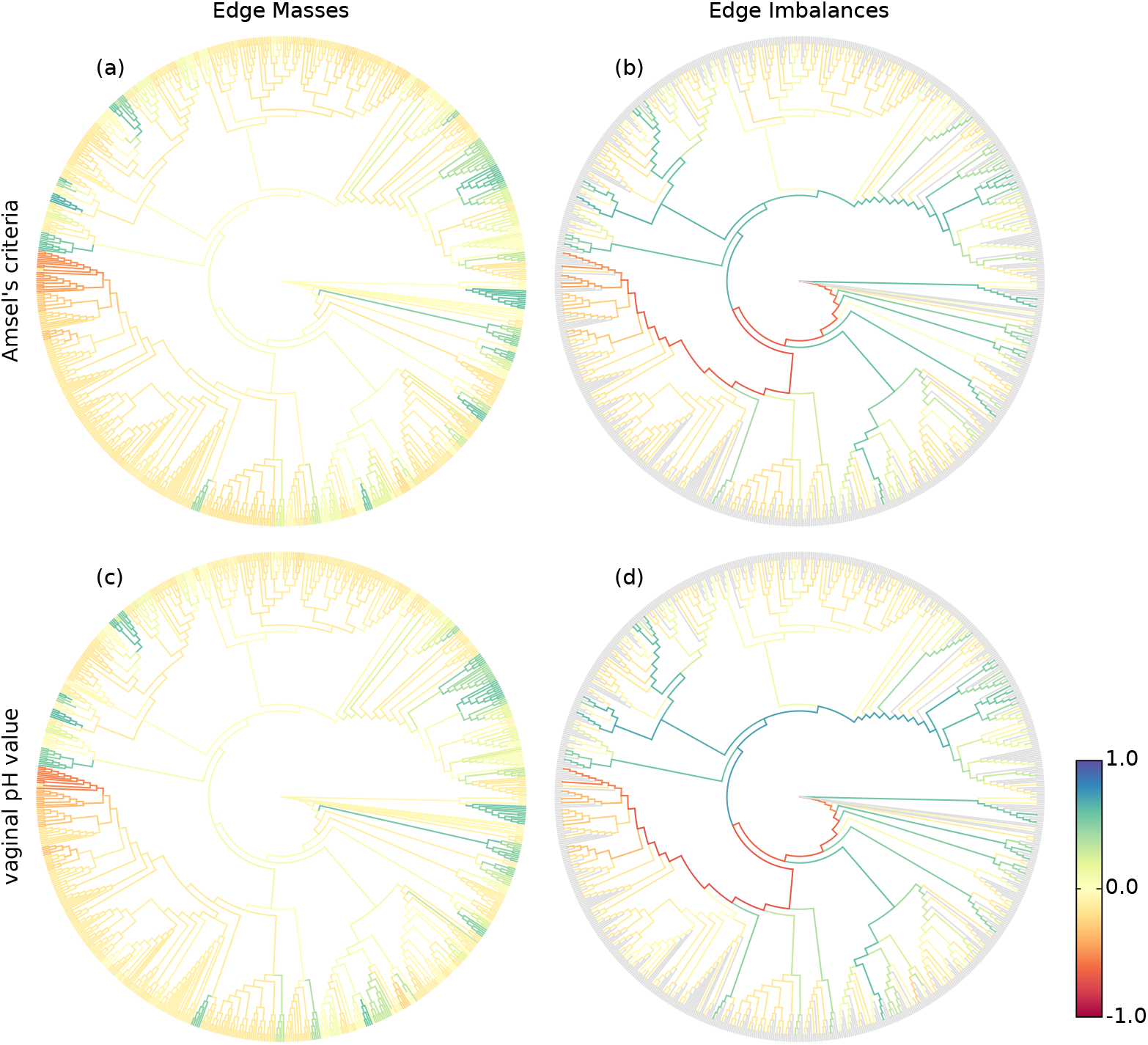
Edge Correlation with more meta-data features. Here, we use additional meta-data features of the BV dataset to show that Edge Correlation yields consistent results with existing methods. In particular, we caltucated Spearman’s Coefficient with Amsel’s criteria [98] in subfigures (a) and (b), as well as with the vaginal pH value in subfigures (c) and (d). Both features were also used in [18] as indicators of Bacterial Vaginosis. The figures are almost identical to the ones shown in S3 Fig; that is, they yield results that are consistent with the previously used Nugent score, as well as consistent with existing methods.

**S5 Fig.**
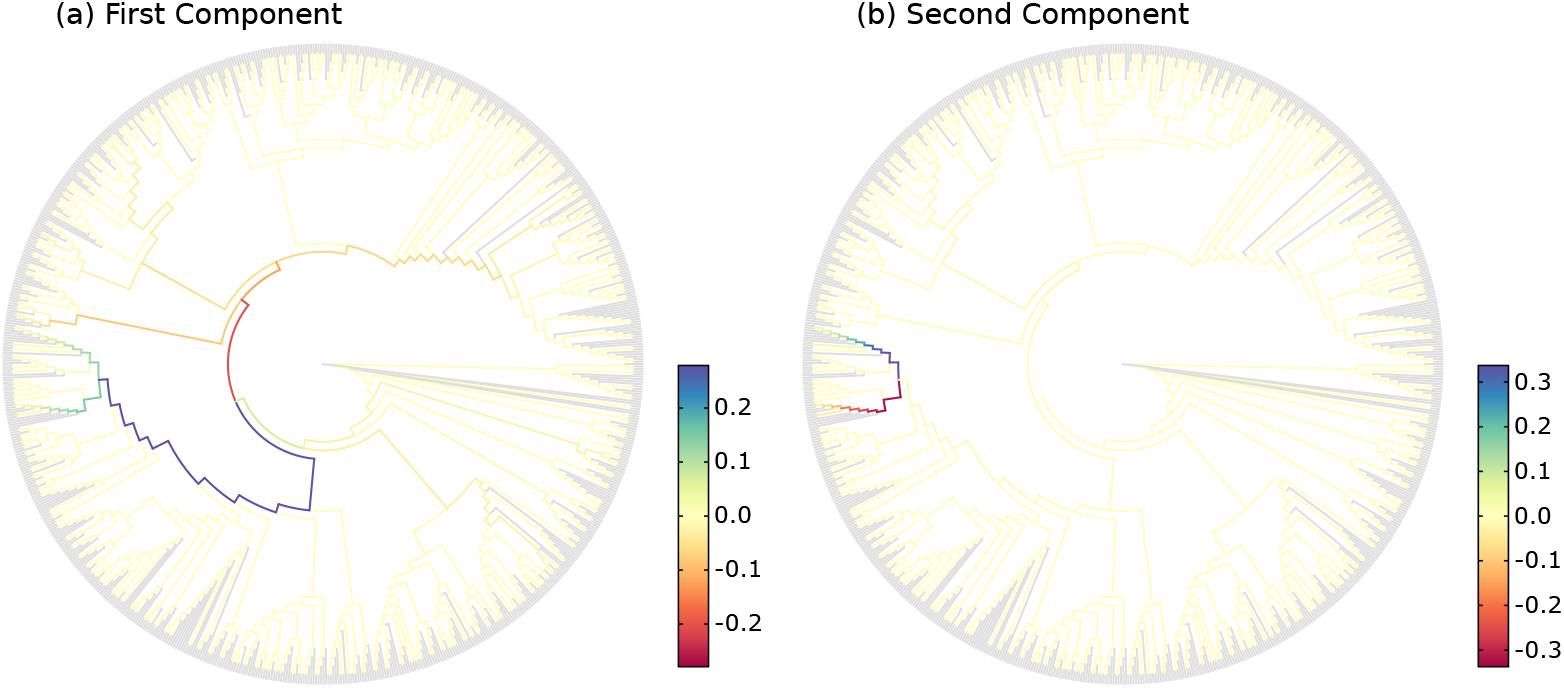
Recalculation of the Edge PCA tree visualization. Subfigures (a) and (b) are recalculations of Figures 4 and 5 of [29], respectively. However, we show them here in our coloring scheme in order to facilitate comparison with other figures. The original publication instead uses two colors for a positive and a negative sign of the principal components, and branch width to show their magnitude. Note that the actual sign is arbitrary, as it is derived from principal components. The figure shows the first two Edge PCA components, visualized on the reference tree. This form of visualization is useful to interpret results such as the Edge PCA projection plot as shown in Fig 8(e) of the main text. It reveals which edges are mainly responsible for separating the samples into the PCA dimensions. Here, the first principal component in (a) indicates that the main PCA axis separates samples based on the presence of placements in the *Lactobacillus* clade, which is what the blue and green path leads to. The second component in (b) then further distinguishes between two species in this clade, namely *Lactobacillus iners* and *Lactobacillus crispatus*.

**S6 Fig.**
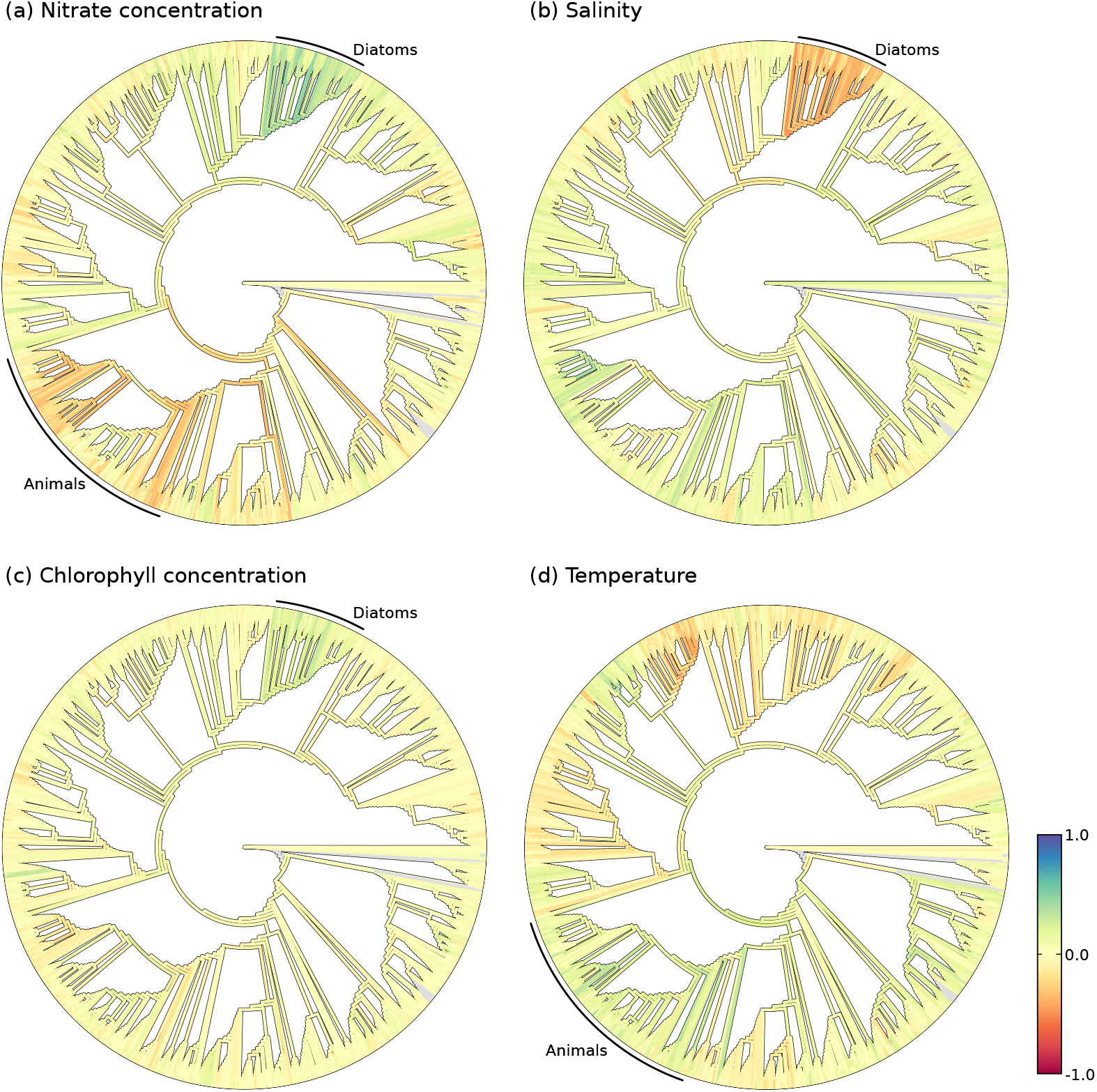
Examples of Edge Correlation using Tara Oceans samples. The figure shows the correlation of Tara Oceans sequence placements with (a) the nitrate, (b) the salinity, (c) the chlorophyll, and (d) the temperature sensor data of each sample. The sensor values range from −2.2 to 33.1 μmol/l (nitrate), from 33.2 to 40.2psu (salt), from −0.02 to 1.55mg/m^3^ (chlorophyll), and from −0.8 to 30.5°C (temperature), respectively. The negative nitrate and chlorophyll concentrations are values below the detection limit of the measurement method (pers. comm. with L. Guidi), and hence simply denote low concentrations. We used Spearman’s Rank Correlation Coefficient, and examine two exemplary clades, namely the *Animals* and the *Diatoms*. Diatoms are mainly photosynthetic, and thus depend on nitrates as key nutrients, which is clearly visible by the high correlation of the clade in (a). Furthermore, the diatoms exhibit positive correlation with the chlorophyll concentration (c), which again is indicative of their photosynthetic behavior. On the other hand, they show a high anti-correlation with the salt content (b). Salinity is a strong environmental factor which heavily affects community structures and species abundances [99], particularly diatoms [100]. The correlations of the animal clade are less pronounced. They exhibit a negative correlation with nitrate (a), as well as an increase in absolute abundance with higher temperatures (d). While these findings are not surprising, they show that the method is able to find meaningful relationships in the data.

**S7 Fig.**
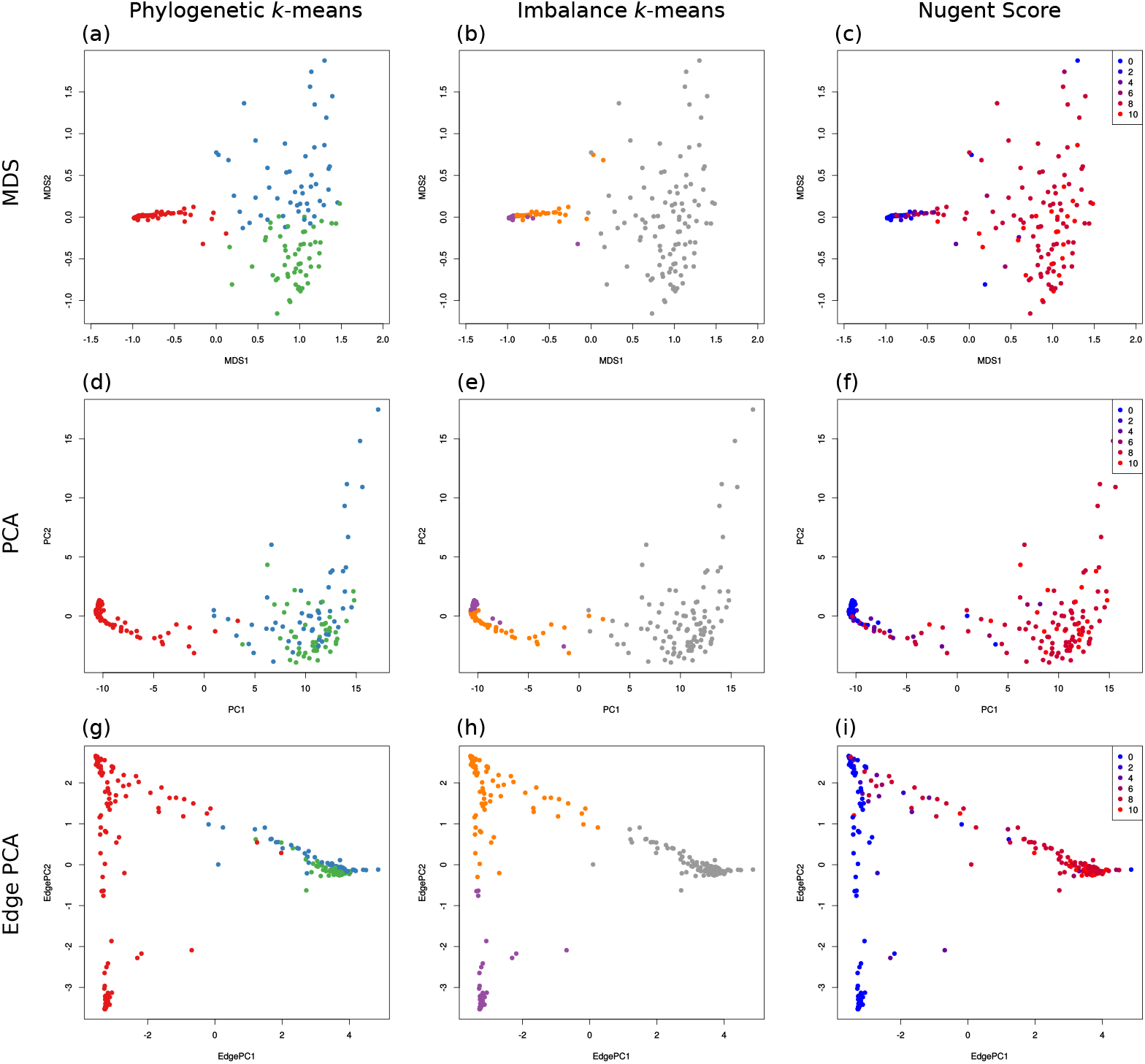
Comparison of *k*-means clustering to MDS, PCA, and Edge PCA. Here, we show and compare the dimensionality reduction methods MDS, PCA, and Edge PCA (one per row). MDS and PCA were calculated on the pairwise KR distance matrix of the BV dataset, Edge PCA was calculated using the placements on the re-inferred RT of the original publication [18]. The plots are colored by the cluster assignments as found by our *k*-means variants (first two columns), and by the Nugent score of the samples (last column). The Nugent score is included to allow comparison of the health status of patients with the clustering results. (a), (d) and (h) are identical to Figs 8(c), (d) and (e) of the main text, respectively. (f) and (i) are recalculations of Figures 4 and 3 of [101], respectively. This figure reveals additional details about how the *k*-means method works, that is, which samples are assigned to the same cluster. For example, the purple cluster found by Imbalance *k*-means forms a dense cluster of close-by samples on the left in (b) and (e), which is in accordance with the short branch lengths of this cluster as shown in Fig 8(b) of the main text.

**S8 Fig.**
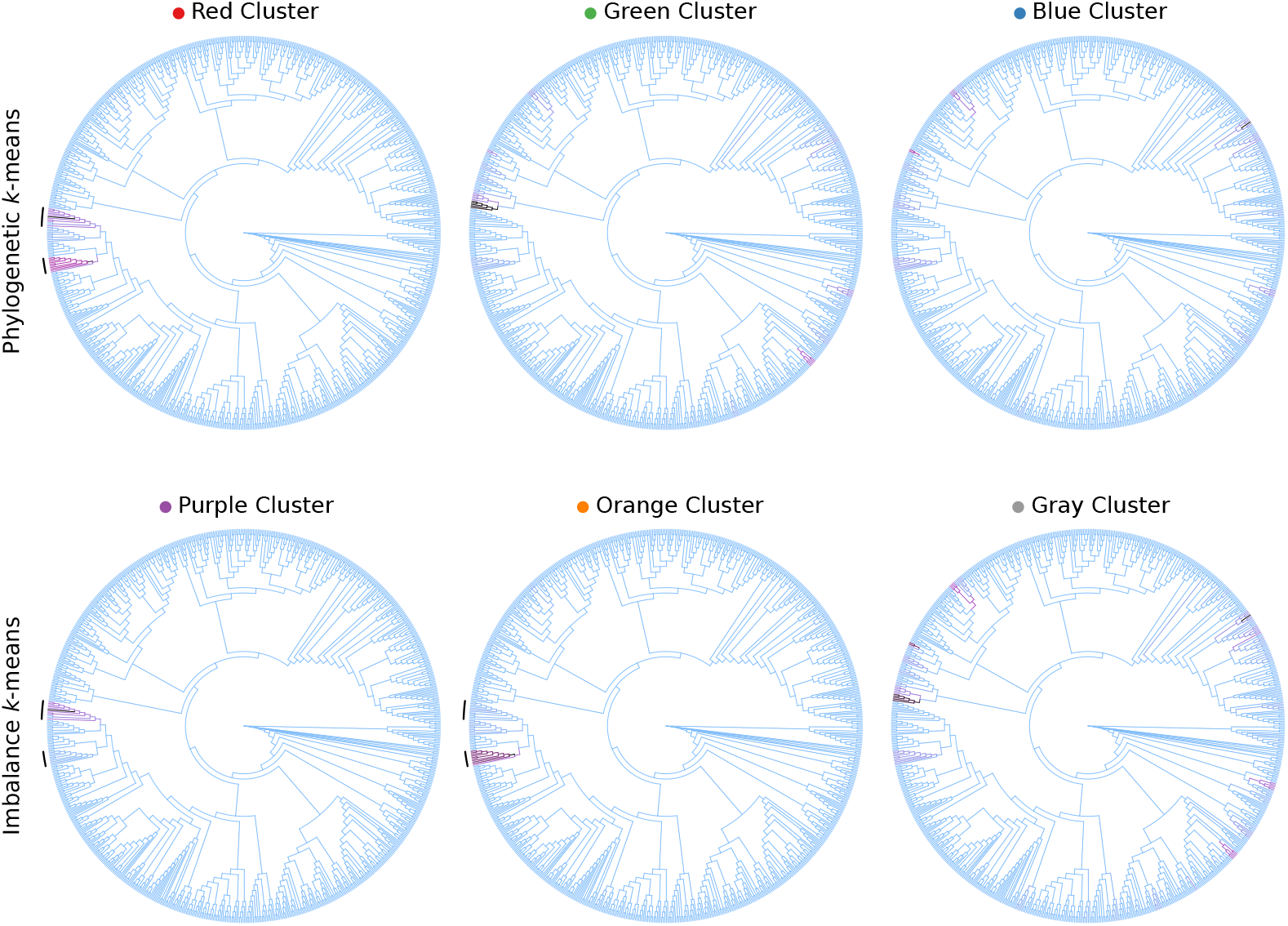
Example of *k*-means cluster centroids visualization. Here we show the cluster centroids as found by our *k*-means variants using the BV dataset, visualized on the reference tree via color coding. The cluster assignments are the same as in Fig 8 of the main text; the first row show the three clusters found by Phylogenetic *k*-means, the second row the clusters found by Imbalance *k*-means. Each tree represents one centroid around which the samples were clustered, that is, it shows the combined masses of the samples that were assigned to that cluster. The edges are colored relative to each other, using a linear scaling of light blue (no mass), purple (half of the maximal mass) and black (maximal mass). As explained in the main text, the samples can be split into three groups: The diseased subjects, which have placement mass in various parts of the tree, as well as two groups of healthy subjects, with placement mass in one of two *Lactobacillus* clades (marked with black arcs on the left of the trees). This grouping is also clearly visible in these trees. The red cluster for example represents all healthy subjects, and thus most of its mass is located in the two *Lactobacillus* clades. The purple and orange clusters on the other hand show a difference in placement mass between those clades. Furthermore, the placement mass of the gray cluster is mostly a combination of the masses of the green and blue cluster, all of which represent diseased subjects. These observations are in accordance with previous findings as explained in the main text.

**S9 Fig.**
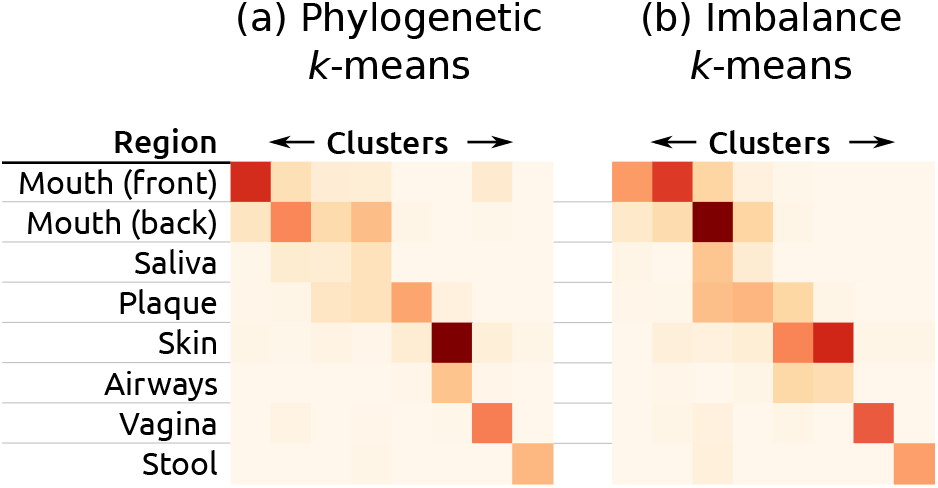
Clustering using Phylogenetic *k*-means on the HMP dataset. *k* is set to 8, instead of *k* := 18 as in the main text, based on a coarse aggregation of the original body site labels as shown in S1 Table. See Fig 8 for the cluster assignment where *k* is set to the original number of labels; there, we also list how the labels were aggregated. Each row represents a body site; each column one of the 8 clusters. The color values indicate how many samples of a body site were assigned to each cluster. Some of the body sites can be clearly separated, while particularly the samples from the oral region are distributed over different clusters. This might be due to homogeneity of the oral samples.

**S10 Fig.**
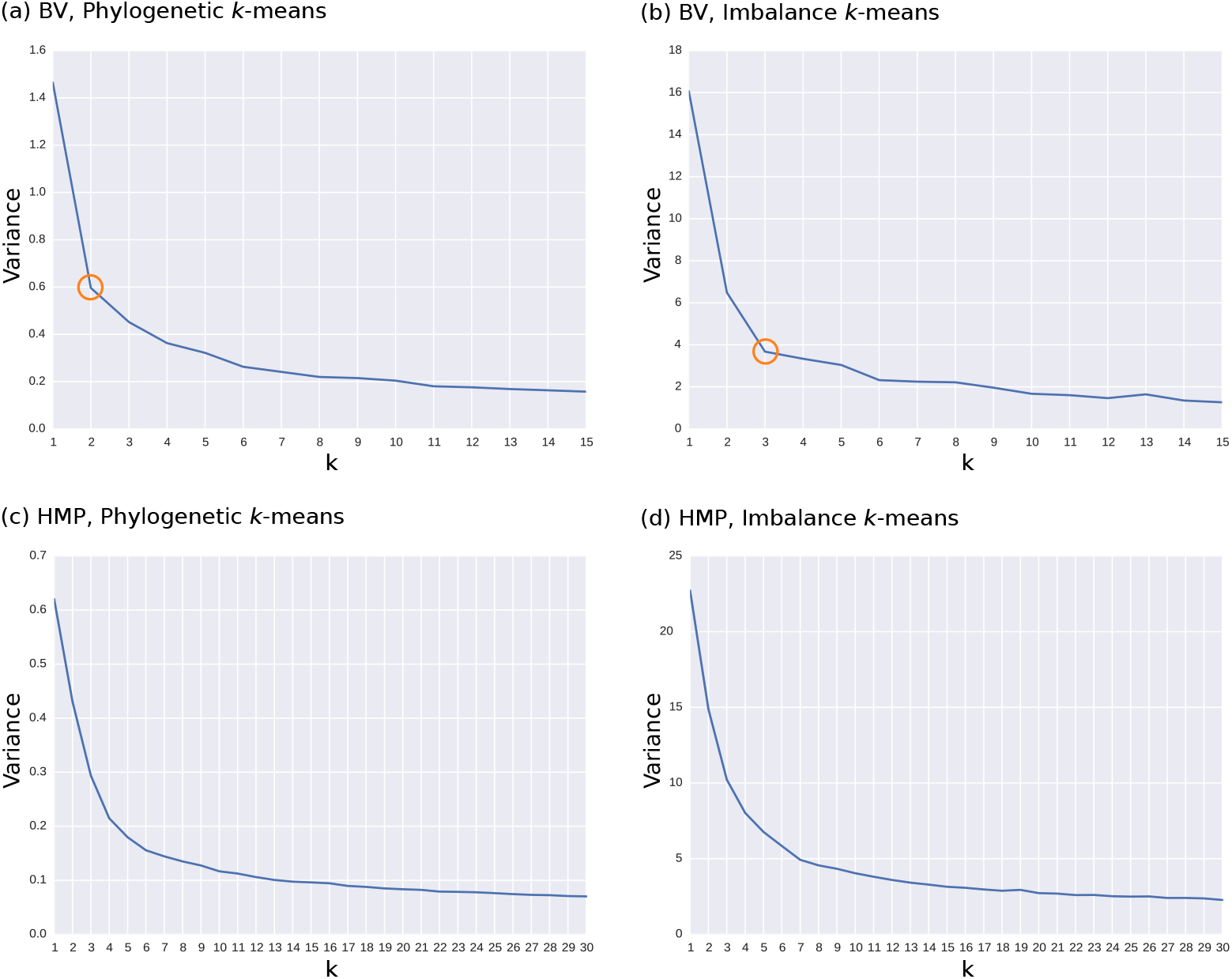
Variances of *k*-means clusters in our test datasets. The figures show the cluster variance, that is, the average squared distance of the samples to their assigned cluster centroids, for different values of *k*. The first row are clusterings of the BV dataset, the second row of the HMP dataset. They were clustered using Phylogenetic *k*-means (first column), and Imbalance *k*-means (second column), respectively. Accordingly, (a) and (c) use the KR distance, while (b) and (d) use the euclidean distance to measure the variance. These plots can be used for the Elbow method in order to find the appropriate number of clusters in a dataset [79]. Low values of *k* induce a high variance, because many samples exhibit a large distance from their assigned centroid. On the other hand, at a given point, higher values of *k* only yield a marginal gain by further splitting clusters. Thus, if the data has a natural number of clusters, the corresponding *k* produces an angle in the plot, called the “elbow”. For example, (a) and (b) exhibit the elbow at *k* := 2 and 3, respectively, which are marked with orange circles. These values are consistent with previous findings, for instance, Fig 8: There, Phylogenetic *k*-means splits the samples into a distinct red cluster and the nearby green and blue clusters, while Imbalance *k*-means yields three separate clusters in purple, orange, and gray. For the HMP dataset, the elbow is less pronounced. We suspect that this is due to the broad reference tree not being able to adequately resolve fine-grained differences between samples, see S1 Text for details. Likely candidates for *k* are 4 – 6 for (c) and around 7 for (d). These values are consistent with the number of coherent “blocks” of clusters, which can be observed in Fig 9. Clearer results for this dataset might be obtained with other methods for finding “good” values for *k*, although we did not test them here.

**S11 Fig.**
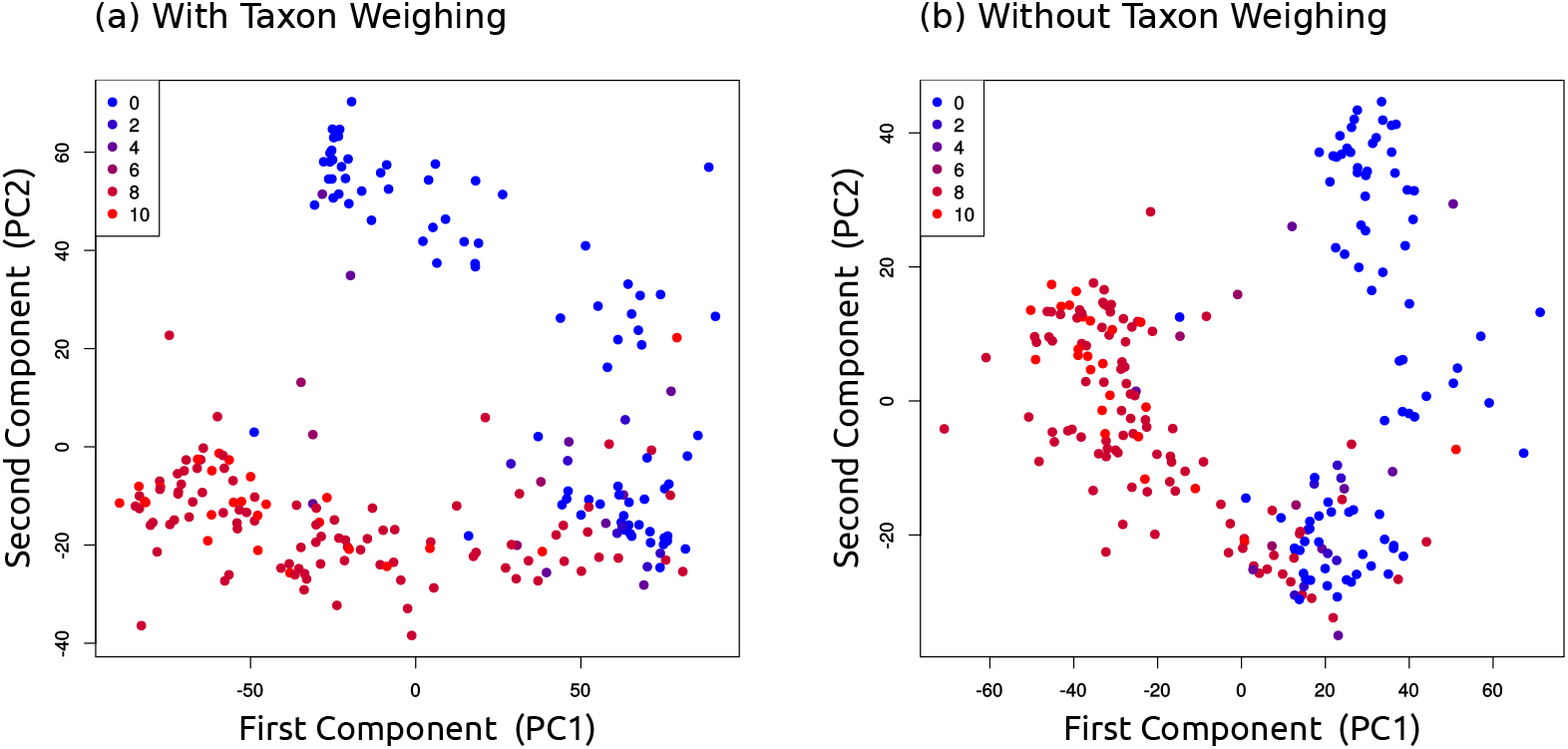
Projection of PCA on the edge balances of the BV dataset. The plots show the first two principal components of a PCA on the per-edge balances, calculated on placements of the data on reference tree of the BV dataset. That is, for each edge of the tree, we calculated the balance (log-ratio of geometric means) of the placement masses of the BV samples between the two sides of the tree induced by the edge. (a) shows the result when using taxon weighting [30] in the balances calculation, while (b) shows the result without taxon weighting. Each item represents a sample, colored by its Nugent score (0 means healthy, 10 means severe illness); the Nugent score had no influence on the PCA calculations. Both plots separate the healthy from the sick patients. In contrast to Edge PCA, the first component of (a) does not fully distinguish between the healthy (blue) and diseased (red) samples. For some reason, the component only takes *Lactobacillus iners* into account, while mostly ignoring *Lactobacillus crispatus*. This can be seen in S12(a) Fig, which shows the eigenvectors of this component visualized on the reference tree. There, the path leading to the *Lactobacillus* clade does not include the branches of *Lactobacillus crispatus*, which is marked with a black arc. Including the second component however, which distinguishes between the two types of *Lactobacillus*, as shown in S12(b) Fig, yields a clear separation of the samples. Subfigure (b) exhibits closer similarities to the Edge PCA plot shown in S7(i) Fig, in that the first component separates healthy from sick, and the second component further splits the healthy individuals apart. The edges that are responsible for these splits can be seen in more detail in S12 Fig, where we visualize the first two principal components of the balances without taxon weighting on the tree.

**S12 Fig.**
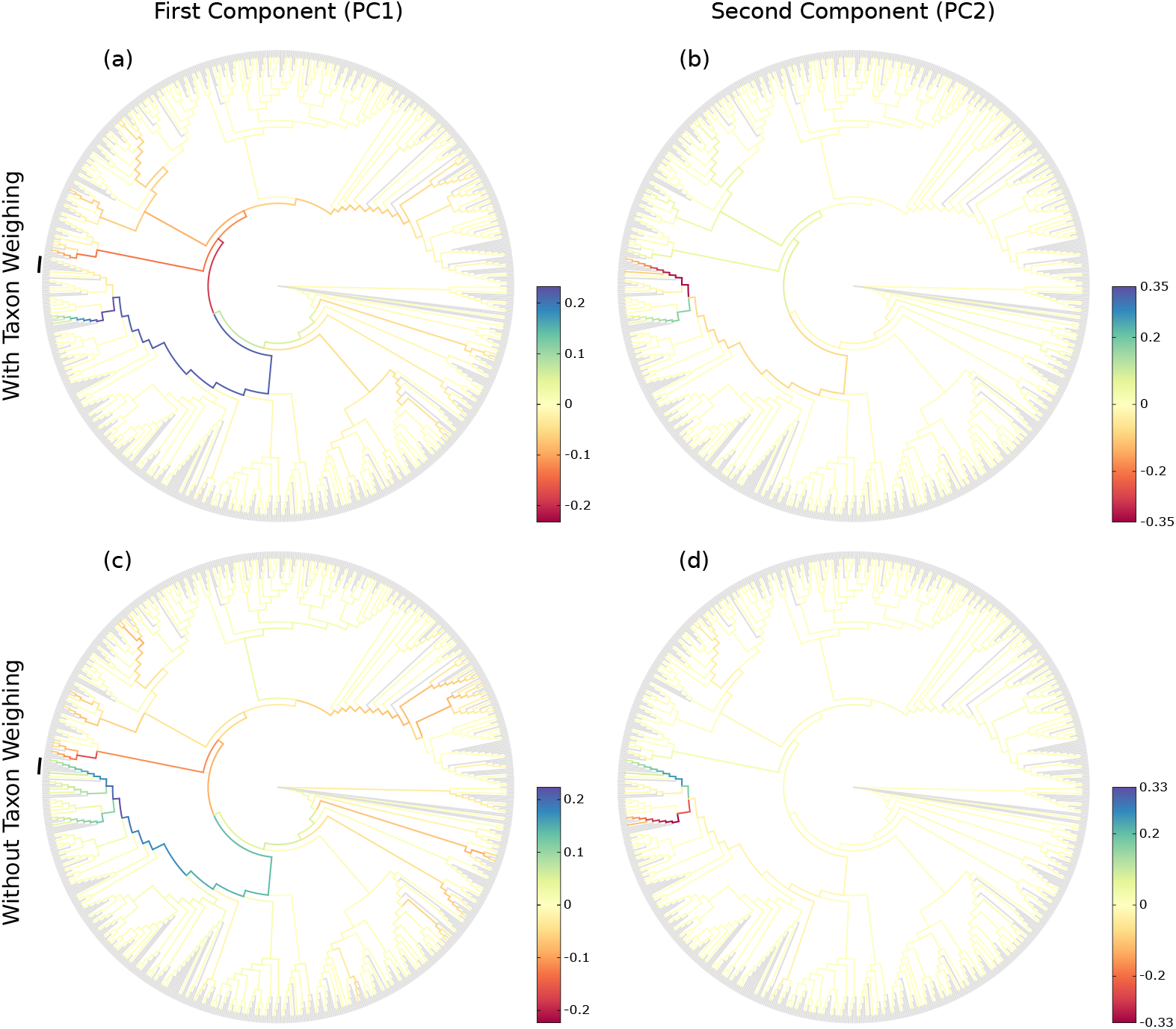
Eigenvectors of PCA on the edge balances of the BV dataset. The figure shows the eigenvectors of the first two principal components of PCA on the per-edge balances with and without taxon weighting, visualized on the reference tree of the BV dataset. See S11 Fig for details on the balances calculation. The visualization of the components on the reference tree is analogous to the Edge PCA tree visualization as for example shown in S5 Fig. As the data that is considered in the PCA corresponds to the edges of the tree, the resulting eigenvectors can be mapped back onto the tree, which is shown here. Each edge is colored according to the corresponding value of the first principal component in (a) and (c), and the second principal component in (b) and (d), respectively. The figure hence indicates how the axes in S11 Fig can be interpreted: The first component leads to the *Lactobacillus* clade, while the second one splits this clade into *Lactobacillus iners* and *Lactobacillus crispatus*. Hence, the results obtained from this analysis are consistent with our previous findings, c. f., S5 Fig and [18]. In (a) and (c), we marked the *Lactobacillus crispatus* clade with a black arc at the left of the tree.

**S13 Fig.**
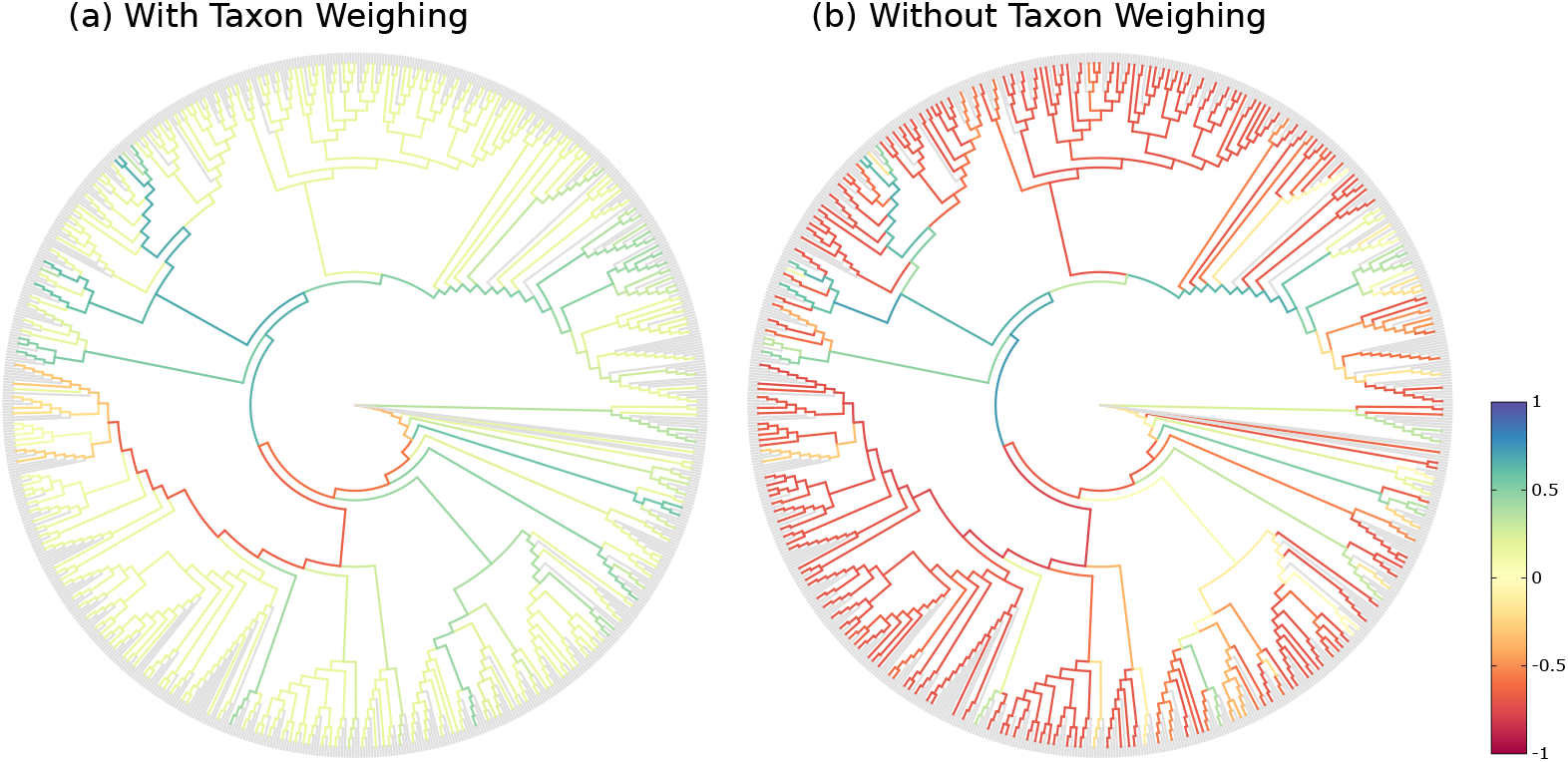
Correlation of the edge balances of the BV dataset with Nugent score. The Edge Correlation method as presented in the main text, and for example shown in S3 Fig, can also be conducted using balances (instead of masses or imbalances). We here show Edge Correlation using Spearman’s Rank Correlation Coefficient, calculated on the per-edge balances and the Nugent score, based on the placement of the BV dataset. That is, for each edge of the tree, we calculated the balance (log-ratio of geometric means) of the placement masses of the BV samples between the two sides of the tree induced by the edge. Then, we calculated the correlation with the Nugent score of each sample, and visualized it on the tree. (a) shows the result when using taxon weighting [30] in the balances calculation, while (b) shows the result without taxon weighting. With taxon weighting in (a), the result is similar to the correlation with imbalances shown in S3(d) Fig: An anti-correlation with the *Lactobacillus* clade is again visible (less placement mass in this clade means higher Nugent score, that is, indicates a more severe illness), while several other clades exhibit a positive correlation with Nugent score. The most striking difference to the previous Edge Correlation trees in S3 Fig is the majority of spuriously anti-correlated (red) edges in the tree without taxon weighting in (b). As mentioned in the main text, this is due to the insensitivity of the geometric mean to the presence of singular large values: If most values are small, so will be their geometric mean, even if a few very large values are also present. As can be seen in S1 Fig and S2 Fig, the clades that exhibit a high anti-correlation here (red) have low placement mass with a low variance. Hence, the geometric mean of the masses in these clades is consistently low across samples, which means that the numerator of the log-ratio in the balances computation has little effect on the correlation. This implies that the denominator, which represents the rest of the tree, drives the anti-correlations seen here. Women affected by BV show a presence of several different bacterial clades, while healthy women without BV almost exclusively have high presence of one of two types of *Lactobacillus* [18]. Hence, in samples with BV (high Nugent score), there are several distinct edges that have an elevated mass, which is enough to change the geometric mean, while in samples without BV (low Nugent score), most of the mass is concentrated on a single edge of the *Lactobacillus* clade, which is not enough to significantly change the geometric mean. In consequence, the denominator of the balance at the spurious edges is consistently larger for samples with BV compared to those without BV. Thus, the balance is smaller for samples with a high Nugent score, which finally explains the observed anti-correlations. Note that despite this, there are still edges that exhibit positive correlation (blue and green), which is where the actual patterns in the data outweigh the insensitivity of the geometric mean.

**S14 Fig.**
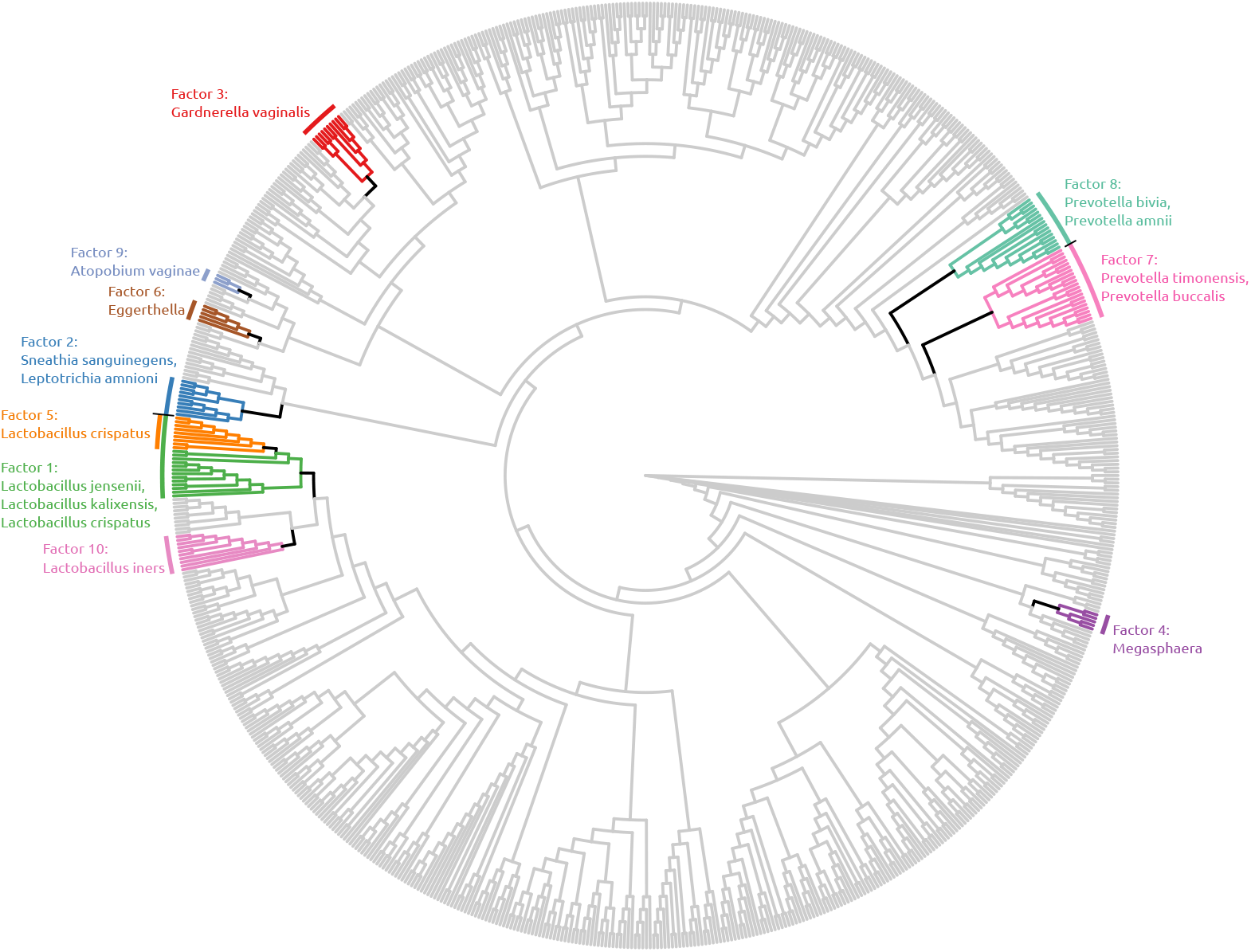
Visualization of the first ten factors of the BV dataset. Here, we show the first ten factors found by Placement-Factorization without taxon weighting on the BV dataset. The black edges are the winning edges of each iteration, which split the tree into several subtrees. For simplicity, we only colored the clades leading away from the (arbitrarily placed) root, while leaving the paraphyletic “remainder” clade in gray. Note that factor 5 is nested in factor 1, that is, it further splits the branches within the first factor, thus separating *Lactobacillus crispatus* from *Lactobacillus jensenii* and *Lactobacillus kalixensis*. See S3 Table for a comparison of the clades separated by each factor to the factors found by the original Phylofactorization. Furthermore, see S16(b) Fig for an ordination of the first two factors, showing how these factors separate the samples in the dataset. The clades found here are consistent with the findings of the original study of the dataset [18]: Healthy women without BV exhibit high abundances of *Lactobacillus*, while women affected by BV have a more diverse vaginal microbiome, containing multiple different bacterial taxa. Hence, the first factor represents the most prominent split of the data into healthy vs. diseased, based on the presence of *Lactobacillus*. Differences within the healthy samples are then further distinguished in factors five and ten, which further split parts of the *Lactobacillus* clade. The remaining factors split away clades that further separate the diseased samples from each other, based on several bacterial taxa. All clades that are found by these factors where shown in the original study to be associated with BV [18], meaning that Placement-Factorization on this dataset yields results that are consistent with previous analyses.

**S15 Fig.**
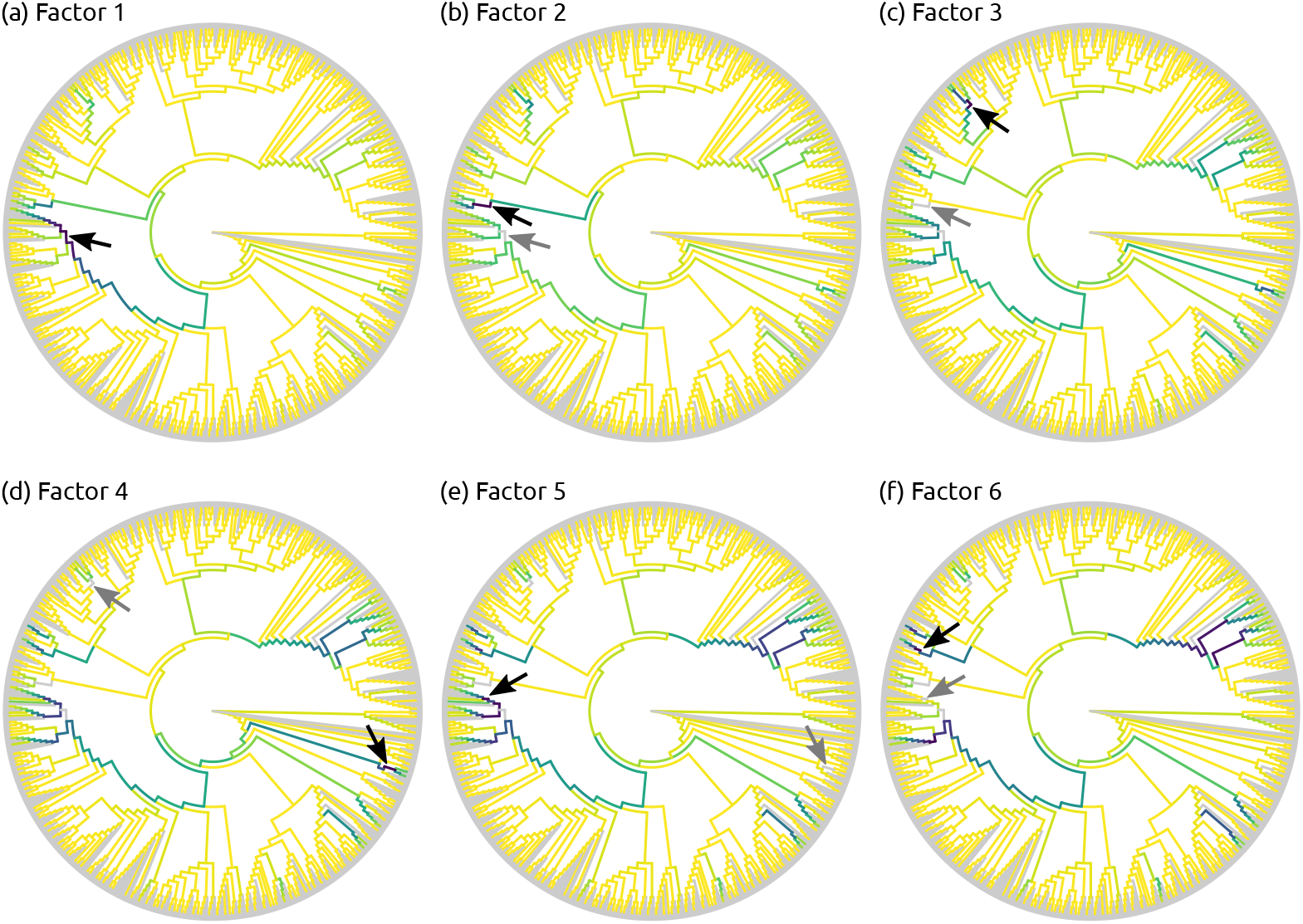
Objective function values for the first six factors of the BV dataset. In each iteration of Phylofactorization and Placement-Factorization, the objective function is evaluated for each edge of the tree (except for edges that were winning previous iterations). The figure visualizes the value of the objective function for the first six iterations of Placement-Factorization of the BV dataset, without taxon weighting. Darker edges represent higher values; the highest value of each iteration (the winning edge) is marked with a black arrow. Gray arrows further mark the winning edge of the respective previous iteration, which allows to examine the effect of “factoring out” an edge: Due to the nature of comparing the two sides induced by an edge, high values of the objective function usually propagate across several connected edges, e. g., the region of dark branches around the marked edge in (a). Once a factor has been split from the tree, the values for the whole path drop, which can be seen by comparing (a) and (b), where the region around the gray arrow has much lower values of the objective function. This behavior can consistently be observed in the other subfigures as well. The clades of the winning edges shown here can also be seen in S14 Fig, and are listed in S3 Table. See also Fig 10 for the according visualization with taxon weights.

**S16 Fig.**
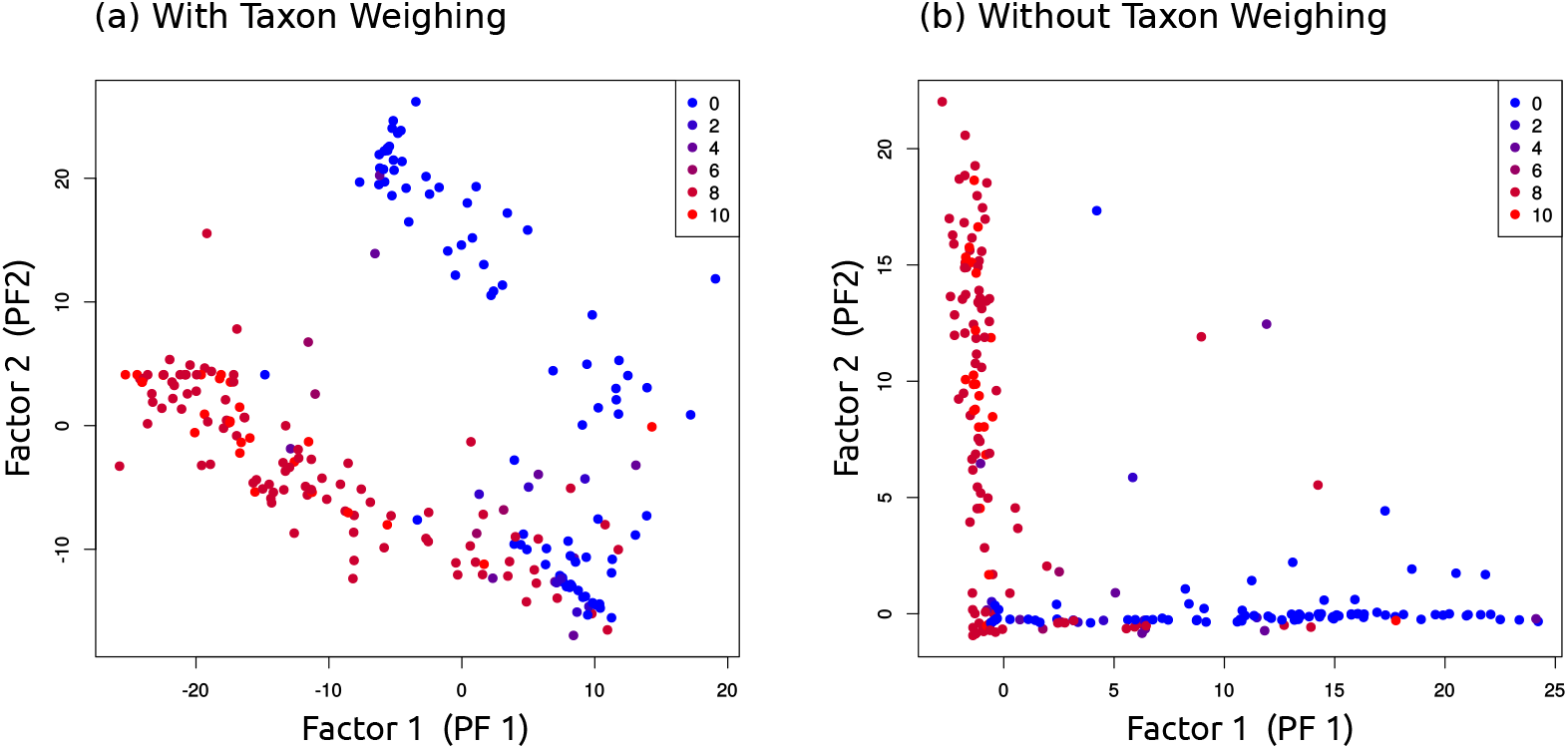
Ordination of the first two factors of the BV dataset. The figure shows ordination-visualization plots of the ILR coordinates of the first two factors found by Placement-Factorization of the BV dataset, (a) with and (b) without taxon weighting. That is, the axes correspond to the splits induced by the first two factors, while values along the axes are the balances of each sample calculated on the sets of edges of each split. Samples are again colored by their Nugent score, with 0 being the healthy patients, and 10 being the patients with severe BV. See Fig 10 for the (winning) edges that correspond to the axes in (a) (with taxon weighting), and see S14 Fig and S3 Table for the edges corresponding to the axes in (b) (without taxon weighting). This type of plot was suggested in [31] as an additional way of depicting how the factors separate samples according to meta-data features, see their Figure 5(a). Note that such visualizations can only reasonably visualize the first two or three factors, which is why they are now discontinued in the original Phylofactorization (pers. comm. with A. Washburne on 2019-01-16). For the BV dataset however, this suffices to show the major features. We also developed a way of visualizing further factors, as for example shown in S18 Fig, which alleviates this issue. On the one hand, Placement-Factorization with taxon weighting in (a) is highly similar to PCA on the balances as shown in S11(a) Fig, despite the fact that PCA does not take the meta-data into account. We suspect that is is due to the nature of the dataset, were the abundances in the *Lactobacillus* clade almost solely dictate the health status of each individual, and roughly half the samples belong to either the healthy or the sick group of patients. Hence, the *Lactobacillus* clade naturally is a major driver of differences between samples, and is thus identified by PCA as the most important component/axis. On the other hand, Placement-Factorization without taxon weighting in (b) yields an ordination that separates healthy from sick patients in the first factor, and further splits the sick patients in the second factor. The reason for this can be seen in the clades that each factor splits away from the tree, as shown in S14 Fig: The first factor separates part of the *Lactobacillus* clades, which explains why it distinguishes samples based on heath status. The second factor separates a clade containing *Sneathia sanguinegens* and *Leptotrichia amnioni*, which is an important clade among several clades that are associated with BV [18]. This can also be seen in S1 Fig, where the healthy patients with low Nugent score almost exclusively exhibit high abundances of *Lactobacillus*, while the diseased patients with high Nugent score show abundances in several clades all over the tree. This serves as a caveat for Phylofactorization, and as an example of the limitations of the type of plot shown here: It is crucial to compute all significant factors—otherwise, important aspects of the data get lost and results are incomplete. In the case of the BV dataset, two axes/factors are sufficient to separate the samples by Nugent score, but they do not explain the multitude of clades that are associated with BV.

**S17 Fig.**
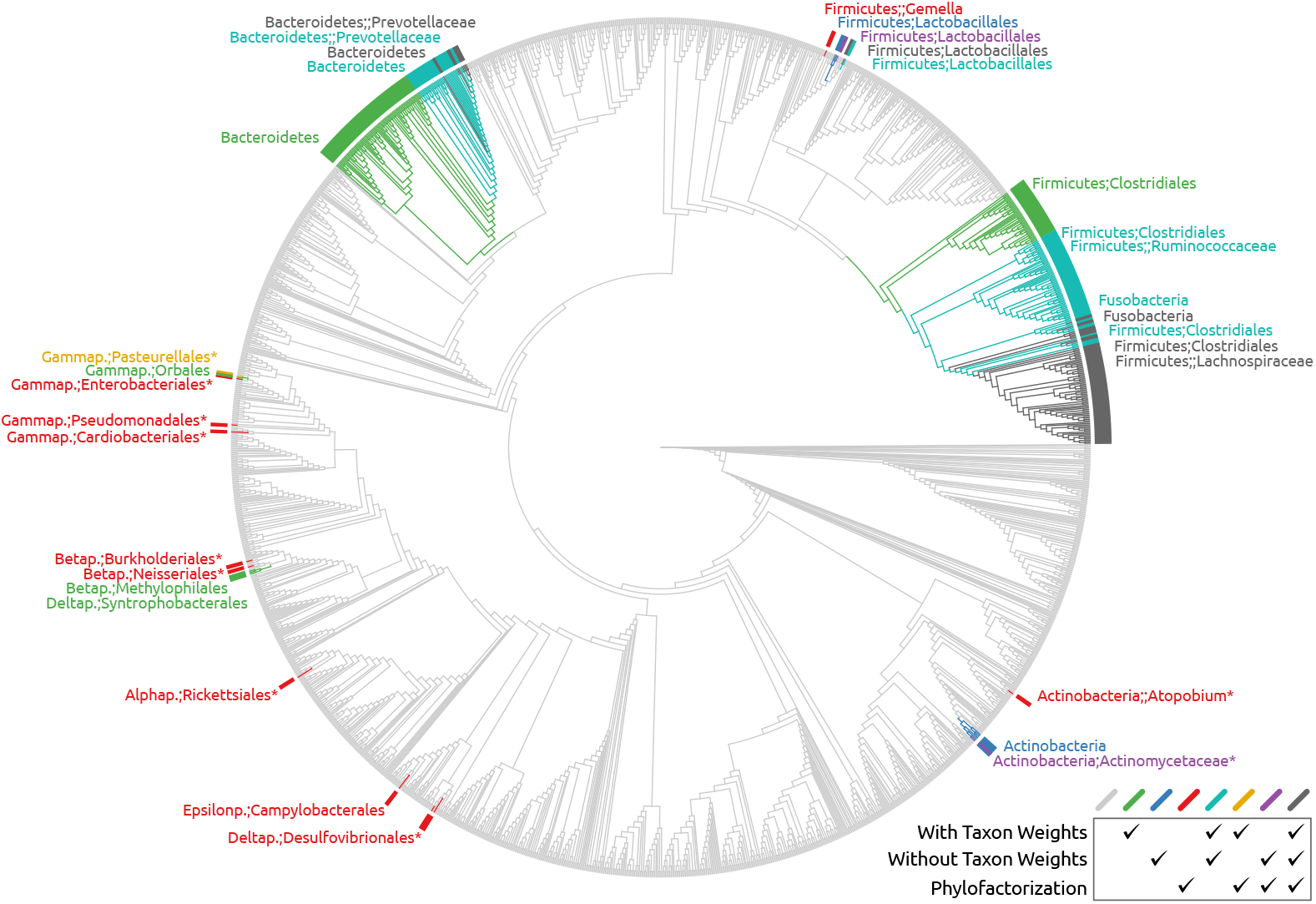
Comparison of factors found in the oral/fecal subset of the HMP dataset. Here, we compare the first 10 factors found by Placement-Factorization with and without taxon weighting on an oral/fecal subset of the HMP dataset to the first 10 factors found by Phylofactorization on their oral/fecal test dataset [31,104]. To this end, we mapped the taxa of the latter to the Silva taxonomy [105,106] that was used for constructing the reference tree shown here [34]. The clades of the tree are colored so that green, blue, and red mark branches that only appear in one of the variants, cyan, yellow, and purple for branches that occurred in two variants, and dark gray for branches that were found by all three variants. For simplicity, we here neglect the order and nesting of factors. That is, if a branch is part of the non-root side of one of the first ten factors, it is colorized here. It is striking that the variant with taxon weights (green) finds larger clades than the other two. We already observed a similar behavior with the BV dataset. However, the values of the objective function indicate that the focus of the factor is in fact much smaller and more in agreement with the other two variants shown here. Again, this behavior is similar to the BV data with taxon weighting; see Fig 10 for details. Furthermore, the several small clades and single branches found by Phylofactorization (red) that are part of the *Actinobacteria* as well as the *Alpha-, Beta-, Gamma*-, and *Deltaproteobacteria* are actually all part of the first factor. They are marked with asterisks (*) here. Due to their OTU tree only having few *Proteobacteria*, these were monophyletic in their tree. They are polyphyletic here, as our tree has more reference taxa from that group. Most of the remaining factors found by Phylofactorization are part of the gray branches of the two large clades in the figure, which are the clades that were found by all three variants. Similarly, our variants found many nested factors (factors that further split a factor of a previous iteration): The two large clades are in fact split into seven nested clades by both variants, with the remaining three factors spread across the rest of the tree (e. g., the green and blue branches). In particular the *Prevotellaceae* and parts of the *Firmicutes* were described in [31] as important clades for the distinction between oral and fecal samples, all of which were found by all three variants here. In total, despite the mismatching trees, most of the found clades agree in all three variants, however with differences in the size of the clades. More importantly, despite their differences, all variants produce factors that are well suited for separating oral from fecal samples, as shown in S18 Fig.

**S18 Fig.**
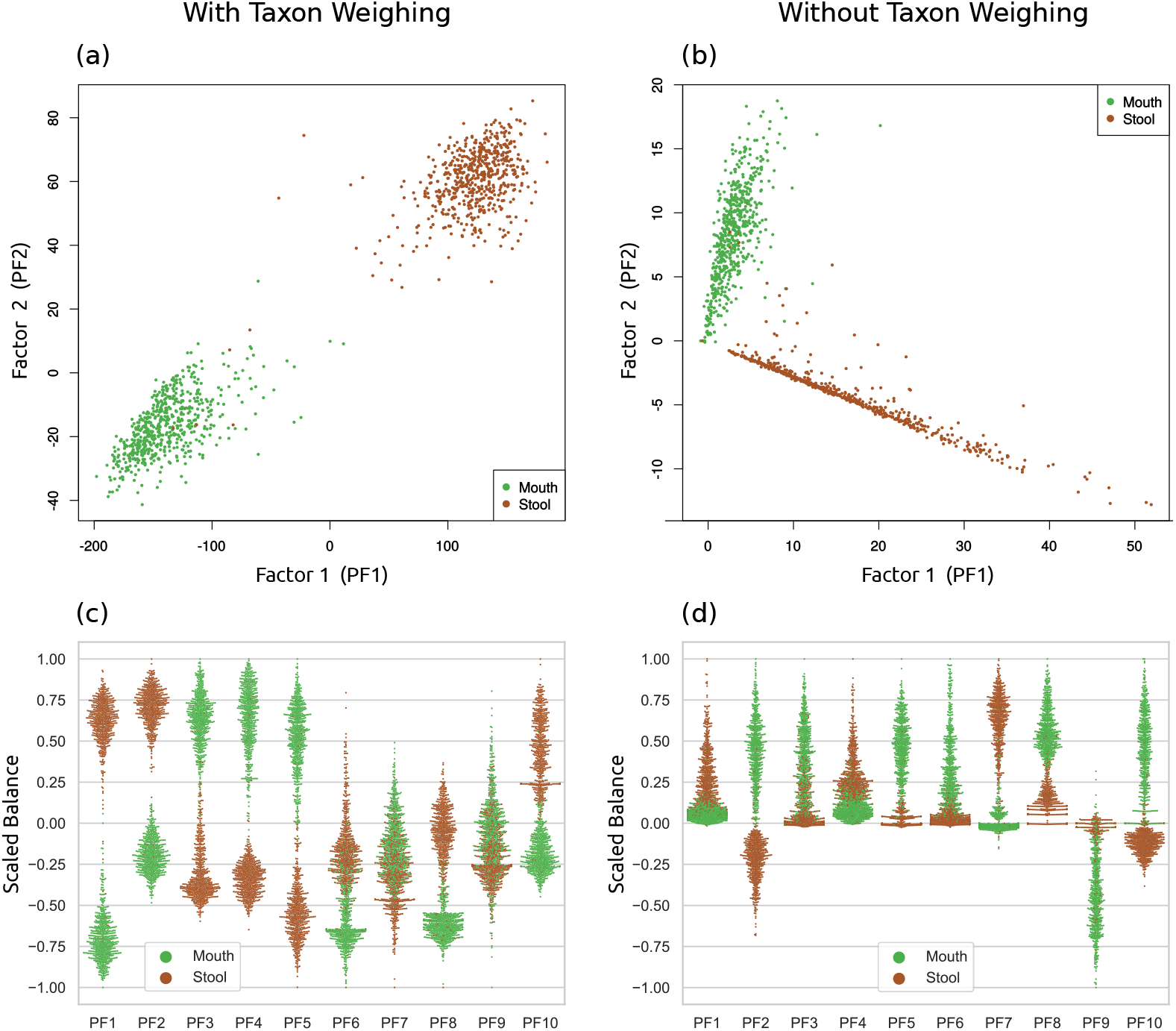
Ordination of an oral/fecal subset of the HMP dataset. The figure shows the ordination-visualization of factors found by Placement-Factorization on our oral/fecal subset of the HMP dataset. In (a) and (b), we show the balances at the winning edges of the first two factors, colorized by the body site of each sample, with and without taxon weighting. In particular, (a) exhibits a clear separation of the two body sites, similar to Figure S3 of [31]. Moreover, in order to examine how well further factors of later iterations split the data, we here employ a visualization of phylofactors, which we call *balance swarm plots*, by plotting the balances of each factor individually. This type of per-factor visualization is similar to, e. g., Figure 4 of [92]. Subfigures (c) and (d) show the first ten factors (PF1–PF10), again with and without taxon weighting, respectively. These can be understood as multi-dimensional scatter plots, where each dimension is shown separately: Each column corresponds to a factor, with the vertical axis being the balances, and horizontal space within each column used to spread samples at nearby positions, revealing their distribution density. That is, the first two columns of (c) and (d) correspond to the scatter plots of (a) and (b), respectively. The balances were scaled to bring them into the [−1.0,1.0] interval for better comparability across factors, while keeping the centering at 0. Subfigure (c) is identical to Fig 11(a) of the main text, and shows that almost all factors individually suffice to separate the data by body site; in (d), the separation is still present, but not as distinguished.

**S19 Fig.**
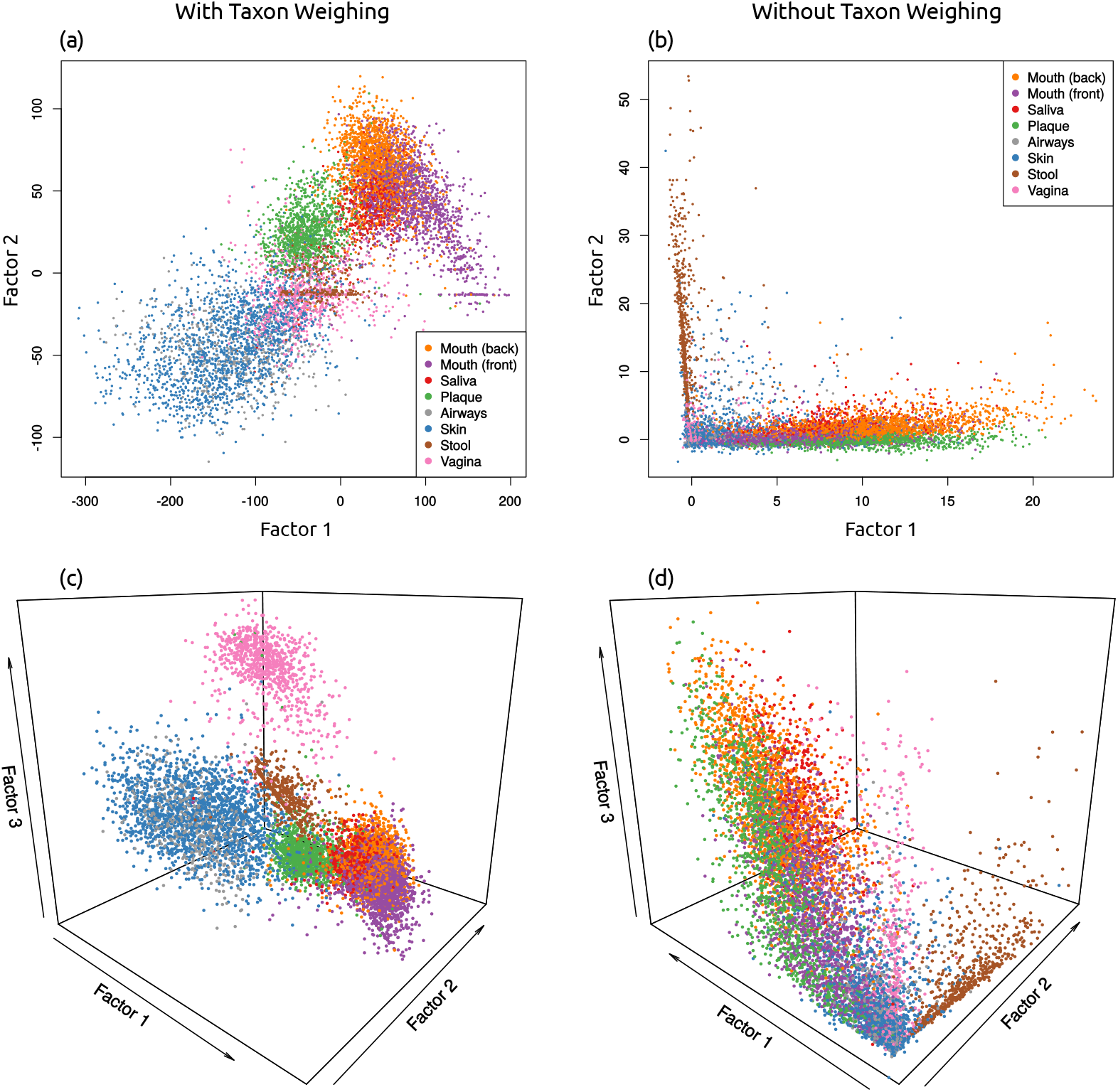
Ordination of Placement-Factorization of the full HMP dataset. In S18 Fig, we use an oral/fecal subset of the HMP dataset [16,17] to compare Placement-Factorization to a case study of the original Phylofactorization. Here, we instead used the whole HMP dataset, labeled by 8 body site regions as listed in S1 Table, to asses how well Placement-Factorization with a GLM objective function can separate samples based on their body site label. Again, we show the balances of the winning edges of the first two and three factors, respectively, with and without taxon weighting. As in S16 Fig and S18 Fig before, the plots with taxon weighting form “clouds”, whereas the plots without form an “L”-shape. In all cases, a separation of the oral samples from the other regions is clearly visible. Noticeably, in (a), a part of the stool and mouth samples form a horizontal line, which indicates that the second factor does not distinguish between samples from those regions. It is striking that the samples from the vaginal region in (a) and (c) are not separated from the other samples until the third factor, which is visible as a pink cloud above the rest of the samples in (c). This again serves as a caveat that one needs to consider enough factors in order to get a complete understanding of the results. To this end, we furthermore show a more detailed version of these scatter plots for multiple factors/dimensions in S20 Fig.

**S20 Fig.**
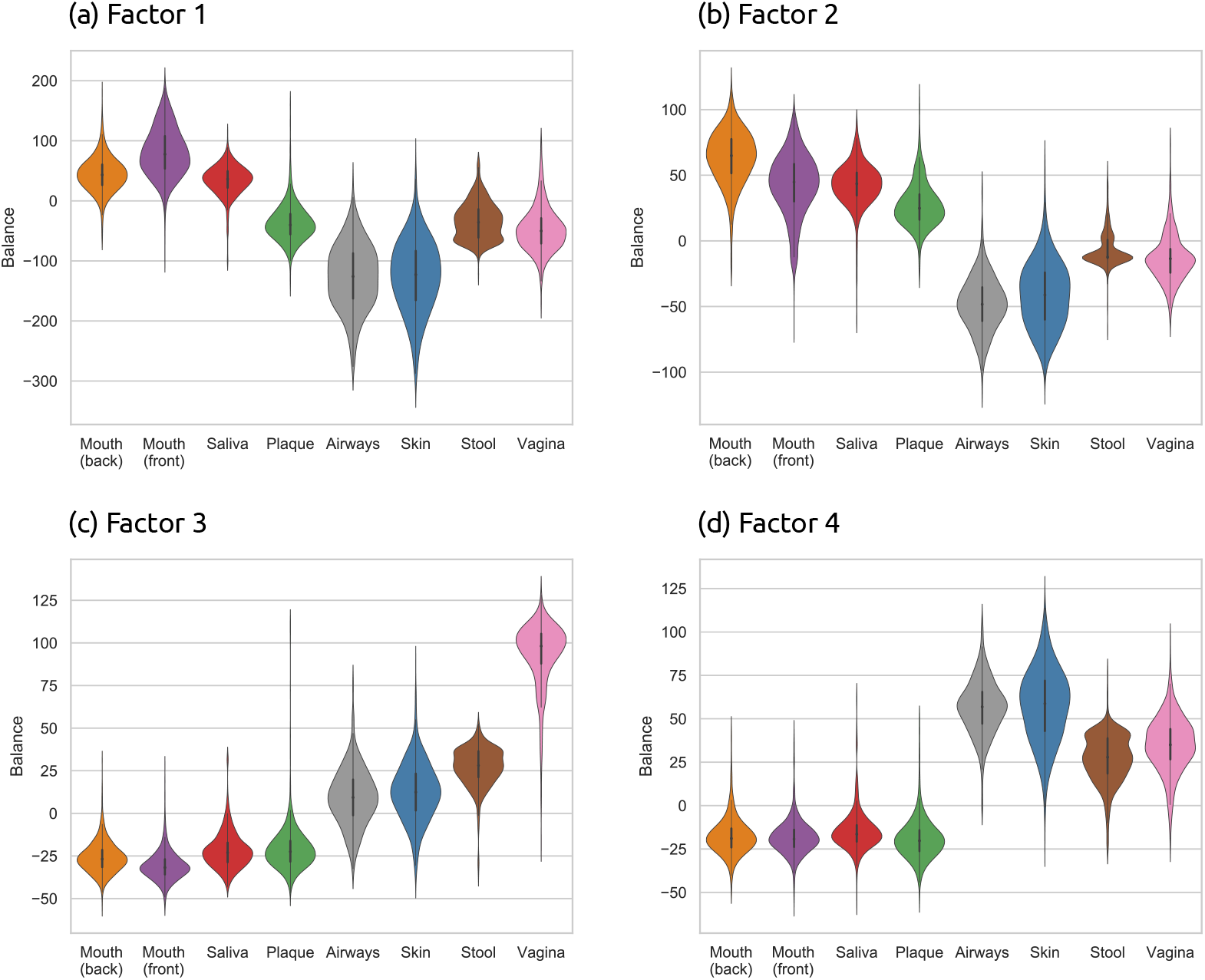
Ordination of the first four factors of the HMP dataset. The *balance swarm plots* as shown in S18(c) Fig and S18(d) Fig allow for a more detailed understanding of how each factor separates the samples. They can be colored by either continuous meta-data variables, similar to Figure 5(a) of [31], or a categorical variable with a limited number of categories, as shown in S18 Fig. However, for the eight body regions that we use for the HMP data, this type of visualization becomes hard to inspect visually. Hence, we here extend on the idea of balance swarm plots, and show the distribution of balances for each factor and for each body sites separately. That is, each subfigure here shows the balances of the winning edge of a factor, grouped by the categorical meta-data variable body site. In other words, each subfigure here represents a disentangled column of a balance swarm plot, where each body site is displayed by its own violin. The data shown here is again the result of Placement-Factorization with taxon weighting on the full HMP dataset, as shown and explained in S19 Fig. See there for the scatter plots that correspond to the first two and three factors shown here. Subfigure (a) is identical to Fig 11(b) of the main text, and included here for comparability. In all subfigures, the oral regions are separated from the other regions: For example, in (a), mouth and saliva samples exhibit balances above 0, in contrast to all other samples. In (b), (c), and (d), the plaque samples join the other oral samples in terms of their balance values. In (b), the stool samples have a distinct bulge near 0, which corresponds to the horizontal line (factor 2) in S19(a) Fig. Furthermore, the third factor in (c) again clearly separates the vaginal samples from the rest, as shown before in S19(c) Fig.

## References

1. Escobar-Zepeda A, Vera-Ponce De León A, Sanchez-Flores A. The road to metagenomics: From microbiology to DNA sequencing technologies and bioinformatics. Frontiers in Genetics. 2015;6(348):1–15. doi:10.3389/fgene.2015.00348.

2. Logares R, Haverkamp THA, Kumar S, Lanzén A, Nederbragt AJ, Quince C, et al. Environmental microbiology through the lens of high-throughput DNA sequencing: Synopsis of current platforms and bioinformatics approaches. Journal of Microbiological Methods. 2012;91(1):106–113. doi:10.1016/J.MIMET.2012.07.017.

3. Pareek CS, Smoczynski R, Tretyn A. Sequencing technologies and genome sequencing. Journal of Applied Genetics. 2011;52(4):413–435. doi:10.1007/s13353-011-0057-x.

4. Niedringhaus TP, Milanova D, Kerby MB, Snyder MP, Barron AE. Landscape of Next-Generation Sequencing Technologies. Analytical Chemistry. 2011;83(12):4327–4341. doi:10.1021/ac2010857.

5. Mignardi M, Nilsson M. Fourth-generation sequencing in the cell and the clinic. Genome Medicine. 2014;6(4):31. doi:10.1186/gm548.

6. Heather JM, Chain B. The sequence of sequencers: The history of sequencing DNA. Genomics. 2016;107(1):1–8. doi:10.1016/J.YGENO.2015.11.003.

7. Morgan JL, Darling AE, Eisen JA. Metagenomic sequencing of an in vitro-simulated microbial community. PLoS ONE. 2010;5(4):1–10. doi:10.1371/journal.pone.0010209.

8. Edwards DJ, Holt KE. Beginner’s guide to comparative bacterial genome analysis using next-generation sequence data. Microbial informatics and experimentation. 2013;3(1):2. doi:10.1186/2042-5783-3-2.

9. Sunagawa S, Mende DR, Zeller G, Izquierdo-Carrasco F, Berger Sa, Kultima JR, et al. Metagenomic species profiling using universal phylogenetic marker genes. Nature Methods. 2013;10(12):1196. doi:10.1038/nmeth.2693.

10. Matsen IV FA. Phylogenetics and the Human Microbiome. Systematic Biology. 2015;64(1):e26–e41. doi:10.1093/sysbio/syu053.

11. Karsenti E, Acinas SG, Bork P, Bowler C, de Vargas C, Raes J, et al. A holistic approach to marine Eco-systems biology. PLoS Biology. 2011;9(10):7–11. doi:10.1371/journal.pbio.1001177.

12. Giner CR, Forn I, Romac S, Logares R, Vargas CD. Environmental Sequencing Provides Reasonable Estimates of the Relative Abundance of Specific Picoeukaryotes. Applied and Environmental Microbiology. 2016;82(15):4757–4766. doi:10.1128/AEM.00560-16.Address.

13. Gran-Stadniczeñko S, Šupraha L, Egge ED, Edvardsen B. Haptophyte Diversity and Vertical Distribution Explored by 18S and 28S Ribosomal RNA Gene Metabarcoding and Scanning Electron Microscopy. Journal of Eukaryotic Microbiology. 2017; p. 1–19. doi:10.1111/jeu.12388.

14. Dupont AÖC, Griffiths RI, Bell T, Bass D. Differences in soil micro-eukaryotic communities over soil pH gradients are strongly driven by parasites and saprotrophs. Environmental Microbiology. 2016;18(6):2010–2024. doi:10.1111/1462-2920.13220.

15. Mahé F, de Vargas C, Bass D, Czech L, Stamatakis A, Lara E, et al. Parasites dominate hyperdiverse soil protist communities in Neotropical rainforests. Nature Ecology & Evolution. 2017;1(4):0091. doi:10.1038/s41559-017-0091.

16. Huttenhower C, Gevers D, Knight R, Abubucker S, Badger JH, Chinwalla AT, et al. Structure, function and diversity of the healthy human microbiome. Nature. 2012;486(7402):207–214. doi:10.1038/nature11234.

17. Methé BA, Nelson KE, Pop M, Creasy HH, Giglio MG, Huttenhower C, et al. A framework for human microbiome research. Nature. 2012;486(7402):215–221. doi:10.1038/nature11209.A.

18. Srinivasan S, Hoffman NG, Morgan MT, Matsen FA, Fiedler TL, Hall RW, et al. Bacterial communities in women with bacterial vaginosis: High resolution phylogenetic analyses reveal relationships of microbiota to clinical criteria. PLOS ONE. 2012;7(6):e37818. doi:10.1371/journal.pone.0037818.

19. Altschul SF, Gish W, Miller W, Myers EW, Lipman DJ. Basic Local Alignment Search Tool. Journal of Molecular Biology. 1990;215(3):403–410. doi:10.1016/S0022-2836(05)80360-2.

20. Shah N, Nute MG, Warnow T, Pop M. Misunderstood parameter of NCBI BLAST impacts the correctness of bioinformatics workflows. Bioinformatics. 2018;doi:10.1093/bioinformatics/bty833.

21. Koski LB, Golding GB. The closest BLAST hit is often not the nearest neighbor. Journal of molecular evolution. 2001;52(6):540–2. doi:10.1007/s002390010184.

22. Matsen FA, Kodner RB, Armbrust EV. pplacer: linear time maximum-likelihood and Bayesian phylogenetic placement of sequences onto a fixed reference tree. BMC Bioinformatics. 2010;11(1):538. doi:10.1186/1471-2105-11-538.

23. Berger S, Krompass D, Stamatakis A. Performance, accuracy, and web server for evolutionary placement of short sequence reads under maximum likelihood. Systematic Biology. 2011;60(3):291–302. doi:10.1093/sysbio/syr010.

24. Barbera P, Kozlov AM, Czech L, Morel B, Darriba D, Flouri T, et al. EPA-ng: Massively Parallel Evolutionary Placement of Genetic Sequences. Systematic Biology. 2018;doi:10.1093/sysbio/syy054.

25. Pace NR. A molecular view of microbial diversity and the biosphere. Science. 1997;276(5313):734–740.

26. Hugenholtz P, Goebel BM, Pace NR. Impact of Culture-Independent Studies on the Emerging Phylogenetic View of Bacterial Diversity. Journal of Bacteriology. 1998;180(18):4765–4774.

27. Nguyen Np, Mirarab S, Liu B, Pop M, Warnow T. TIPP: taxonomic identification and phylogenetic profiling. Bioinformatics. 2014;30(24):3548–3555. doi:10.1093/bioinformatics/btu721.

28. Kozlov AM, Zhang J, Yilmaz P, Glöckner FO, Stamatakis A. Phylogeny-aware identification and correction of taxonomically mislabeled sequences. Nucleic Acids Research. 2016;44(11):5022–5033. doi:10.1093/nar/gkw396.

29. Matsen FA, Evans SN. Edge principal components and squash clustering: using the special structure of phylogenetic placement data for sample comparison. PLOS ONE. 2011;8(3):1–17. doi:10.1371/journal.pone.0056859.

30. Silverman JD, Washburne AD, Mukherjee S, David LA. A phylogenetic transform enhances analysis of compositional microbiota data. eLife. 2017;6:e21887. doi:10.7554/eLife.21887.

31. Washburne AD, Silverman JD, Leff JW, Bennett DJ, Darcy JL, Mukherjee S, et al. Phylogenetic factorization of compositional data yields lineage-level associations in microbiome datasets. PeerJ. 2017;5:e2969. doi:10.7717/peerj.2969.

32. Sunagawa S, Coelho LP, Chaffron S, Kultima JR, Labadie K, Salazar G, et al. Structure and function of the global ocean microbiome. Science. 2015;348(6237):1–10. doi:10.1126/science.1261359.

33. Guidi L, Chaffron S, Bittner L, Eveillard D, Larhlimi A, Roux S, et al. Plankton networks driving carbon export in the oligotrophic ocean. Nature. 2016;532(7600):465–470. doi:10.1038/nature16942.

34. Czech L, Barbera P, Stamatakis A. Methods for Automatic Reference Trees and Multilevel Phylogenetic Placement. Bioinformatics. 2018; p. 299792. doi:10.1093/bioinformatics/bty767.

35. Berger S, Stamatakis A. Aligning short reads to reference alignments and trees. Bioinformatics. 2011;27(15):2068–2075. doi:10.1093/bioinformatics/btr320.

36. Berger S, Stamatakis A. PaPaRa 2.0: A Vectorized Algorithm for Probabilistic Phylogeny-Aware Alignment Extension. Heidelberg: Heidelberg Institute for Theoretical Studies; 2012.

37. Eddy SR. Profile hidden Markov models. Bioinformatics. 1998;14(9):755–763.

38. Eddy SR. A new generation of homology search tools based on probabilistic inference. In: Genome Informatics. vol. 23. World Scientific; 2009. p. 205–211.

39. Tavaré S. Some probabilistic and statistical problems in the analysis of DNA sequences. American Mathematical Society: Lectures on Mathematics in the Life Sciences. 1986;17:57–86. doi:citeulike-article-id:4801403.

40. Strimmer K, Rambaut A. Inferring confidence sets of possibly misspecified gene trees. Proceedings of the Royal Society of London B: Biological Sciences. 2002;269(1487):137–142.

41. von Mering C, Hugenholtz P, Raes J, Tringe SG, Doerks T, Jensen LJ, et al. Quantitative Phylogenetic Assessment of Microbial Communities in Diverse Environments. Science. 2007;315(5815):1126–1130. doi:10.1126/science.1133420.

42. Gloor GB, Macklaim JM, Pawlowsky-Glahn V, Egozcue JJ. Microbiome Datasets Are Compositional: And This Is Not Optional. Frontiers in Microbiology. 2017;8:2224. doi:10.3389/fmicb.2017.02224.

43. Aitchison J. The statistical analysis of compositional data. Chapman and Hall London; 1986.

44. Jackson DA. Compositional data in community ecology: The paradigm or peril of proportions? Ecology. 1997;78(3):929–940. doi:10.1890/0012-9658(1997)078[0929:CDICET]2.0.CO;2.

45. Tsilimigras MCB, Fodor AA. Compositional data analysis of the microbiome: fundamentals, tools, and challenges. Annals of Epidemiology. 2016;26(5):330–335. doi:10.1016/j.annepidem.2016.03.002.

46. Gloor GB, Macklaim JM, Vu M, Fernandes AD. Compositional uncertainty should not be ignored in high-throughput sequencing data analysis. Austrian Journal of Statistics. 2016;45(4):73. doi:10.17713/ajs.v45i4.122.

47. Weiss S, Xu ZZ, Peddada S, Amir A, Bittinger K, Gonzalez A, et al. Normalization and microbial differential abundance strategies depend upon data characteristics. Microbiome. 2017;5(1):27. doi:10.1186/s40168-017-0237-y.

48. Gotelli NJ, Colwell RK. Quantifying biodiversity: procedures and pitfalls in the measurement and comparison of species richness. Ecology Letters. 2001;4(4):379–391. doi:10.1046/j.1461-0248.2001.00230.x.

49. McMurdie PJ, Holmes S. Waste Not, Want Not: Why Rarefying Microbiome Data Is Inadmissible. PLoS Computational Biology. 2014;10(4):e1003531. doi:10.1371/journal.pcbi.1003531.

50. Logares R, Sunagawa S, Salazar G, Cornejo-Castillo FM, Ferrera I, Sarmento H, et al. Metagenomic 16S rDNA Illumina tags are a powerful alternative to amplicon sequencing to explore diversity and structure of microbial communities. Environmental Microbiology. 2014;16(9):2659–2671. doi:10.1111/1462-2920.12250.

51. Edgar RC. Search and clustering orders of magnitude faster than BLAST. Bioinformatics. 2010;26(19):2460–2461. doi:10.1093/bioinformatics/btq461.

52. Mahé F, Rognes T, Quince C, de Vargas C, Dunthorn M. Swarm: Robust and fast clustering method for amplicon-based studies. PeerJ. 2014;2:1–12. doi:http://dx.doi.org/10.7287/peerj.preprints.386v1.

53. Mahé F, Rognes T, Quince C, De Vargas C, Dunthorn M. Swarm v2: Highly-scalable and high-resolution amplicon clustering. PeerJ. 2015;.

54. Rognes T, Flouri T, Nichols B, Quince C, Mahé F. VSEARCH: a versatile open source tool for metagenomics. PeerJ. 2016;4:e2584.

55. Gloor GB, Wu JR, Pawlowsky-Glahn V, Egozcue JJ. It’s all relative: analyzing microbiome data as compositions. Annals of epidemiology. 2016;26(5):322–9. doi:10.1016/j.annepidem.2016.03.003.

56. Evans SN, Matsen FA. The phylogenetic Kantorovich-Rubinstein metric for environmental sequence samples. Journal of the Royal Statistical Society Series B: Statistical Methodology. 2012;74:569–592. doi:10.1111/j.1467-9868.2011.01018.x.

57. Lozupone C, Knight R. UniFrac: a New Phylogenetic Method for Comparing Microbial Communities. Applied and Environmental Microbiology. 2005;71(12):8228–8235. doi:10.1128/AEM.71.12.8228.

58. Lozupone CA, Hamady M, Kelley ST, Knight R. Quantitative and Qualitative *β* Diversity Measures Lead to Different Insights into Factors That Structure Microbial Communities. Applied and Environmental Microbiology. 2007;73(5):1576–1585. doi:10.1128/AEM.01996-06.

59. Lovell D, Pawlowsky-Glahn V, Egozcue JJ, Marguerat S, Bähler J. Proportionality: A Valid Alternative to Correlation for Relative Data. PLOS Computational Biology. 2015;11(3):e1004075. doi:10.1371/journal.pcbi.1004075.

60. Dunthorn M, Otto J, Berger SA, Stamatakis A, Mahé F, Romac S, et al. Placing environmental next-generation sequencing amplicons from microbial eukaryotes into a phylogenetic context. Molecular Biology and Evolution. 2014;31(4):993–1009. doi:10.1093/molbev/msu055.

61. Letunic I, Bork P. Interactive tree of life (iTOL) v3: an online tool for the display and annotation of phylogenetic and other trees. Nucleic acids research. 2016;44(W1):W242–5. doi:10.1093/nar/gkw290.

62. Stamatakis A. RAxML version 8: a tool for phylogenetic analysis and post-analysis of large phylogenies. Bioinformatics. 2014;30(9):1312–1313. doi:10.1093/bioinformatics/btu033.

63. Yu G, Smith DK, Zhu H, Guan Y, Lam TTY. ggtree: an r package for visualization and annotation of phylogenetic trees with their covariates and other associated data. Methods in Ecology and Evolution. 2017;8(1):28–36. doi:10.1111/2041-210X.12628.

64. Everitt BS, Skrondal A. The Cambridge Dictionary of Statistics. 4th ed. Cambridge University Press; 2010.

65. Mallows CL. A Note on Asymptotic Joint Normality. Ann Math Statist. 1972;43(2):508–515. doi:10.1214/aoms/1177692631.

66. Rachev ST. The Monge-Kantorovich Mass Transference Problem and its Stochastic Applications. Theory of Probability and its Applications. 1985;29(4):647–676.

67. Levina E, Bickel P. The earth mover’s distance is the Mallows distance: some insights from statistics. Eighth IEEE International Conference on Computer Vision. 2001; p. 251–256. doi:10.1109/ICCV.2001.937632.

68. Villani C. Optimal transport: old and new. Springer Science & Business Media; 2008.

69. Michener CD, Sokal RR. A quantitative approach to a problem in classification. Evolution. 1957;11(2):130–162.

70. Sokal RR. A statistical method for evaluating systematic relationship. University of Kansas science bulletin. 1958;28:1409–1438.

71. Legendre P, Legendre LFJ. Numerical Ecology. Developments in Environmental Modelling. Elsevier Science; 1998.

72. MacQueen J. Some methods for classification and analysis of multivariate observations. Proceedings of the Fifth Berkeley Symposium on Mathematical Statistics and Probability. 1967;1(233):281–297. doi:citeulike-article-id:6083430.

73. Kelley DR, Salzberg SL. Clustering metagenomic sequences with interpolated Markov models. BMC Bioinformatics. 2010;11(1):544. doi:10.1186/1471-2105-11-544.

74. Lloyd SP. Least Squares Quantization in PCM. IEEE Transactions on Information Theory. 1982;28(2):129–137. doi:10.1109/TIT.1982.1056489.

75. Arthur D, Vassilvitskii S. k-means++: The Advantages of Careful Seeding. In: Proceedings of the eighteenth annual ACM-SIAM symposium on Discrete algorithms. Society for Industrial and Applied Mathematics Philadelphia, PA, USA; 2007. p. 1027–1035.

76. Kanungo T, Mount DM, Netanyahu NS, Wu AY, Piatko CD, Silverman R, et al. A Local Search Approximation Algorithm for k-Means Clustering. Computational Geometry. 2003;28(2-3):89–112.

77. Bottou L, Bengio Y. Convergence properties of the k-means algorithms. In: Advances in neural information processing systems; 1995. p. 585–592.

78. Arthur D, Vassilvitskii S. How Slow is the K-means Method? In: Proceedings of the Twenty-second Annual Symposium on Computational Geometry. SCG ‘06. New York, NY, USA: ACM; 2006. p. 144–153.

79. Thorndike RL. Who belongs in the family? Psychometrika. 1953;18(4):267–276.

80. Rousseeuw PJ. Silhouettes: A graphical aid to the interpretation and validation of cluster analysis. Journal of Computational and Applied Mathematics. 1987;20:53–65.

81. Bischof H, Leonardis A, Selb A. MDL Principle for Robust Vector Quantisation. Pattern Analysis & Applications. 1999;2(1):59–72. doi:10.1007/s100440050015.

82. Pelleg D, Moore AW, Others. X-means: Extending K-means with Efficient Estimation of the Number of Clusters. In: ICML. vol. 1; 2000. p. 727–734.

83. Tibshirani R, Walther G, Hastie T. Estimating the number of clusters in a data set via the gap statistic. Journal of the Royal Statistical Society: Series B (Statistical Methodology). 2001;63(2):411–423.

84. Hamerly G, Elkan C. Learning the k in k-means. In: Thrun S, Saul LK, Schölkopf PB, editors. Advances in Neural Information Processing Systems 16. MIT Press; 2004. p. 281–288.

85. Morton JT, Sanders J, Quinn RA, McDonald D, Gonzalez A, Vázquez-Baeza Y, et al. Balance Trees Reveal Microbial Niche Differentiation. mSystems. 2017;2(1). doi:10.1128/mSystems.00162-16.

86. Egozcue JJ, Pawlowsky-Glahn V, Mateu-Figueras G, Barceló-Vidal C. Isometric Logratio Transformations for Compositional Data Analysis. Mathematical Geology. 2003;35(3):279–300. doi:10.1023/A:1023818214614.

87. Egozcue JJ, Pawlowsky-Glahn V. Groups of Parts and Their Balances in Compositional Data Analysis. Mathematical Geology. 2005;37(7):795–828. doi:10.1007/s11004-005-7381-9.

88. Pawlowsky-Glahn V, Egozcue JJ, Tolosana-Delgado R. Modeling and Analysis of Compositional Data. Chichester, UK: John Wiley & Sons; 2015.

89. Egozcue JJ, Pawlowsky-Glahn V. Changing the Reference Measure in the Simplex and its Weighting Effects. Austrian Journal of Statistics. 2016;45(4):25. doi:10.17713/ajs.v45i4.126.

90. Good IJ. On the Estimation of Small Frequencies in Contingency Tables. Journal of the Royal Statistical Society Series B (Methodological). 1956;18(1):113–124.

91. Washburne AD, Silverman JD, Morton JT, Becker D, Crowley D, Mukherjee S, et al. Phylofactorization - a graph partitioning algorithm to identify phylogenetic scales of ecological data. bioRxiv. 2018; p. 235341. doi:10.1101/235341.

92. Washburne AD, Silverman JD, Morton JT, Becker DJ, Crowley D, Mukherjee S, et al. Phylofactorization: a graph partitioning algorithm to identify phylogenetic scales of ecological data. Ecological Monographs. 2019;(e01353). doi:10.1002/ecm.1353.

93. Nelder JA, Wedderburn RWM. Generalized Linear Models. Journal of the Royal Statistical Society Series A (General). 1972;135(3):370–384. doi:10.2307/2344614.

94. McCullagh P, Nelder JA. Generalized Linear Models. vol. 37. CRC press; 1989.

95. Agresti A. An Introduction to Categorical Data Analysis. 3rd ed. Wiley-Interscience; 2018.

96. Pawlowsky-Glahn V, Buccianti A. Compositional Data Analysis: Theory and Applications. John Wiley & Sons; 2011.

97. Nugent RP, Krohn MA, Hillier SL. Reliability of diagnosing bacterial vaginosis is improved by a standardized method of gram stain interpretation. Journal of clinical microbiology. 1991;29(2):297–301.

98. Amsel R, Totten PA, Spiegel CA, Chen KCS, Eschenbach D, Holmes KK. Nonspecific vaginitis: Diagnostic Criteria and Microbial and Epidemiologic Associations. The American Journal of Medicine. 1983;74(1):14–22. doi:10.1016/0002-9343(83)91112-9.

99. Lozupone CA, Knight R. Global patterns in bacterial diversity. Proceedings of the National Academy of Sciences. 2007;104(27):11436–11440. doi:10.1073/pnas.0611525104.

100. Potapova M. Patterns of Diatom Distribution In Relation to Salinity. Kociolek J, Seckbach JP, editors. Springer; 2011.

101. Matsen FA, Evans SN. Edge principal components and squash clustering: using the special structure of phylogenetic placement data for sample comparison. arXiv. 2011;.

102. Mardia KV. Some Properties of Classical Multi-Dimesional Scaling. Communications in Statistics-Theory and Methods. 1978;7(13):1233–1241.

103. Krzanowski WJ, Marriott F. Multivariate Analysis. Wiley; 1994.

104. Caporaso JG, Lauber CL, Costello EK, Berg-Lyons D, Gonzalez A, Stombaugh J, et al. Moving pictures of the human microbiome. Genome Biology. 2011;12(5):R50. doi:10.1186/gb-2011-12-5-r50.

105. Quast C, Pruesse E, Yilmaz P, Gerken J, Schweer T, Yarza P, et al. The SILVA ribosomal RNA gene database project: improved data processing and web-based tools. Nucleic Acids Research. 2013;41(D1):D590–D596. doi:10.1093/nar/gks1219.

106. Yilmaz P, Parfrey LW, Yarza P, Gerken J, Pruesse E, Quast C, et al. The SILVA and “All-species Living Tree Project (LTP)” taxonomic frameworks. Nucleic Acids Research. 2014;42(D1):D643–D648. doi:10.1093/nar/gkt1209.

107. Dunn JC. A Fuzzy Relative of the ISODATA Process and Its Use in Detecting Compact Well-Separated Clusters. Journal of Cybernetics. 1973;3(3):32–57. doi:10.1080/01969727308546046.

108. Bezdek JC. Pattern Recognition with Fuzzy Objective Function Algorithms. Advanced applications in pattern recognition. Plenum Press; 1981.

109. Kriegel HP, Kröger P, Sander J, Zimek A. Density-based clustering. Wiley Interdisciplinary Reviews: Data Mining and Knowledge Discovery. 2011;1(3):231–240.

110. Berger S, Czech L. PaPaRa 2.0 with MPI; 2016. Available from: https://github.com/lczech/papara_nt.

111. Barbera P. EPA-ng – Massively Parallel Phylogenetic Placement of Genetic Sequences; 2017. Online: https://github.com/Pbdas/epa-ng.

112. Katoh K, Misawa K, Kuma K, Miyata T. MAFFT: a novel method for rapid multiple sequence alignment based on fast Fourier transform. Nucleic Acids Research. 2002;30(14):3059–3066. doi:10.1093/nar/gkf436.

113. Katoh K, Standley DM. MAFFT Multiple Sequence Alignment Software Version 7: Improvements in Performance and Usability. Molecular Biology and Evolution. 2013;30(4):772–780. doi:10.1093/molbev/mst010.

114. Katoh K, Kuma Ki, Toh H, Miyata T. MAFFT version 5: improvement in accuracy of multiple sequence alignment. Nucleic Acids Research. 2005;33(2):511–518. doi:10.1093/nar/gki198.

115. Kozlov A, Darriba D, Flouri T, Morel B, Stamatakis A. RAxML-NG: A fast, scalable, and user-friendly tool for maximum likelihood phylogenetic inference. bioRxiv. 2018; p. 447110. doi:10.1101/447110.

116. Mahé F. Fred’s metabarcoding pipeline; 2016. Available from: https://github.com/frederic-mahe/swarm/wiki/Fred’s-metabarcoding-pipeline.

117. Zhang J, Kobert K, Flouri T, Stamatakis A. PEAR: A fast and accurate Illumina Paired-End reAd mergeR. Bioinformatics. 2014;30(5):614–620. doi:10.1093/bioinformatics/btt593.

118. Martin M. Cutadapt removes adapter sequences from high-throughput sequencing reads. EMBnetjournal. 2011;17(1):10. doi:10.14806/ej.17.1.200.

119. Matsen FA, Hoffman NG, Gallagher A, Stamatakis A. A format for phylogenetic placements. PLoS ONE. 2012;7(2):1–4. doi:10.1371/journal.pone.0031009.

120. Vinh NX, Epps J, Bailey J. Information Theoretic Measures for Clusterings Comparison: Variants, Properties, Normalization and Correction for Chance. Journal ofMachine Learning Research. 2010;11:2837–2854.

